# Genomic and Epigenetic Interplay Drives Adaptive Fusion via Reproduction Trade-Off

**DOI:** 10.1101/2025.09.26.678705

**Authors:** Yang Chen, Lin Bi, Xiaoli Fan, Shuang Liang, Chengyuan Li, Yihong Wang, Sophie G Martin, Gaowen Liu

**Affiliations:** Shenzhen Key Laboratory of Synthetic Genomics, Guangdong Provincial Key Laboratory of Synthetic Genomics, Key Laboratory of Quantitative Synthetic Biology, Shenzhen Institute of Synthetic Biology, Shenzhen Institute of Advanced Technology, Chinese Academy of Sciences, Shenzhen 518055, China; School of Pharmacy, Faculty of Medicine, Macau University of Science and Technology, Macau SAR, China; Faculty of health sciences, University of Macau, Avenida da Universidade, Taipa, Macau SAR, China; Dept of Molecular, Cell, and Developmental Biology, University of California, Los Angeles, USA; Dept. of Molecular and Cellular Biology, University of Geneva, 30, quai Ernest-Ansermet CH-1211, Geneva 4, Switzerland; Dept. of Fundamental Microbiology, Faculty of Biology and Medicine, University of Lausanne, Switzerland

**Keywords:** Sexual–asexual trade-off, Genetic assimilation, Network rewiring, Latent capacity, Subtelomeric silencing, *Schizosaccharomyces pombe*, Experimental evolution

## Abstract

Investment in sexual reproduction has long been seen as a fitness cost for asexual proliferation, raising the question of how cells evolve to accommodate cyclical demands of sexual and asexual cycles. We addressed this question by evolving *Schizosaccharomyces pombe* lacking the critical fusion gene Prm1 through 18 cycles of alternating meiosis and mitosis. Initially, transcriptional tweaks offered a fragile means to fuse despite Prm1’s absence. However, a telomeric 1% genome deletion that epigenetically silenced Clr5 ultimately locked in a latent, Prm1-independent fusion pathway involving the mating regulator Ste11 and the pheromone-MAPK scaffold Sms1. This adaptation, though efficient at restoring sexual reproduction, came at the cost of mitotic growth. By contrast, rationally constructing mutations in a “synthetic evolution” approach bypassed the trade-off, enabling simultaneous high fusion efficiency and near–wild-type proliferation. Our findings illustrate how epigenetic de-repression and genetic assimilation converge to create innovative solutions under cyclical selection pressures with trade-offs.

## Introduction

Cellular evolution emerges from the interplay between the genome and the environment, which together define the space of possible adaptive solutions. Genomic alterations such as large-scale rearrangements and aneuploidy can provide rapid but often costly ways to compensate for lethal drug stress or essential gene loss^1–4^. In parallel, cells rely on transcriptional and epigenetic plasticity to enact rapid, reversible responses to stress. Such plasticity can generate heterogeneous cell states, influencing adaptive cell fate decisions within an isogenic population^5,6^. In cancer, for example, sublethal drug exposure induces chromatin remodeling that transiently reprograms gene expression, conferring reversible drug tolerance ^7–9^. Adaptation therefore emerges not from the genome alone but from a multilayered regulatory system, where genomic changes, transcriptional programs, and epigenetic mechanisms interact to shape cellular evolvability. Yet how these layers coordinate over evolutionary timescales to transform transient responses into stable adaptive states remains poorly understood.

The nature and dynamics of environmental stress further sculpt these trajectories. Constant pressure often drives the emergence of specialists streamlined for narrow niches, while unpredictable fluctuations favor generalists with broad tolerance^10–13^. Periodic, predictable cycles such as seasonal changes or nutrient oscillations can instead promote oscillatory or alternating strategies, where populations repeatedly shift between adaptive states. Experimental evolution studies illustrate this principle: yeast populations exposed to alternating salt and oxidative stress evolve cross-protection, such that adaptation to one stressor confers resistance to another ^14,15^. At the population level, these outcomes reflect strategic behaviors—trade-offs, bet-hedging, and oscillatory dynamics—that emerge from genome–environment interactions^11,16^. Trade-offs arise when mutually exclusive programs, such as mitosis and meiosis, compete for shared resources^17,18^

Despite their theoretical prominence, direct experimental evidence for how transient strategies become stabilized into heritable traits has remained limited. Genome sequencing has catalogued mutations across long-term evolution^19–21^.Yet the dynamic relationship between short-term regulatory plasticity and long-term genomic change remains unclear. Do transcriptional and epigenetic shifts provide durable solutions on their own, or must stable adaptation ultimately be encoded at the genomic level?

A prime example of how cells navigate shifting stress regimes is the transition from the mitotic to the meiotic cycle in *Schizosaccharomyces pombe*, which is triggered by nitrogen starvation. This switch involves profound reorganization, encompassing epigenetic, transcriptional, translational and post-translational regulation^22–28^. In turn, sexual reproduction enhances adaptation by facilitating genomic recombination^28^. At the transcriptional level, the master regulator s*te11* integrates multiple upstream signaling pathways to activate sexual differentiation programs^29^. Notably, *ste11* expression is itself subject to multi-layered regulation^30^, including epigenetic control via heterochromatin factors such as *clr5*, which modulates chromatin accessibility and thereby influences developmental commitment^31^. Together, these mechanisms highlight how the S. *pombe* sexual cycle provides a powerful model for dissecting the interplay of genomic, epigenetic and transcriptional factors in driving adaptive transitions. Moreover, it offers a tractable system for probing how cells maintain, or fail to maintain, a generalist state when repeatedly challenged by nutrient fluctuations.

Here, we address these questions through experimental evolution of *prm1*Δ mutants, in which deletion of a pivotal fusion protein, imposes severe defects in sexual reproduction ^32,33^. By evolving populations through repeated mitosis–meiosis cycles and profiling their transcriptomes and genomes, we found that cells initially adapted through transcriptional oscillations buffering the fusion defects, but stable rescue emerged from a genomic deletion that induced subtelomeric silencing of the chromatin regulator *clr5*, thereby derepressing a mating program. This genetic assimilation event leveraged non-fusion mating genes *ste11*, *sms1*, and *adg2*, co-opting them into an alternative mechanism for Prm1-independent fusion. Finally, synthetic evolution demonstrated that rational design can bypass the natural constraints of trade-offs, producing genotypes that balance fusion and growth more effectively than spontaneous evolution. Our results reveal how genome–environment interplay drives the transition from plastic strategies to genetic assimilation, illuminating general principles of evolvability under stresses.

## Results

### Rapid adaptive evolution of fusion in cells lacking the Prm1 fusion protein

Prm1 is a critical factor for cell fusion. While expressed at low levels during vegetative growth, the gene is expressed at higher levels in mating conditions^33^, and Prm1 protein localizes to the cell-cell fusion site (Figure 1A)^34^, largely overlapping with the type V myosin Myo52 that marks the actin fusion focus^35^, and the sterol probe D4H labelling the plasma membrane^36^. As previously reported^33^, deletion of *prm1* in both mating partners causes a severe fusion defect, reducing fusion efficiency from nearly 100% to ∼ 8% (Figure S1A-B, Table S1B; fusion efficiency quantified by microscopy as percentage of fused cell pairs amongst all cell pairs). As in *S. cerevisiae*^32^, *prm1* deletion in *S. pombe* blocks cell fusion after cell wall digestion, as illustrated by the formation of membrane invaginations between partner cells (Figure 1B). The severe, yet incomplete, fusion defect of bilateral *prm1*Δ mutants contrasts with the milder phenotypes in unilateral crosses^33^ and in other species^32^, suggesting that Prm1-independent pathways were evolutionarily plastic and alternative fusion mechanism could be potentiated by experimental evolution. Hence, we designed an asexual-sexual alternative evolutionary experimental scheme by selecting rare *prm1*Δ spores, which form upon successful cell fusion.

**Figure 1:**
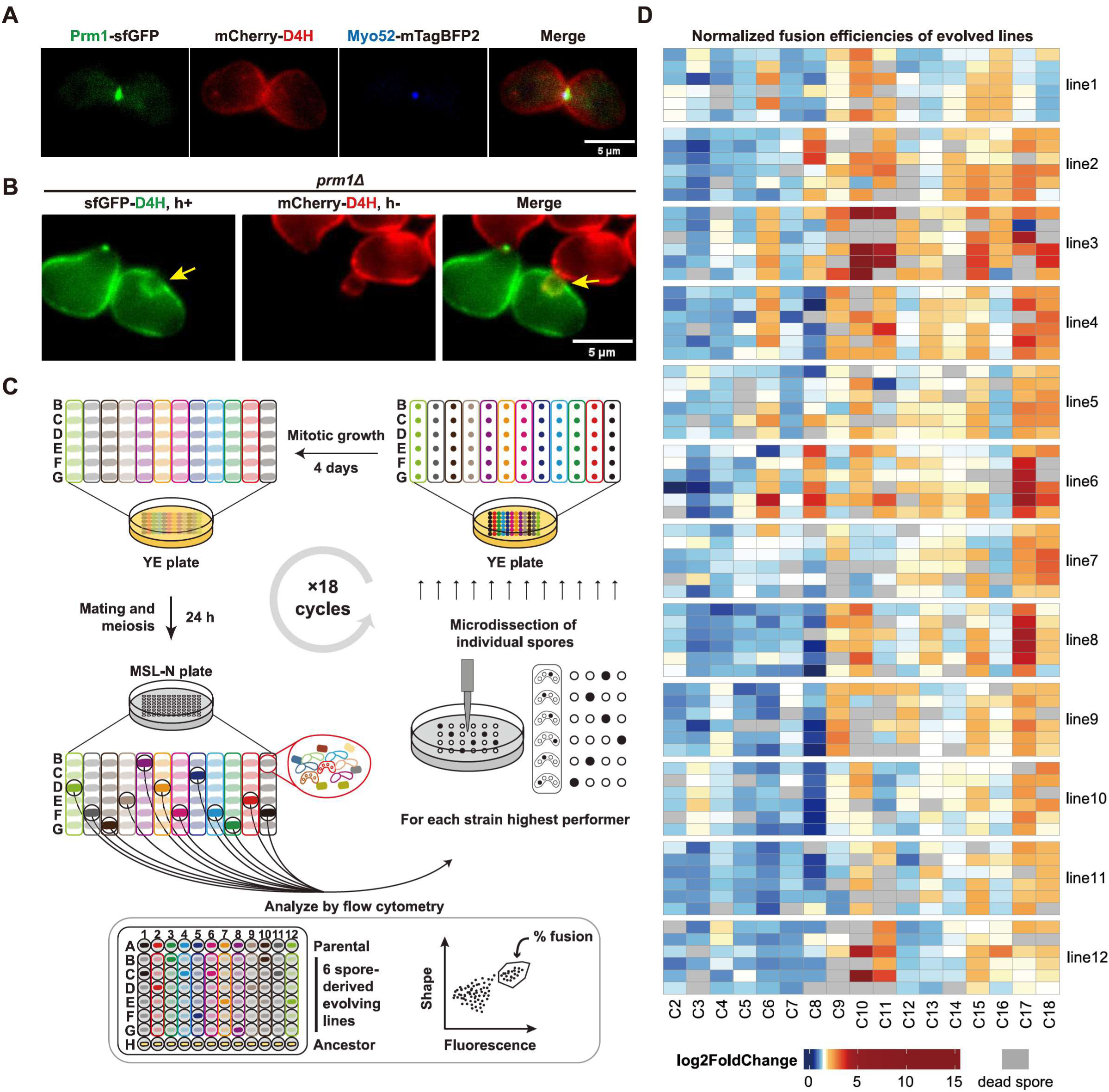
*prm1*Δ cells fusion defect and experimental evolution workflow. (**A**) Maximum intensity projection images from an inverted fluorescence microscope of *h90* wild-type strains expressing Prm1-sfGFP, mCherry-D4H (used to label the plasma membrane^36^) and Myo52-mTagBFP2, during fusion. (**B**) Mating of *h-prm1*Δ mCherry-D4H and *h+ prm1*Δ sfGFP-D4H cells. A yellow arrow points to invagination formed between two mating cells with one red and one green membrane. (**C**) Detailed experimental workflow of the 18 evolutionary cycles. Round shapes represent single cells or spores, elongated shapes represent colonies or groups of cells. The different colours label parallel evolving line. The different shading indicate heterogeneity within each line. (**D**) Heatmap of fusion efficiency fold change in the 12 evolving lines with 6 selected spores at each cycle, measured by flow cytometry. Dead spores are indicated in light grey (Table S1A).

To experimentally test whether *prm1*Δ cells could evolve enhanced fusion efficiency, we conducted a laboratory evolution experiment involving repeated cycles of nitrogen starvation, inducing sexual reproduction and meiosis, followed by spore selection, and growth on rich medium, selecting for biomass amplification through mitosis. We first generated a reporter strain expressing a *pmeu31*-GFP cassette, specifically activated during meiosis upon successful cell fusion, facilitating high-throughput screening (Figure S1C-D). We then isolated 96 single *prm1*Δ cells and induced mating and sporulation on MSL-N plates. After 24 hours, the percentage of GFP-positive cells was quantified using flow cytometry amongst all cells, as proxy for fusion efficiency. Note that this indirect quantification amongst all cells yields lower figures (∼29% in WT; ∼5% in *prm1*Δ; Figure S1D). The 12 colonies displaying the highest fusion efficiencies were chosen as the founders of 12 parallel evolving lines.

The experimental evolution scheme is presented in Fig. 1C. From each selected colony, we went back to the MSL-N plate and selected individual spores from six independent tetrads by tetrad dissection, ensuring a stringent selection for the fusion-positive phenotypes. These spores were subsequently cultured vegetatively on rich medium for 4 days to amplify biomass. The resulting colonies were transferred back to MSL-N plates for 24 hours to induce the next cycle of mating, sporulation, and fusion quantification and spore selection. We established a systematic evolutionary scheme where the 12 parallel lines were arrayed on columns 1 to 12 of a 96-well plate. Within each column, row A contained the parental colony; rows B-G contained the 6 amplified spores from the parent; row H consistently contained 12 freshly thawed ancestor *prm1*Δ controls from −80°C stocks to ensure comparability across cycles. In each subsequent round, we repeated this selective dissection process, identifying for each line the single highest-performing strain among the six spore-generated colonies by flow-cytometry, and micro-dissecting six spores from that strain for the next round (Figure 1C). This iterative selection mechanism aimed to facilitate strong phenotype retention while simultaneously enabling adaptive diversification through independent evolutionary routes.

Throughout this evolutionary experiment, we recorded all flow cytometry-measured fusion efficiencies (normalized to the simultaneously tested ancestor strain to account for experimental variability) and compared them to the initial ancestor strain (Table S1A). Remarkably, fusion efficiency did not increase linearly but rather displayed cyclical fluctuations across evolutionary cycles (Figure 1D, Table S1A). Eventually, a notable overall increase was achieved in all evolving lines: the ancestor fusion rate (∼8%) significantly increased to 10-34% (absolute fusion efficiency measured by microscopy) in the evolved strains (Figure S1E, Table S1B). Notably, line 3 exhibited the highest fusion efficiency, which appeared at, and remain high from, the 9th cycle (Figure 1D, S1E).

Given these fluctuating patterns, we hypothesized that the adaptive increases in fusion efficiency might result from transient changes at the transcriptome level rather than stable genomic modifications. To test this hypothesis, we first proceeded to detailed transcriptomic profiling of these evolved strains.

### Evolutionary trajectories reveal transient transcriptomic shifts in dynamic

We performed RNA-seq on the strains with highest fusion-efficiency selected from each evolutionary line at every third evolutionary cycles (cycles 3, 6, 9, 12, 15, and 18). Triplicate RNA-seq data were collected from both vegetative growth (0 hours) and 5 hours post-starvation conditions.

Through systematic pattern analysis, we found that gene expression changes exhibited variation patterns across evolutionary time points in most strains (Figure 2A, Figure S2-3, Table S1C-D). We analyzed the amplitude of variation across all 216 transcriptomic profiles for samples in mating from 5 hours post-starvation condition and focused on the patterns of the top 20% of genes that displayed the greatest variability (Figure S4A). The other 80 % of the genes did not exhibit significant variation in abundance or patterns, ruling out batch effect in the RNA sequencing. Notably, line 3 exhibited a markedly distinct pattern among all profiles (Figure 2A). Upon analyzing line 3 separately, we discovered that its highly variable genes skewed the overall variability (Figure S4B, Table S1E). In order to analyze common changes in all evolved lines, we defined a set of highly variable genes (HVG, 1,290 genes) by combining the top 20% most variable genes from other strains with the top 20% from line 3 (Figure S4C, Table S1E). Unsupervised clustering analysis of the HVG set revealed that these genes formed four clusters and could be classified by Gene Ontology (GO) terms in three categories (Fig 2A): 1) pheromone response MAPK cascade (terms in orange); 2) metabolic process (terms in green); and 3) cell cycle related processes (terms in blue).

**Figure 2:**
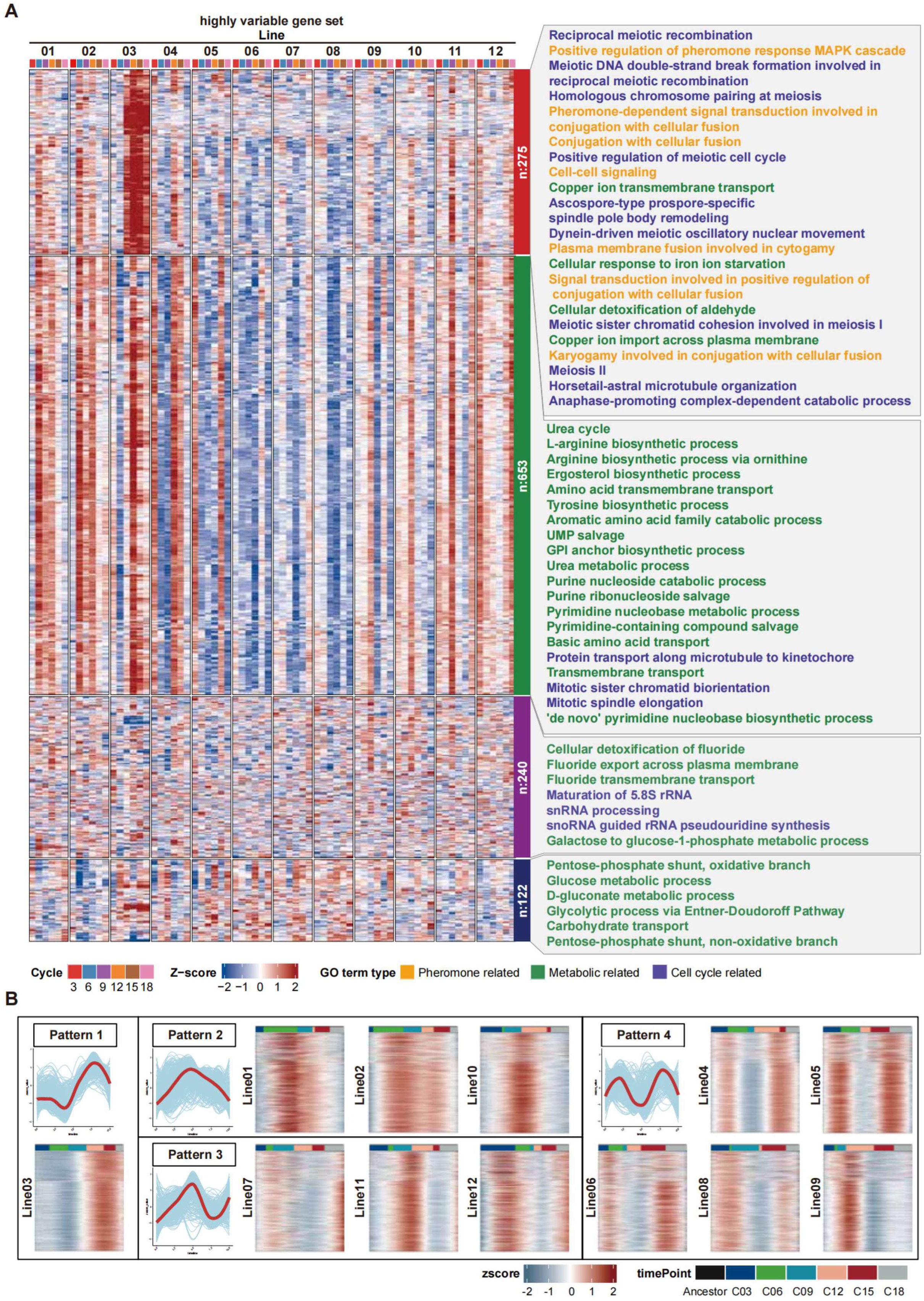
Oscillatory transcriptomic patterns during evolutionary timeline. (**A**) Fuzzy c-means clustering of the highly variable gene set transcriptomic profile 5h after nitrogen starvation. The samples are single individual strains with highest fusion efficiency at evolutionary cycles 3, 6, 9, 12, 15 and 18 in each 12 evolving lines. The right panel shows Gene Ontology (GO) enrichment analysis of Biological Process terms for the four resulting clusters. GO terms with statistical significance (p < 0.01) are shown, with color-coded categories: pheromone-related (orange), metabolic process-related (green), and cell cycle-related (blue). (**B**) Trajectory inference analysis patterns of the 928 highly variable genes exhibiting oscillatory dynamics (corresponding to clusters 1 and 2 in (A); Figure S4A-C) in the 12 evolving lines, grouped into four representative evolutionary patterns. In each box, plots show expression dynamics over pseudo-time, with individual gene trajectories (blue lines) and average trend for the pattern (red line); heatmaps show Z-scored, gene-normalized expression for each lineage assigned to a given pattern. The coloured bar above each heatmap represents the evolutionary cycles, with its length scaled to the inferred pseudo-evolutionary time of that specific lineage.

To investigate the potential oscillatory patterns observed in our clustering analysis, we used trajectory inference — a computational method that arranges samples along a pseudo-time scale based on gene expression similarity — to reconstruct the distinct gene expression trajectories for each evolving lineage^37,38^. This analysis revealed that the majority of genes (928, Fig S4C in orange circle, Table S1E), corresponding to the first two clusters in Figure 2A, exhibited oscillatory dynamics across the 12 parallel evolving lines (Figure 2B, S4D), with distinct evolutionary expression patterns characterized by periodic fluctuations (patterns 2 to 4). Pattern 4 exhibited highly frequent oscillations, indicating substantial transcriptomic variability between cycles. Uniquely, line 3 exhibited an oscillatory pattern in the beginning but stabilized significantly starting at cycle 9 (pattern 1), suggesting the establishment of a robust adaptive expression state.

The resemblance between oscillatory transcriptomic (Figure 2A) and the fluctuating cell fusion efficiency (Figure 1D) suggests that the oscillatory changes could be adaptive. This hypothesis is supported by functional enrichment analyses, showing that oscillating genes were significantly enriched in functions related to pheromone signaling and metabolic processes— pathways critical for modulating cell fusion under nitrogen-limiting conditions—and the cell cycle. We hypothesize that these oscillatory dynamics are a systems-level consequence of the cell’s regulatory network navigating the conflicting selective pressures of mitotic growth and sexual reproduction.

### Transcriptome Trajectories Trace the Mitosis–Meiosis Balancing Act

The oscillatory transcriptional pattern, also observed in the mitotic growth samples, prompted an exploration into potential trade-offs between mitotic and meiotic cycles (Figure S2, Table S1D). To this end, we measured both growth rates and fusion efficiency across all 72 evolved strains in triplicate (Table S1I-J). Our analysis revealed an inverse correlation between fusion efficiency and growth rate, with the most significant shift occurring between cycles 9 and 12 (Figure 3A, Table S1I-J). This inverse correlation weakened in subsequent generations with both growth rate and fusion efficiency improving at cycle 18 (Figure 3A).

**Figure 3:**
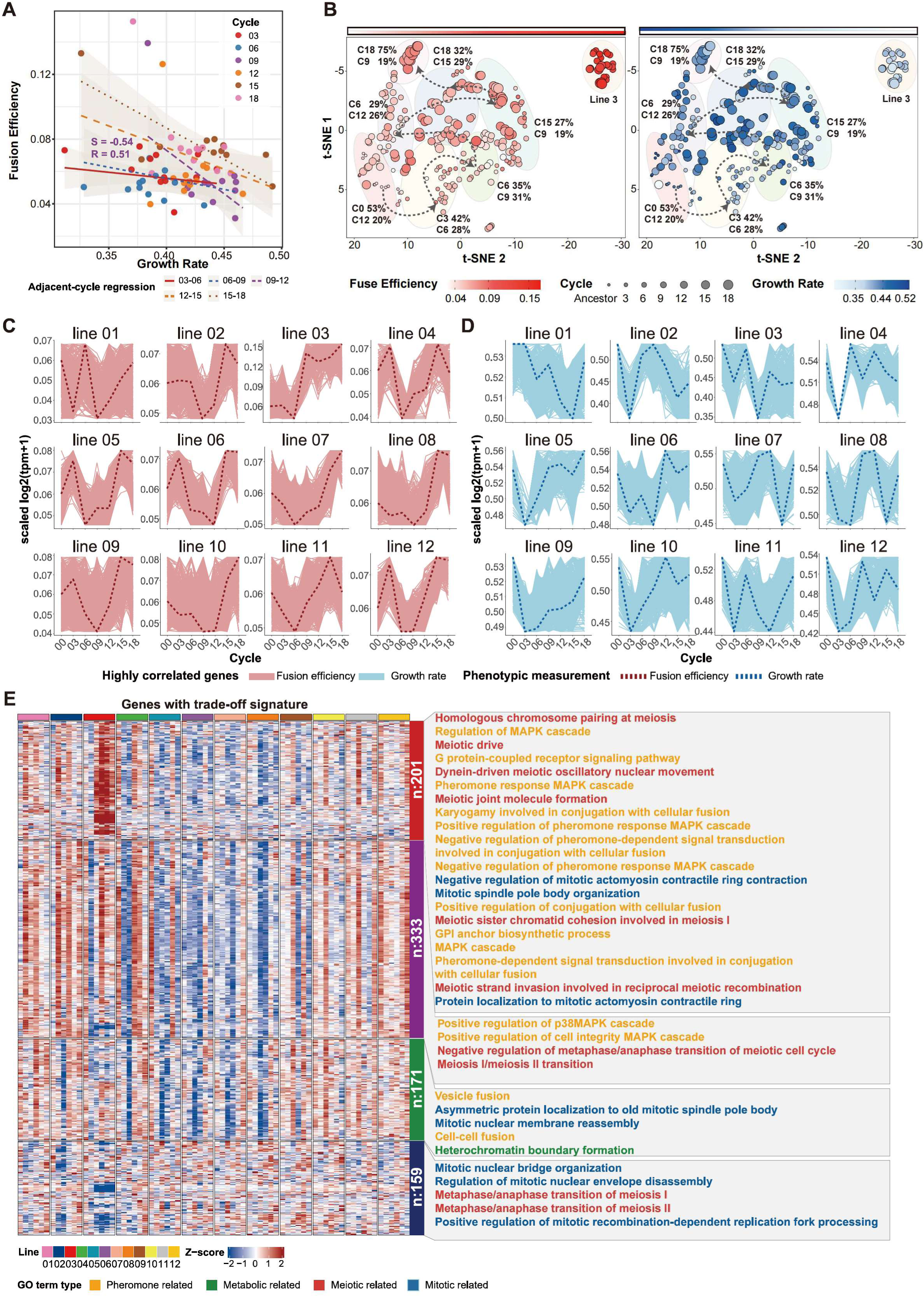
Transcriptomic and phenotypic co-analysis reveals a sex-growth trade-off. (**A**) Linear regression analysis of mean mating efficiencies (measured by Flow Cytometry in triplicates) and growth rates in all samples (Table S1I-J) selected for further analysis (same strains as in Figure 2). S = slope; R = correlation score. The S and R shown on the figure show the slope for samples of cycle 9 to cycle 12. Other S and R values can be found in Table S1J. (**B**) Unsupervised t-SNE clustering of all evolved sequenced samples (in triplicates; same strains as above), along with ancestor strain controls. Each dot represents one sample. Left and right panels are colored according to phenotypic measurements: fusion efficiency (red gradient) and growth rate (blue gradient), respectively. Dot size corresponded to the evolutionary cycle, from small to large. Major t-SNE clusters are circled in semi-transparent ovals. Dashed lines indicate a trajectory determined by the dominant cycle in each cluster. For each cluster, C# indicates the proportion of samples in the dominant and following evolutionary cycle (C = cycle; # = cycle number). The isolated cluster of line 3 was excluded from the trajectory analysis. (**C**) Expression trajectories (red lines) of the top 10% of genes in each line with tight alignment to fusion efficiency (red dashed line, relationship measured by LSE; Table S1G,I) from RNAseq data 5h post nitrogen starvation (log2(TPM+1), min-max scaled per lineage); X axis = evolutionary cycles; Y axis = normalized scale of each line. (**D**) Expression trajectories (blue lines) of the top 10% of genes with tight alignment to growth rate (blue dashed line, relationship measured by LSE; Table S1G,J) from RNAseq data during vegetative growth (T0; log2(TPM+1), min-max scaled per lineage); X axis = evolutionary cycles; Y axis = normalized scale of each line. **(E**) Fuzzy c-means clustering of the transcriptomic profile of the curated a set of 864 genes with trade-off signature (i.e. showing opposite correlation with fusion efficiency and growth rate; from 3C-D and S4E-F; Table S1G) from RNAseq 5h post nitrogen starvation. The right panel shows Gene Ontology (GO) enrichment analysis of Biological Process terms for the four resulting clusters. GO terms with statistical significance (p < 0.1) are highlighted, with color-coded categories: pheromone-related (orange), metabolic-related (green), mitotic cycle-related (blue) and meiotic cycle-related (red).

To visualize the relationship between transcriptomic changes and fitness phenotypes, we performed t-SNE clustering on the transcriptome profiles of evolved strains (Figure 3B, Table S1F). In the resulting plots, each point represents an individual sample, with its size corresponding to the evolutionary cycle and its red color shade indicating fitness for cell fusion. The unsupervised clustering arrays strains across two main axes, with fusion efficiency increasing towards the plot top-right corner. Clusters (highlighted by the pale colored ellipses), identified by the predominant evolutionary cycle within each successive cluster, show a zig-zagging evolutionary trajectory, thus mapping the population’s progression through time (indicated by dashed arrows). Mapping of the growth rate (blue) showed an oscillation between high and low growth rates, with clear anti-correlation between red and blue shades for individual strains. This zig-zagging trajectory reflects the oscillatory patterns in gene expression and fitness, with an overall displacement towards higher fusion efficiency. Note that line 3’s special pattern was clustered to the upright corner, showing a particularly strong trade-off between high fusion efficiency and low growth rate.

To dissect the molecular drivers of the phenotypic trade-off, we pursued a two-pronged approach. First, to empirically identify genes linked to fitness, we analyzed transcriptome profiles from two key conditions: vegetative growth (for growth rate) and 5h post-nitrogen starvation (for fusion efficiency). Using a least-squares estimate (LSE) method, we identified the top 10% of genes whose expression trajectories showed the highest positive or negative correlation with the growth rate and fusion efficiency trajectories for each of the 12 evolved lines (Fig 3C-D, S4E-F). From this initial pool, we curated a final set of 864 genes that exhibited a clear trade-off signature: their expression was positively correlated with fusion efficiency while being negatively correlated with growth rate, or vice versa in at least one line (Table S1G). As anticipated, a Gene Ontology (GO) enrichment analysis of this gene set revealed functions in cell cycle control, as well as functions critical for fusion such as plasma membrane remodeling, cellular polarity, and vesicle dynamics (Figure 3E). These analyses linking transcriptomic profiles to phenotypic traits suggests that multiple biological processes, particularly those related to the cell cycle, contribute to the observed oscillatory patterns. Such oscillations may represent an adaptive strategy enabling cells to navigate periodically changing environments.

Taken together, these data suggested that the phenotypic trade-off between growth and fusion is governed by an underlying regulatory conflict between mitotic and meiotic gene expression programs in the first few cycles of evolution.

### Genomic rearrangement-mediated epigenetic *clr5* silencing boosts fusion efficiency in the fittest e-Strain

Among all evolved strains, line 3 stands out in reaching highest fusion efficiency (Figure 1D), exhibiting distinct HVG expression pattern (Fig 2A-B) and forming a stand-alone cluster in t-SNE analysis (Fig 3B). While transient transcriptional plasticity appears to be a common adaptive strategy, enabling a dynamic balance between mitosis and meiosis, line 3 seemingly found a way to escape this trade-off, reaching a high level of fusion from cycle 9. The line 3 cycle 9 strain is hereafter referred to as the *fittest evolved strain* (e-Strain). We examined whether a stable genomic alteration could be responsible for cementing the phenotype in this fittest strain.

To uncover the underlying mechanism, we performed whole-genome sequencing across all evolved lines at six evolutionary timepoints, the same strains for which we did RNA-seq analysis. Most strains accumulated only sparse single-nucleotide variants (SNVs), few indels, or structural variations, predominantly located in noncoding regions and increasing slowly over time (Figure 4A, Figure S5A-B, Table S1H). Furthermore, the different lines shared very few common genomic changes other than the ones that already existed in the *prm1*Δ ancestor strain (Figure 4A, Figure S5C), suggesting that the adaptive fusion efficiency phenotype most likely was not due to these genomic changes.

**Figure 4:**
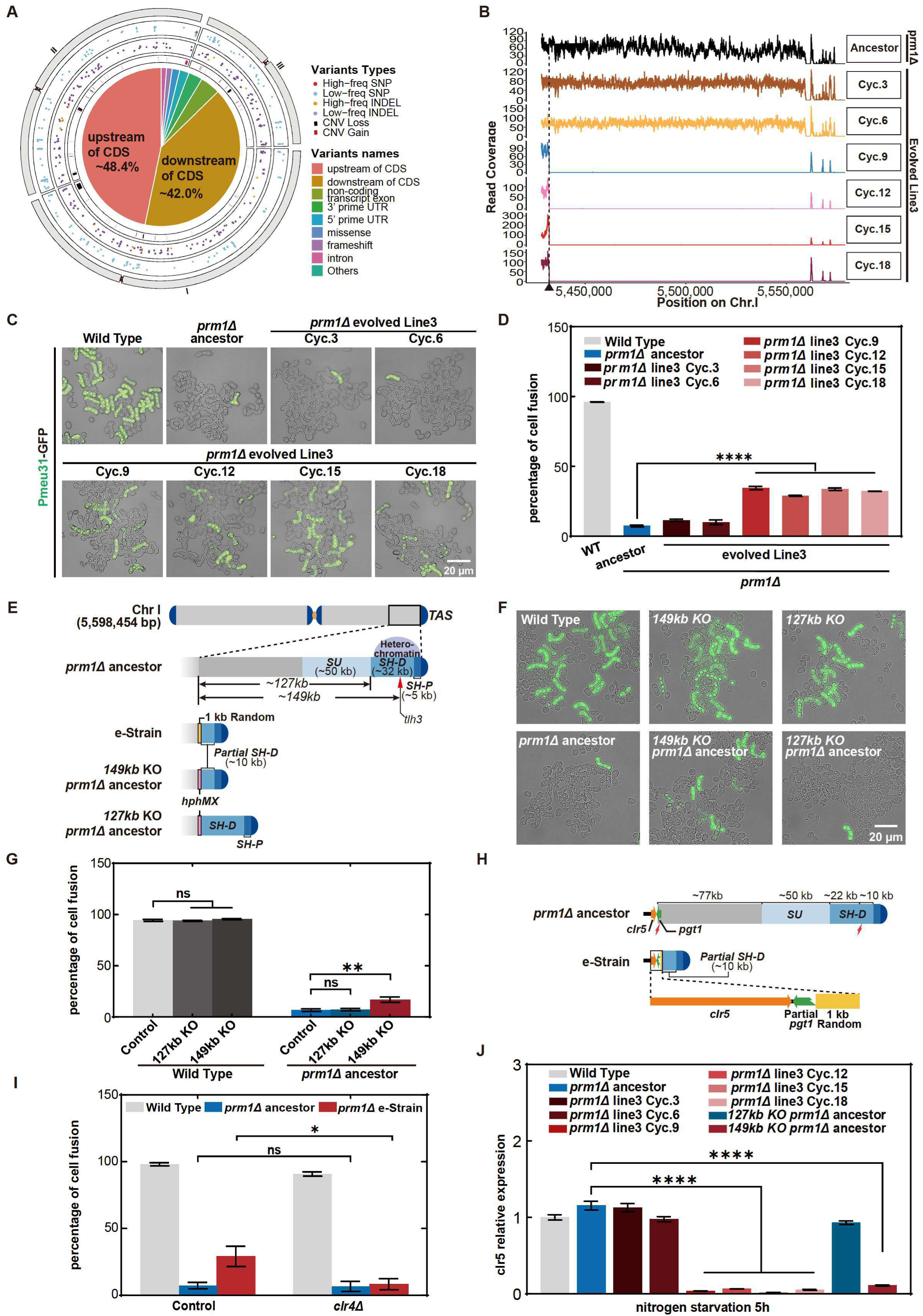
Loss of *clr5* expression at the edge of a 149 kb deletion in the fittest evolved strain. **(A)** Summary of all genomic mutation identified in the 12 evolving lines arrayed over the entire genome. Mutations are categorized by SNP, indel and CNV. The central pie-chart classifies the various mutations. (**B**) Chromosome I telomere adjacent region sequencing coverage in *prm1*Δ ancestor and evolving line 3. (**C**) Representative fusion efficiency pictures of *h90* WT, *prm1*Δ ancestor and evolving line 3 strains. Successful fusion is shown by *p^meu31^-sfGFP* (green), which is only expressed after successful fusion. (**D**) Quantification of fusion efficiency in strains as in (C) from microscopic picture (N ≥ 1000 for each sample, Table S1B, Supplementary Information 5.4). (**E**) Schematic representation of chromosome I telomeric structure in *prm1*Δ ancestor, line 3 cycle 9 (e-Strain), and 149kb and 127kb knockout (KO) strains. Random indicates a ∼1kb sequence with unknown identity after blast search (Figure S6A). (**F**) Representative fusion efficiency pictures of *h90* WT and *prm1*Δ ancestor strains, in the presence or absence of the indicated large-scale knockouts (149 kb KO and 127 kb KO), 24h post nitrogen starvation. Successful fusion is shown by *p^meu31^-sfGFP* (green). (**G**) Quantification of fusion efficiency in strains as in (F), N ≥ 500 for each sample (Table S1B). (**H**) Schematic representation of *clr5* location in *prm1*Δ ancestor strain and line 3 cycle 9 strain (e-Strain). (**I**) Quantification of fusion efficiency in *h90* WT, *prm1*Δ ancestor and *prm1*Δ e-Strain, with or without *clr*4Δ, 48h post nitrogen starvation from microscopic picture (N ≥ 100 for each sample, Table S1B). (**J**) qPCR quantification of *clr5* expression in WT, *prm1*Δ ancestor,evolving line 3, 127kb KO *prm1*Δ ancestor and 149kb KO *prm1*Δ ancestor (Table S1K) at 5h post nitrogen starvation.

In contrast, strain Line 3, harbored a unique 149-kb deletion adjacent to the telomeric region of chromosome I (Figure 4B). This mutation was first detected in the e-Strain at cycle 9 and became fixed in subsequent generations (Figure 4B), a timeline that perfectly corresponded with the rescue and stabilization of the low fusion efficiency phenotype in *prm1*Δ cells (Figure 4C-D, Table S1B). The compelling correlation between this large genomic deletion and the rescued phenotype prompted us to hypothesize that it was the adaptive mutation responsible for the increased fusion efficiency.

To test this hypothesis, we first sought to characterize and engineer the deletion manually. Long-read sequencing of the e-Strain confirmed the precise deletion junctions, revealing the replacement of the region with a ∼1-kb insert (Figure 4E, S6A). Notably, the breakpoint is adjacent to one of the paralogs *tlh3* locus, suggesting the deletion may have resulted from a subtelomeric rearrangement with other *tlh* genes^39^. The deleted sequence encompassed the entire subtelomeric unique (SU) region and the majority (22 kb of 32 kb) of the distal subtelomeric homologous (SH-D) region, leaving the proximal subtelomeric homologous (SH-P) region intact^39–42^. We then engineered this 149-kb deletion (∼1.2% of the *S. pombe* genome) in the ancestral *prm1*Δ strain (Figure 4E). The resulting *prm1*Δ 149kb KO cells exhibited a significantly increased fusion efficiency of ∼17 %, confirming that the 149-kb deletion is sufficient for the phenotypic rescue and is indeed an adaptive mutation (Figure 4F-G, Table S1B).

We next investigated whether the entire 149-kb deletion was necessary for the rescue. We first reasoned that the loss of one or several genes might be responsible. However, individual deletions of seven genes within the region, including *isp5* and *pgt1*(Figure S9, highlighted in blue, TableS1I), or larger deletions of 10 kb, 20 kb, and 127 kb (Figure S6B, Table S1I) from the euchromatin side, failed to rescue the phenotype. The 127-kb deletion was specifically designed to leave the SH-D region intact, and its failure to restore fusion efficiency (Figure 4F-G, Table S1B), in contrast to the success of the full 149-kb deletion, suggests that the removal of both the SU and SH-D regions is important for the rescue.

Given that the SU and SH regions are known to buffer against the spread of subtelomeric heterochromatin^41^, we hypothesized that the deletion allows this silencing to extend into adjacent gene-rich regions, downregulating a gene whose function regulates fusion. To test this, we disrupted subtelomeric silencing by deleting *clr4*, coding for histone H3K9 methyltransferase essential for this process. While *clr4*Δ in *prm1+* strain had no obvious fusion defect, deleting it in the e-Strain reverted its fusion efficiency to the low levels of the *prm1*Δ ancestor (Figure 4I, Table S1B), confirming that subtelomeric silencing is essential for the rescue. We then examined the expression of *clr5*, the first intact heterochromatin-related gene adjacent to the chunk deletion (Figure 4H). We found that *clr5* mRNA levels were significantly decreased in the evolved strains after cycle 9 and in the engineered 149ko strain, but not in the 127ko strain (Figure 4J, Table S1K), suggesting that the adaptive 149-kb deletion leads subtelomeric silencing to spread to *clr5*.

We hypothesized that the strong decrease in *clr5* expression, mediated by the 149kb deletion, is the adaptive rescue of fusion efficiency in the evolved strain. Direct visualization of endogenously tagged Clr5-mCherry showed a marked decrease in protein levels in the e-Strain fusing cells compared to controls (Figure 5A). Furthermore, deletion of *clr5* in the *prm1*Δ ancestor background was sufficient to increase fusion efficiency to 40% (Figure 5B-C, Table S1B), demonstrating that loss of *clr5* is sufficient to recapitulate the evolved phenotype. By contrast, deletion of *clr5* in the evolved strain did not boost fusion efficiency, and overexpression of *clr5* in the evolved strain significantly decreased the fusion efficiency of the evolved strain (Figure S9, Table S1I).

**Figure 5:**
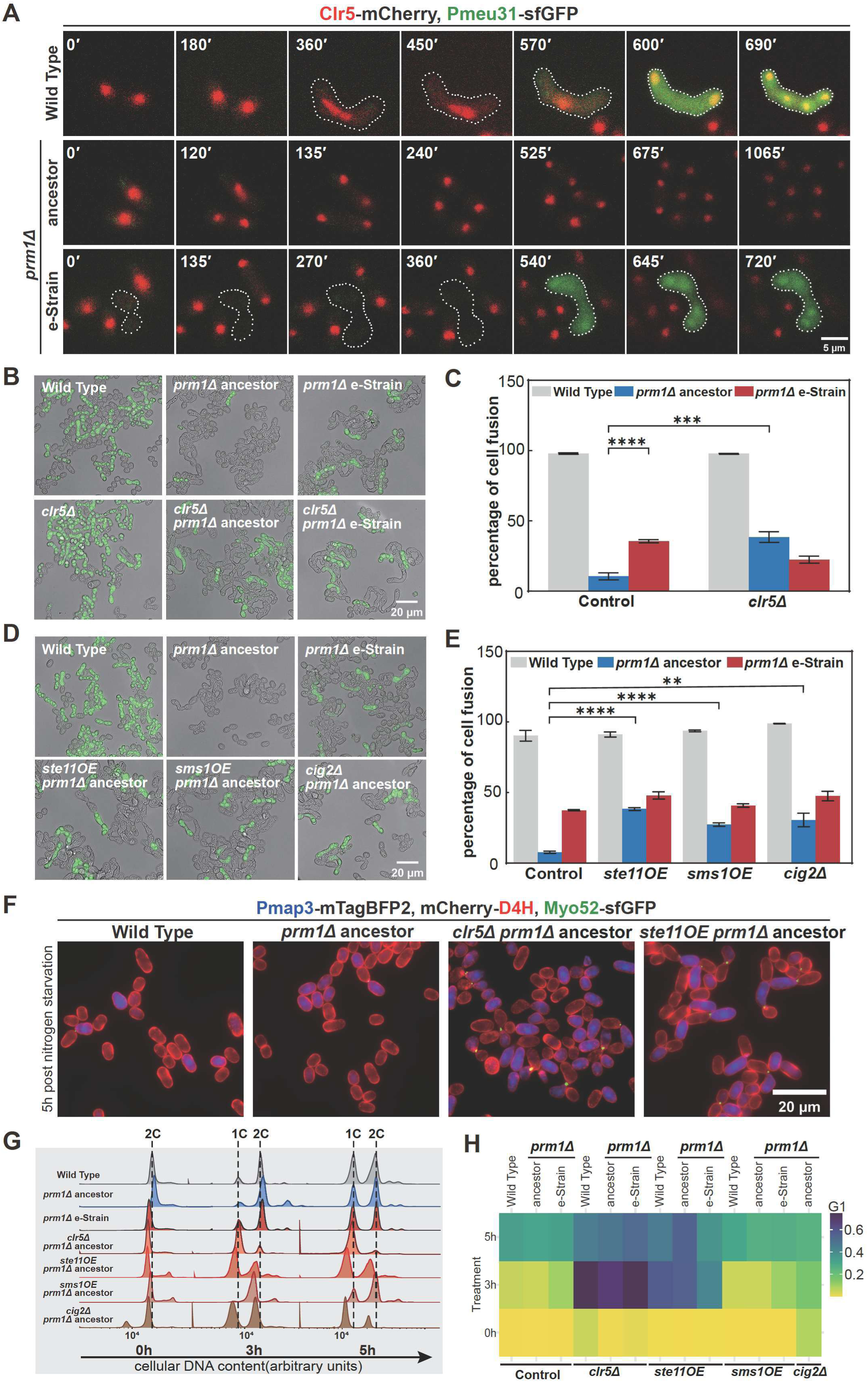
Dissecting the genetic compensatory pathway of the evolved *prm1*Δ strain. (**A**) Localization of Clr5-mCherry during mating in *h90* WT, *prm1*Δ ancestor and *prm1*Δ e-Strain. *p^meu31^*-sfGFP is expressed in zygotes (dashed outlines). In the time lapse, t0 corresponds to 5h post nitrogen starvation in MSL-N liquid. **(B**) Representative fusion efficiency pictures of *h90* WT, *prm1*Δ ancestor and *prm1*Δ e-Strain, with or without *clr5*Δ, 48h post nitrogen starvation. Successful fusion is shown by *p^meu31^-sfGFP* (green). (**C**) Quantification of fusion efficiency in strains as in (B), N ≥ 1000 for each sample (Table S1B). (**D**) Representative fusion efficiency pictures of *h90* WT, *prm1*Δ ancestor, and *prm1*Δ e-Strain, as well as *prm1*Δ ancestor with *ste11*OE, *sms1*OE and *cig2*Δ, 48 hours post nitrogen starvation. Successful fusion is shown by *p^meu31^-sfGFP* (green).(**E**) Quantification of fusion efficiency in strains as in (D), N ≥ 1000 for each sample (Table S1B). (**F**) Representative fluorescence microscopy images capturing early mating events in WT, *prm1*Δ ancestor, and *prm1*Δ ancestor with *clr5*Δ or *ste11*OE, 5h post nitrogen starvation. All strains express *p^map3^*-mtagBFP2 (strongly induced during mating in P cells), mCherry-D4H (labeling the plasma membrane) and Myo52-GFP (labeling the actin fusion focus). Quantification is shown in Figure S10E-H. **(G**) DNA content profiles in *h90* WT, *prm1*Δ ancestor, e-Strain, and *clr5*Δ*, ste11*OE*, sms1*OE and *cig2*Δ in the *prm1*Δ ancestor background, measured at 0, 3, and 5h of nitrogen starvation. Cells were stained with propidium iodide and analyzed by flow cytometry (Table S1L). (**H**) Heatmap showing the percentage of the G1-arrested population in the control, measured as in (G) for indicated strains (Table S1L).

Together, these data indicate that a spontaneous structural variant in line 3 led to position-effect-mediated epigenetic silencing of *clr5*, which in turn stabilized an otherwise transient transcriptional adaptation to increase cell fusion efficiency.

### Enhancement of fusion success in *prm1***Δ** cells by repeated trials

Clr5 is a chromatin-silencing factor that functions independent of histone H3K9 methylation to silence mating-type cassettes and other genomic loci, including the *ste11* gene^31^. This suggests Clr5 promotes alternative fusion mechanisms indirectly through gene regulation. To pinpoint the key effectors of the alternative fusion pathway, we used several bioinformatic approach to generate a high-confidence list of candidate genes. First, we performed a comparative transcriptomic analysis between the e-Strain, the engineered *clr5*Δ *prm1*Δ strain and the *prm1*Δ ancestor to identify genes consistently altered in the high-fusion backgrounds (Figure S7). By integrating comparative transcriptomic analysis, using machine learning to identify key genes from all evolving transcriptomics data (Figure S8, Supplementary Information 4.1), and cross-referencing them with our list of fusion-correlated genes (Table S1G, Supplementary information 4.1), we converged on a prioritized list of ∼100 candidates for functional validation (Figure S9, Table S1I). The effect of deletion and/or overexpression of these components on the fusion efficiency of the *prm1*Δ ancestor strain was systematically tested, revealing only a handful of genes with significant effect on *prm1*Δ fusion efficiency.

Among these, the master transcriptional regulator Ste11, which promotes the sexual transcriptional program by inducing expression of all pheromone-MAPK components^43,44^, and a known Clr5 target^31^, was upregulated in the *prm1*Δ e-Strain and *clr5*Δ strains (Figure S7, gene name highlighted in bold), as also confirmed by qPCR in line3 after 9^th^ cycle (Figure S10A, Table S1K). Overexpression (OE) of *ste11* in the *prm1*Δ ancestor increased fusion efficiency to ∼35% but produced no additive effect in the e-Strain, indicating that endogenous Ste11 upregulation was already saturated (Figure 5D-E, Table S1B). Thus, boosting sexual differentiation by Ste11 OE increases fusion efficiency in *prm1*Δ.

To probe downstream effectors, we examined MAPK pathway components. OE of *byr2* ^45^*, ste4* ^46^, *byr1* ^47^, or *spk1* ^48^, coding for the MAP3K, MAP3K cofactor, MAP2K, and MAPK, respectively, had minimal effects (Figure S10B, Table S1I). By contrast, OE of *sms1*, encoding the hemi-arrestin MAPK scaffold that promotes signal transduction through the MAPK cascade^49^, increased fusion efficiency to ∼25% in the *prm1*Δ ancestor background, but not in the evolved strain (Figure 5D-E, Table S1B). Ste11 binds the *P.sms1* prmoter in a one-hybrid assay (Figure S10C), confirming *sms1* as a transcriptional target of Ste11^50^, which was indeed more highly expressed, like Ste11, since cycle 9 in evolved line 3 (Figure S10A, D Table S1K). In agreement with this, *sms1* expression was upregulated during meiosis induction in both *prm1*Δ e-Strain (Figure S7A) and *clr5*Δ in *prm1*Δ ancestor strains (Figure S7B). These data support the notion that adaptive fusion in the stably evolved *prm1*Δ line involves the activation of the Clr5-Ste11-Sms1/MAPK module.

With key genetic players unmasked, we next set out to determine the specific cellular processes by which *clr5*Δ and *ste11* OE strains promote cell fusion in *prm1*Δ ancestor background. We first observed that these strains exhibited accelerated mating behaviors, with expression of the mating-responsive *Pmap3*-mTagBFP2 in more cells (Figure 5F), faster fusion focus formation (indicated by Myo52-GFP, Figure 5F), and increased flocculation and mating pairs formation (Figure 5F, Figure S10E-H, Table S1B) at 5 hours post nitrogen starvation. This overall faster progression into the sexual cycle suggested an early G1 arrest. Indeed, DNA content analysis showed a higher G1 population in the e-Strain 3h post-nitrogen starvation, an early arrest exacerbated in the *clr5*Δ *prm1*Δ ancestor and OE of *ste11*strain but not *sms1* (Figure 5G-H, Table S1L). We next investigated whether cell cycle arrest was sufficient to promote higher cell fusion in *prm1*Δ cells. Remarkably, deletion of the B-type cyclin Cig2, a G1/S cyclin promoting S-phase entry^51^, not only arrested cells faster in G1, but also significantly increased fusion efficiency to ∼24% in the *prm1*Δ ancestor (Figure 5D-E, Table S1B). Thus, one mechanism of cell fusion enhancement in absence of Prm1 is to promote early mitotic arrest, likely enhancing the time for fusion attempts through parallel mechanisms.

Indeed, we observed that deletion of *clr5* or overexpression of *ste11* increased the fraction of *prm1*Δ cells successively trying to mate with more than one partner regardless of eventual fusion success (Figure 6A-B, Figure S11A, Movie S1,Table S1M). Unexpectedly, overexpression of *sms1* also promoted “second-partner” fusion events (Figure 6C), even though it did not induce faster G1 arrest. In these cells, patches of Cdc42-GTP (labeled by the CRIB bioreporter ^52^), which define zones of active exploration for partner cells ^53,54^, were more prominent than in the parental *prm1*Δ ancestor (Figure 6D, Table S1M, Movie S1). This suggests that Sms1 also increases fusion efficiency by promoting mating encounters, likely by enhancing pheromone-MAPK signalling^49^. Thus, one mechanism to bypass the fusion defect of *prm1*Δ is not to repair the defect, but to promote repeated trials, thus increasing overall success.

**Figure 6:**
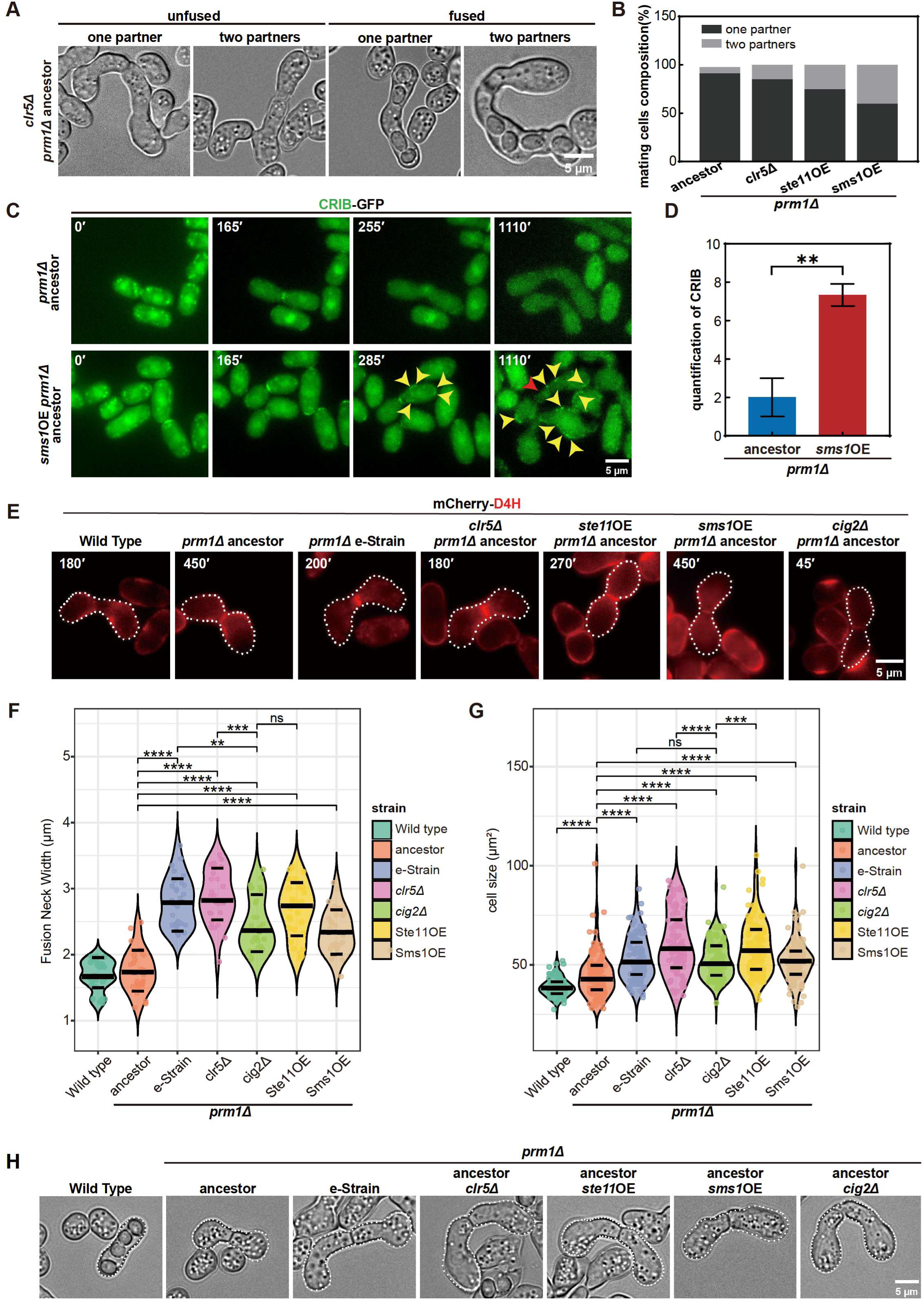
Dissecting the cellular compensatory behavior of the evolved *prm1*Δ strain. (**A**) Representative light microscopy images of *prm1*Δ ancestor strain with *clr5*Δ, showing mating pairs in both unfused and fused conditions, with or without additional partners. (**B**) Quantification of the fraction of cells attempting mating with one or more partners, as shown in (A). N ≥ 100 for each sample (Table S1M). (**C**) Representative fluorescence microscopy images of CRIB-GFP (labelling Cdc42-GTP patches) in *prm1*Δ ancestor with or without *sms1OE*. In the time lapse, t0 corresponds to 5h post nitrogen starvation in MSL-N liquid. Red arrowhead points to successful fusion; yellow arrowheads point to CRIB-GFP patches. (**D**) Quantification of the number of CRIB-GFP patches in each cell as shown in (C). N ³ 42 mating pairs for each sample(Table S1M). **(E)** Representative fluorescence microscopy images of mating pairs in indicated strains expressing mCherry-D4H. Images are at indicated time in time lapse started at t0 = 5 hours post nitrogen starvation in MSL-N liquid. White dashed outlines indicate mating pairs used for fusion neck width measurement. (**F**) Quantification of fusion neck width in strains shown in (E). N = 30 for each sample (Table S1N). (**G**) Quantification of cell size area in strains shown in (H). N=100 for each sample (Table S1O). (**H**) Representative light microscopy images of mating pairs that either successfully fused or failed to fuse after 48h at 25°C in indicated strains. White dashed outlines indicate the regions used for quantification.

These three genes displayed an oscillatory pattern in all evolving lines, with *clr5* oscillating in opposite direction than *ste11* and *sms1* (Figure S11B), consistent with the idea that these three genes are adaptive to fusion efficiency without *prm1*. Taken together, these data show that fusion efficiency in the evolved strain is driven by a rewired mating transcriptional program: *clr5* downregulation triggers *ste11* upregulation, which in turn elevates *sms1* expression. This cascade enhances partner encounters by promoting accelerated G1 arrest (*clr5*Δ and *ste11* OE) and more robust partner search (*sms1* OE), to boost Prm1-independent fusion attempts.

### Cellular Architecture and Membrane Dynamics Rewiring Drive Alternative Fusion Strategies

We asked whether rescue of the *prm1*Δ fusion defect in evolved strains also relies on increased intrinsic fusogenic capacity, in addition to cell cycle changes. We noticed that evolved strains exhibited morphological adaptations. Microscopy revealed substantial widening of the fusion neck in the e-Strain and the *clr5*Δ, *ste11* OE and *sms1* OE *prm1*Δ strains, with neck widths roughly 1.5 wider than in the *prm1*Δ ancestor strain (Figure 6E-F, Table S1N). While this morphological change may partly stem from the early formation of cell pairs, as confirmed by the observation that *cig2*Δ pairs also exhibit wider necks, the effect of *cig2*Δ was comparatively smaller, suggesting additional, possibly adaptive function in promoting fusion. We also observed that *clr5*Δ*, cig2*Δ, Ste11-OE, and evolved strains displayed significantly increased cell size— both in length and width—relative to wild-type and *prm1*Δ ancestors (Figure 6G-H, Table.S1O). Such morphological adaptations may facilitate fusion by creating a larger interface for membrane contact, and/or altering biophysical parameters like membrane contact curvature and tension^55,56^, thereby increasing the likelihood of a successful event.

To search for possible target genes of the *clr5-ste11-sms1* module in increasing the cells’ fusogenic properties, we compared the differentially expressed genes in *clr5*Δ, *ste11* OE, and *sms1* OE strains. This revealed a small set of commonly regulated genes (Figure S11C). Among them, Adg2^30^, a predicted GPI-anchored adhesion glycoprotein, is a candidate fusion facilitator. Indeed, overexpression of this gene in the *prm1*Δ ancestor, significantly increased fusion efficiency to ∼50% (Figure 7A-B, Table S1B), suggesting it is an effector contributing to alternative fusion mechanisms. This pinpoints *adg2* upregulation as a critical component of the evolved rescue pathway. Remarkably, deletion of *adg2* revealed its conditional essentiality in sexual reproduction: while dispensable for mating and fusion in the wild-type and *prm1*Δ ancestor, *adg2* was absolutely required in the e-Strain, where its loss resulted in 0% zygotes (Figure 7C, Table S1I.). Thus, *adg2* upregulation in the evolved strain acquired a different adaptive role to rescue fusion efficiency (Figure S11B). This gene was also oscillating in the other evolved lines (Figure S11B). Finally, to dissect the cellular mechanisms of rescue, we related formation of membrane invaginations, a hallmark of failed fusion in *prm1*Δ cells, with fusion outcome (Figure 1B, Figure 7D-E, Table S1P). In the *prm1*Δ ancestor strain, about 58% of pairs formed an invagination and the low percentage of fusion success occurred in cells without invagination, suggesting that invagination formation is a terminal, non-productive phenotype. In agreement with this, all rescue mutants dramatically boosted intrinsic fusion success in mating pairs that did not form an invagination but had more modest effects in cells with invagination. These results suggest that the main, but not only, zone of action of the adaptive fusion mechanism is in un-stalled productive mating events.

**Figure 7:**
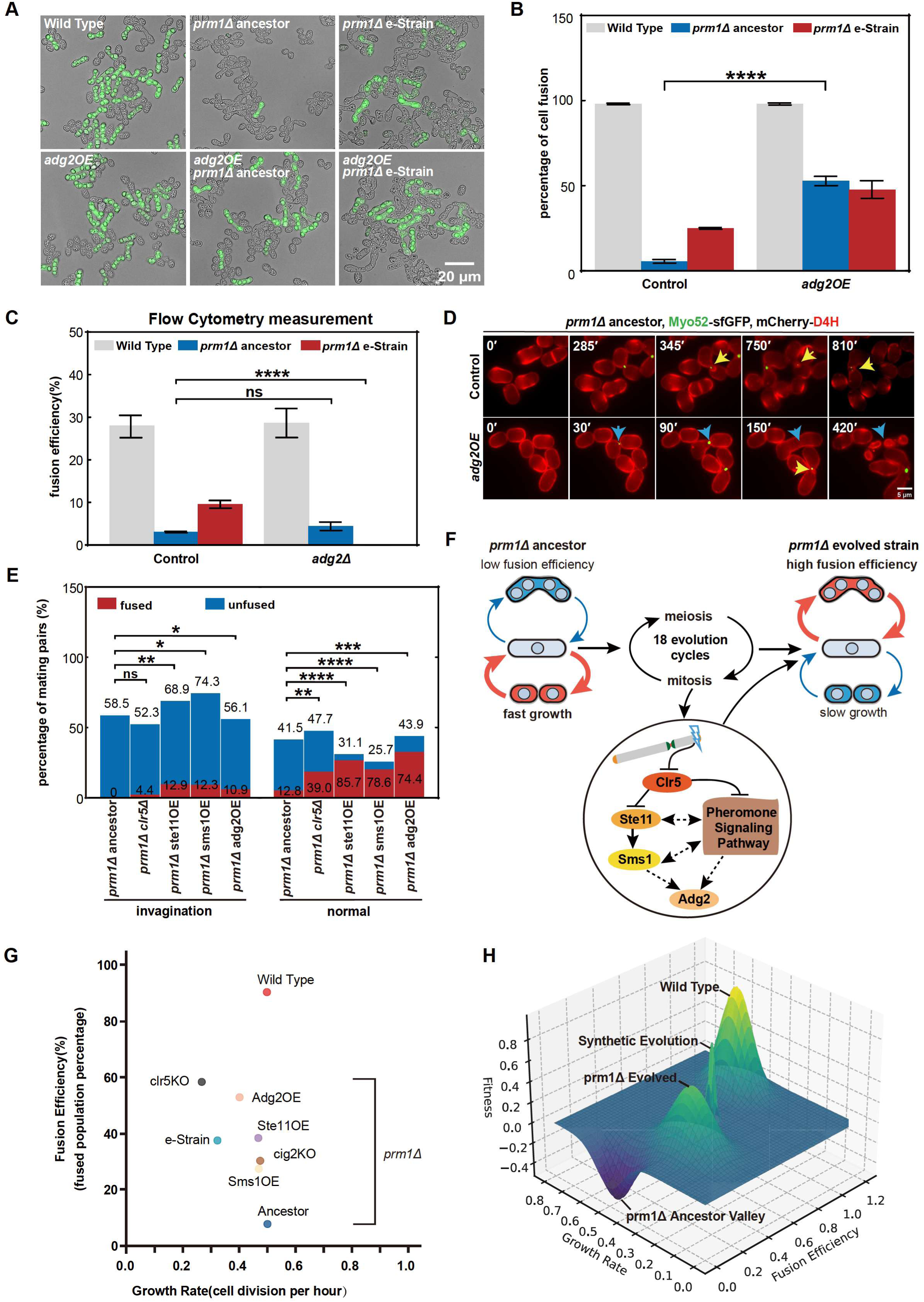
Adaptive rescue of fusion efficiency by *adg2* overexpression and fitness landscape. (**A**) Representative images showing cell fusion efficiency of *h90* WT, *prm1*Δ ancestor and *prm1*Δ e-Strain, with or without *adg2*OE, 48h post nitrogen starvation. (**B)** Quantification of cell fusion efficiency in strains as in (A). N ≥ 1000 in each sample (Table S1B). (**C**) Mating success as measured by flow cytometry in WT, *prm1*Δ ancestor, and *prm1*Δ e-Strain with or without *adg2*Δ. All strains express *p^meu31^*-sfGFP. N = 30,000 cells (Table S1I). (**D**) Representative fluorescence microscopy images of *prm1*Δ ancestor with or without *adg2*OE, expressing mCherry-D4H and Myo52-GFP during the fusion process. Yellow arrows indicate cell-cell contact sites with membrane invaginations. Blue arrows show a fusion site without membrane invagination, leading to successful fusion. In the time lapse, t0 corresponds to 5h post nitrogen starvation in MSL-N liquid. (**E**) Quantification of mating pairs with or without membrane invaginations, as shown in the representative images in (D) for *prm1*Δ ancestor, with or without *clr5*Δ, *ste11*OE, *sms1*OE, and *adg2*OE. N ≥ 86 for each sample (Table S1P). Fisher-exact test was used for statistical analysis. **(F)** Schematic working model of the molecular mechanism underlying the adaption of the e-Strain to *prm1* deletion. (**G**) Correlation analysis between fusion efficiency and growth rate in all engineered strains (Table S1I-J). (**H**) Proposed fitness landscape showing the WT peak, *prm1*Δ valley, *prm1*Δ e-Strain stable peak, and synthetic evolution peak.

In summary, the evolved strain utilizes sequential changes, including Clr5 downregulation, Ste11 upregulation, and increases in Sms1 and Adg2 expression, to achieve efficient fusion (Figure 7F). Our findings show that evolved rewiring in *prm1*Δ strains was initiated by a large genomic deletion, leading to epigenetic-mediated de-repression of mating regulators, which exploit G1 arrest timing, cell morphology, polarized growth and protein surface expression pathways to overcome – to a point – intrinsic fusion deficiencies. These multilayered adaptations converge to establish a partial Prm1-independent fusion mechanism under strong selection pressure.

### Synthetic Evolution Recovers Generalist Fitness by Bypassing Natural Trade-offs

Our study of the evolved *prm1*Δ strains showed that spontaneous adaptation in *prm1*Δ cells incurred a fundamental constraint: fusion efficiency increased at the cost of reduced mitotic growth. This trade-off positioned the evolved strain as a “fusion specialist” atop a narrow fitness peak characterized by high mating success but compromised proliferative capacity. This raises the question whether this trade-off also affects the individually engineered most impactful mutations, or whether it can be bypassed by rational design.

To probe this point, we measured both fusion efficiency and growth rate in *clr5*Δ, *cig2*Δ, Ste11-OE, Sms1-OE, and Adg2-OE in the *prm1*Δ ancestor background respectively. As shown earlier, all engineered strains exhibited substantial increases in fusion efficiency (20–50%), validating the individual contributions of these genetic modifications. However, while these strains exhibited some growth cost, only *clr5*Δ imposed a significant growth rate reduction, whereas the other engineered strains preserved proliferative capacity within ∼10% of the wild-type rate (Figure 7G, TableS1J).

These results suggest that while spontaneous evolution uncovers effective rescue pathways under strong selection, it may settle on local optima constrained by genomic architecture and regulatory entrenchment. In contrast, synthetic evolution, informed by evolutionary insight, enables direct construction of strains that escape natural trade-offs. By leveraging mutations like *adg2*-OE, which confer high fusion efficiency without strongly compromising growth, we successfully engineered a new fitness peak—one inaccessible to spontaneous evolution—but reachable through rational design (Figure 7H).

The fusion-growth trade-off maps onto a fitness landscape, where wild-type cells occupy a generalist peak, *prm1*Δ mutants fall into a deep fusion-deficient valley, and evolved strains climb toward a narrow specialist ridge. The synthetic peak, being extremely steep and narrow, indicates a theoretical state of high fitness but low robustness. Such a peak suggests that while the cells could achieve optimal performance at given scenario, they would be highly sensitive to environmental changes or mutations, making this state difficult to attain naturally.

## Discussion

Evolutionary biology is rich with elegant theories that capture fundamental dilemmas of life — from the trade-off between sexual and asexual reproduction^57–60^ to Waddington’s principle of genetic assimilation^61^. These ideas are widely regarded as true and supported by comparative and theoretical studies, yet direct evidence from experimental evolution has remained scarce. In this study, we provide such evidence in a single system. By evolving yeast through repeated asexual–sexual cycles, we support two long-standing predictions in action. First, we showed genetic assimilation: transient transcriptional oscillations that buffered fusion defects became stabilized when a subtelomeric deletion was acquired. Second, we observed a clear trade-off between sexual and asexual reproduction: lineages that improved mating efficiency did so at the expense of mitotic growth. Together, these findings show how experimental evolution can transform theoretical frameworks into mechanistic insight, linking regulatory plasticity, genomic change, and evolutionary trade-offs in a single trajectory.

### Trade-off between sexual and asexual reproduction

Our evolution experiment revealed an inverse correlation between fusion efficiency and growth rate, supported by both transcriptomic and phenotypic measurements. Across 12 evolving lines, oscillatory gene expression patterns tracked with alternating gains and losses in fusion and proliferation, mapping a zig-zag trajectory in fitness space. Genes driving these oscillations were enriched for not only pheromone signaling, metabolic processes, but also cell-cycle control, indicating that cells repeatedly shifted between transcriptional states specialized for fusion or for growth. Line 3 was exceptional: after early oscillations, it locked into a genetically encoded high-fusion, slow-growing phenotype. These findings mirror classical life-history trade-offs described across taxa, where investment in sexual reproduction reduces immediate proliferation, as seen in aphids alternating clonal and sexual phases, plants balancing seed set with clonal spread, or yeast and diatoms sacrificing growth for mating capacity^62–68^. Similar trade-off logic also applies to microbial stress evolution: yeast adapting to thermotolerance comes at the cost of growth at ancestral temperatures^69^, fitness gains in one stress reduce performance in others^70^, and populations alternating between salt and oxidative stress evolve asymmetric cross-protection rather than true generalism^14^.

Our system extends these principles by showing them in real time at the molecular level. Both transcriptomes were shaped by deletion of *prm1*, a gene that is normally dispensable for mitotic growth. The selection pressure for mating in absence of *prm1* not only forced cells to invest more in sexual reproduction but also compromised mitotic growth. Thus, although Prm1 itself acts only under mating conditions, its absence reshaped the set of accessible adaptive trajectories, illustrating how genes important only in inducible or differentiated states can profoundly influence both robustness and transcriptome-wide evolvability. Such genes, invisible under vegetative growth, emerge as hidden nodes of adaptive potential when selective pressures alternate.

### Escaping Constraints Through Synthetic Evolution

The sexual-asexual cycle trade-off likely constrained the evolution of *prm1*Δ cells, compared to robust wild-type cells, whose genome and transcriptome have been sculpted over millions of years to equip high performance in both sexual and asexual cycles. Indeed, line 3’s assimilation produced a specialist solution—robust fusion at the cost of reduced growth. Thus, while natural evolution restored significant levels of fusion, it remained bound by trade-offs.

By contrast, we found that rationally combining mutations could bypass these constraints. Overexpression of *adg2, ste11,* or *sms1*, or deletion of *cig2* in ancestral *prm1*Δ, each improved fusion efficiency – to higher rates than evolved line 3 – while maintaining near-wild-type growth. These engineered strains effectively “leapt” to higher fitness peaks that spontaneous evolution rarely accesses under standard conditions. This contrast highlights the difference between natural and designed adaptation. Natural evolution climbs local optima, constrained by genomic architecture and regulatory entrenchment. Synthetic evolution, informed by evolutionary insight, can bypass these limits to create genotypes that balance traits more effectively than nature’s solutions. Such engineered states may be fragile, occupying steep and narrow peaks less robust to environmental change. However, these observations suggests that sexual and asexual reproduction may not be in fundamental opposition, but that the observed trade-off is encoded in the regulatory networks that ensure these two processes occur at different times and conditions.

### Adaptive strategy: from transcriptional plasticity to genomic assimilation

Mechanistic dissection of e-Strain’s physiological adaptation suggests a two-stage process. Early in evolution, adaptation relied on transcriptional plasticity based bet-hedging strategies ^62^, in which phenotypic heterogeneity allows some cells to succeed under alternating asexual and sexual demands. Most lineages showed these plastic shifts without fixation, exemplifying genetic accommodation. Stability emerged only when a spontaneous 149-kb subtelomeric deletion appeared in cycle 9 of Line 3, exemplifying previously reported rare events^71^: genetic assimilation sensu Waddington^61,72^. This deletion extended subtelomeric silencing into the euchromatic *clr5* region. Interestingly, because *clr5* encodes a chromatin regulator, its repression created a recursive logic in which an epigenetic mechanism acted on an epigenetic regulator, triggering a cascade of secondary transcriptional changes that stabilized high fusion efficiency. In this way, genome rearrangement and epigenetic regulation acted to transform a transient response into a stably inherited phenotype. This evolutionary trajectory exemplifies how genomic and epigenomic changes can consolidate transcriptional plasticity into a heritable adaptive state.

### Rewiring of signaling pathways for adaptive fusion mechanisms

The rescue mechanisms we identified raise an important question: did evolved cells boost existing capacity for fusion, or did novel functions emerge^1,73,74^? Our data suggest both processes were at play. First, existing mating regulatory circuits were repurposed to repair fusion. Upregulation of the master transcription factor *ste11* accelerated entry into the mating program. The lengthening of G1 phase is by itself adaptive, as shown by deletion of the G1/S cyclin *cig2,* which is sufficient to increase fusion rates due to its prolonged G1 arrest. In parallel, the hemi-arrestin MAPK scaffold Sms1 promoted repeated mating attempts, increasing the chances of success even after initial failure. Collectively, these shifts did not restore Prm1’s fusogenic role but altered cell-cycle dynamics and pheromone-response behaviors to raise the overall probability of fusion. Second, adaptation exploited latent capacity. *Adg2*, a GPI-anchored surface protein dispensable in wild type or *prm1*Δ ancestor cells, became essential in the e-Strain. Further investigation will need to study its mode of action. Overexpression of *adg2* in the ancestral *prm1*Δ significantly boosted fusion efficiency, while its deletion abolished mating in the e-Strain. This highlights how genes that are nonessential in one background can acquire critical roles under new selective regimes. These findings underscore a principle: adaptation often proceeds not through novel invention but by rewiring existing circuitry and unmasking hidden redundancy. In our case, faster differentiation, prolonged mating readiness, and recruitment of a latent adhesion factor converged to establish a robust Prm1-independent fusion pathway.

### Limitations of the study

While our work provides evidence for the interplay of transcriptional plasticity, epigenetic silencing, and genetic assimilation in the adaptive rescue of fusion, several limitations should be acknowledged. First, our experimental evolution was restricted to 18 asexual–sexual cycles. Although this timeframe was sufficient to uncover a stable genomic assimilation event, longer-term evolution might reveal additional adaptive routes or alternative solutions. Second, transcriptomes and genomes were sequenced every third cycles. This sampling resolution was chosen to balance depth and throughput, but it necessarily reduces our ability to capture transient states. Third, although we functionally validated several key regulators (Clr5, Ste11, Sms1, and Adg2), the precise molecular mechanism of how Adg2 facilitates fusion remains to be dissected.

## Resource availability

### Lead contact

Further information and requests for resources and reagents should be directed to, and will be fulfilled by, the lead contact, Gaowen Liu.

### Materials availability

All key plasmids and the engineered *Schizosaccharomyces pombe* strains generated in this study, are available from the lead contact upon reasonable request

### Data and code availability

NGS, RNA-seq data generated in this study have been deposited in the National Center for Biotechnology Information (NCBI) Sequence Read Archive (SRA) under accession number PRJNA1334516 and PRJNA1242055 respectively. All code used in the processing and analysis of sequencing data and generation of figures are available from the lead contact upon request.

## Methods

### Yeast Strains and Media

#### Yeast Strains

All *S. pombe* strains used in this study are homothallic (h90) and are derivatives of the standard laboratory strain ySM1396 (yGL705). The decision to use a prototrophic h90 strain was to prevent any evolutionary biases that could arise from nutrient dependencies in auxotrophic strains. The ancestor strain for the evolution experiment was a *prm1*Δ mutant with an integrated sfGFP reporter driven by the *meu31* promoter at the *ura4* locus, selected with NATMX (nourseothricin resistance). This strain was designated as the starting point for the evolution experiments. For rescue experiments, genetic modifications were introduced into the wild-type like strain yGL450 (*h90, ura4::pMEU31-sfGFP-NATMX*), the *prm1*Δ ancestor yGL447, and the evolved e-Strain yGL636 (line 3, cycle 9). Detailed genotypes of constructed strains can be found in Table S1Q.

#### Yeast Media

YES Medium: Supports mitotic cell proliferation, maintaining vegetative growth and serves as a baseline for cell population analysis.

MSL-N Liquid Medium: Designed for synchronized meiosis induction, optimized for live-cell dynamic tracking, such as overnight time-lapse microscopy to monitor meiotic progression.

MSL-N agar Medium: Facilitates post-meiotic developmental analysis by providing a solid matrix for quantitative phenotyping, particularly for assessing fusion efficiency via colony morphology

#### Plasmid and Strain Construction

Unless specified otherwise, all overexpression and knockout constructs were generated using the In-Fusion HD Cloning Kit for seamless cloning via homologous recombination. Candidate gene sequences were obtained from PomBase to design PCR primers. For overexpression plasmids, the pGL51 plasmid served as the backbone, and constructs included the actin promoter (*pAct*), the candidate gene CDS, a C-terminal mCherry tag, and a hygromycin resistance cassette (hphMX), using the 5′ and 3′ UTRs of the *ade6* gene as homologous arms for integration^75^. For knockout constructs, the 5′ and 3′ UTRs of the target gene were used as homologous arms to replace the endogenous CDS with the hphMX selection marker, for detailed procedures, see the Suppkementary information 5.6. Prm1 tagging plasmids were constructed by inserting a fluorescent protein and a selectable marker between the *prm1* CDS and its 3’ UTR using In-Fusion cloning into a pGL42 backbone. Primers used for construction are listed in Table S1R. Strains were generated using the lithium acetate transformation method. Wild-type (yGL450), *prm1*Δ ancestor (yGL447), and *prm1*Δ evolved (yGL636) strains were grown to exponential phase. Plasmids were linearized by restriction enzyme digestion. Yeast cells were harvested, washed with 1× TE/LiAc buffer, and resuspended. Denatured salmon sperm DNA and linearized plasmid were added to the cell suspension, followed by incubation with a PEG/TE-LiAc solution for at least 3 hours at 30°C. After a heat shock at 42°C for 5 minutes, cells were recovered and plated on YES agar. Colonies were replica-plated onto selective media. Successful transformants were confirmed by PCR genotyping using genomic DNA extracted with an isolation buffer (250 mM LiAc, 1% SDS). Genotyping primers are listed in Table S1R. Some planned strains failed to be constructed and are marked in grey in Fig S9.

#### Fusion Efficiency Quantification

Fusion efficiency was quantified using three different methods depending on the experimental goal. In all cases, cells were first pre-cultured in MSL-N liquid medium for 5 hours. For all fusion efficiency analyses, statistical significance was assessed using the Student’s t-test. Data are presented as mean ± SD (*p < 0.05, **p < 0.01, ***p < 0.001, ****p < 0.0001).

1. Quantification by Microscopy (Plates): For absolute fusion efficiency, cells were spotted on MSL-N agar, incubated for 24 hours, and then scraped for imaging. Fused cell pairs (tetrads) and unfused cell pairs were counted from approximately 15 images per sample using ImageJ (v1.53). The fusion percentage was calculated as: % fusion = (Fused pairs / (Fused pairs + Unfused pairs)) × 100. This method was used to validate the fusion efficiency of the highest-performing strains from the 18th cycle (Figure S1 and Table S1B) and for strains from Line 3 across multiple cycles (Figure 4C and Table S1B).
2. Quantification by Flow Cytometry (FACS): For high-throughput screening, cells were mated on MSL-N agar for 24 hours, harvested into a 96-well plate, and analyzed on a BD FACSCelesta™ flow cytometer. The percentage of GFP-positive cells (expressed under pmeu31 promoter), detected in the FITC-A channel, was used as a measure of relative fusion efficiency. Data were analyzed using FlowJo software (v10.6.2). Because this method quantifies positive cells relative to all cells (even those that did not engage in mating), it only provides an indirect estimate of fusion efficiency with lower values than upon direct microscopy measurements. However, it clearly shows relative changes (Figure S1C-D). Statistical results are presented in Figure 1D, 3A-B, 7C, S6B, S9, S10B and Table S1I.
3. Quantification by Live-Cell Imaging (Slides): For dynamic monitoring, cells were transferred to MSL-N microscopy pads after the 5-hour liquid pre-incubation. Mating was observed over 24–48 hours on a Nikon Eclipse TI2-E microscope. Quantification was performed as described for plate-based microscopy. Statistical results are presented in Fig 5, Fig7 and Table S1B.

#### Growth Rate Measurement

To measure mitotic growth rate, all 72 evolved strains were revived and cultured in a 96-deep-well plate with 1 mL of liquid YE medium overnight. The next day, cultures were normalized to an OD of 0.02 in 200 µL of YE medium in a 96-well microtiter plate. Growth was monitored in a BioTek Epoch2 microplate reader at 30°C for 16 hours, with OD measured every 5 minutes with continuous shaking. The growth data were analyzed using a customized R script (Figure 3A, Table S1J).

#### Genomic Sequence Analyses

DNA was extracted from 72 evolved strains, plus wild-type and ancestor controls, using a standard phenol-chloroform protocol. Sequencing libraries were prepared by the University of Lausanne Genomic Technologies Facility and sequenced on an Illumina Hi-seq platform. Raw sequencing data were processed using Varathon (v1.0.0) to detect single nucleotide variants (SNVs), INDELs, structural variants (SVs), and copy number variations (CNVs). High-confidence variants were retained (QUAL > 30). Functional Annotation: Variants were functionally annotated using SnpEff (v5.1d) with the *S. pombe* reference genome (PomBase release 2024.08). Variant Visualization: Circos plots were generated with the circlize R package (v0.4.15) to visualize SNP density, INDEL distribution, and CNVs across the genome (Figure 4A outer ring, Table S1H). Coverage profiles were visualized with ggplot2 (Figure 4B). Variant burden was visualized using violin plots and stacked bar charts created with ggplot2 (Figure S5A-B). Variant impact analysis is shown in a heatmap (Figure S5C).

To identify large-scale structural variants, the e-strain and wild-type genomes were analyzed using long-read sequencing on the Oxford Nanopore Technologies (ONT) platform. The e-strain genome was de novo assembled using canu (v2.2) within the LRSDAY pipeline. Whole-genome alignment between the wild-type and e-strain assemblies was performed using nucmer (v3.1) to identify syntenic blocks. Structural variants were annotated using SyRI (v1.7.0), with translocation events being manually curated to avoid artifacts from telomeric sequence homology (Figure S6A). Deletion junctions were confirmed by Sanger sequencing (Figure S6A). Sequencing data are deposited in NCBI SRA database under accession number PRJNA1242055.

#### Transcriptomic Analysis

For each of the 72 evolved strains plus controls, three biological replicates were collected for RNA-sequencing. Cells were cultured to mid-log phase, transferred to MSL-N medium for 5 hours, harvested, and rapidly frozen in liquid nitrogen. RNA extraction and sequencing (Illumina NovaSeq 6000, 150 bp paired-end) were performed by Novogene. Raw reads were assessed with FastQC (v1.4.0) and trimmed with Trim Galore (v0.6.10). Cleaned reads were aligned to the reference CDS genome (PomBase release 2024.08.20) and quantified using Salmon (v1.4.0) to generate Transcripts-Per-Million (TPM) values.Batch Correction: To correct for technical variability across six sequencing runs, a unified TPM expression matrix was log -transformed (log (TPM + 1)) and subjected to batch correction using the ComBat method from the sva R package (v3.48.0). Differential Expression Analysis: Differential expression was analyzed separately for 0h and 5h timepoints using the limma R package (v3.56.2) on the batch-corrected, log-transformed data. Pairwise comparisons between each evolved strain and the ancestor were performed, with differentially expressed genes identified using an adjusted p-value < 0.05 and a |log fold change| ≥ 1 (stringent) or ≥ 0.5 (exploratory). Various tools were used for downstream analysis, including ClusterGVis for fuzzy c-means clustering and heatmap visualization (Figure S2-3), clusterProfiler for Gene Ontology (GO) enrichment, and the bulkPseudotime R package for pseudotemporal ordering of gene expression (Figure 2B). Integration of transcriptomic and phenotypic data was performed using t-SNE (Fig 3B, Table S1F) and linear regression (Figure 3A, Table S1J). Candidate genes were identified using multiple methods including OptICA, cMonkey, and UpSetR (Figure S8A-C).

#### RT-qPCR Analysis

To validate expression levels of specific genes (*clr5*, *ste11*, *sms1*), RT-qPCR was performed. Total RNA was extracted from cells grown in YES and MSL-N for 5 hours using the Yeast Total RNA Extraction Kit. RNA quality was assessed, and 2 µg was used for first-strand cDNA synthesis with the All-In-One 5× RT MasterMix with gDNA Removal Kit. Quantitative PCR was performed on a qTOWER³ real-time PCR thermal cycler using PerfectStart® Green qPCR SuperMix. Relative gene expression was calculated using the 2-ΔΔCt method, with *act1* serving as the endogenous control (Table S1K).

#### Microscopy and Image Analysis

Time-Lapse Microscopy was performed on a Nikon Eclipse TI2-E inverted fluorescence microscope with a 100× objective. Cells were spotted on 2% MSL-N agarose pads and sealed with VALAP. Images were typically captured every 15 minutes for approximately 20 hours.

Cell area and length were quantified using Cellpose for automated segmentation of images. Segmented regions of interest (ROIs) were imported into ImageJ for measurement, with results presented in Figure 6G-H. The length of the fusion neck in early-stage mating pairs was measured from time-lapse videos using the ‘Straight’ tool in ImageJ (v1.53). At least thirty mating pairs were measured per sample, with results in Figure 6E-F. Time-lapse videos were inspected frame-by-frame to score mating behaviors, such as cells attempting to mate with multiple partners (Figure 6A-B, and Supplementary Table S1M). The number of fluorescent foci per cell was quantified using the multi-point tool in ImageJ (Figure 6C-D, Table S1M). The membrane invagination phenotype in *prm1*Δ cells was imaged by crossing *prm1*Δ D4H-mCherry with *prm1*Δ D4H-sfGFP strains (Figure 1B). The presence or absence of invagination in fused and unfused cells was scored to quantify the phenotype in rescue strains (Figure 7D-E, Table S1P). For all statistical analyses of microscopy and image data, the Student’s t-test was used. Data are presented as mean ± SD (*p < 0.05, **p < 0.01, ***p < 0.001, ****p < 0.0001).

#### Yeast One-Hybrid (Y1H) Assay

To test for DNA-protein interactions, specifically between the transcription factor Ste11 and the *Sms1* promoter, a Y1H assay was performed using a kit from Coolaber Bioscience (results in Supplementary Figure S10C). Promoter fragments of *Sms1* were ligated into the pAbAi bait plasmid, and the *Ste11* CDS was ligated into the pGADT7 prey plasmid. The linearized pBait-AbAi plasmid was transformed into the Y1HGold yeast strain and integrated into the genome. The optimal concentration of the selection agent Aureobasidin A (AbA) was determined to screen for background expression. The prey plasmid was then transformed into the competent bait strain, and interactions were validated by assessing growth on selective media (SD/-Leu) containing the predetermined AbA concentration.

#### Cell Cycle Analysis

To analyze cell cycle profiles, DNA content was measured by flow cytometry after propidium iodide (PI) staining. Log-phase cells were harvested and fixed in 70% ethanol. Cells were sequentially treated with RNase A and proteinase K before being resuspended in sodium citrate solution and stained with PI. DNA content was analyzed on a flow cytometer, and the data were processed using FlowJo software to gate G1 and G2 populations and calculate their frequencies, while excluding doublets. The results of this experiment are shown in Figures 5G and 5H, and Supplementary Table S1L.

### Author Contributions

GL and SGM conceived the initial idea; GL supervised all aspects of the project; SGM supervised evolution and cell biology part of the project; GL, YC, SL, YHW and LB performed experiments; LB, CYL and GL performed genomic and computational aspects of the project; XLF discovered *clr5* expression change; GL, LB, YC and CYL analyzed the data; GL wrote the manuscript with feedback from SGM; GL, YC and LB prepared figures. All authors read and agreed to its content.

## Acknowledgments

Authors thank Lilin Du for reagents; Jun Peng, Manying He, Yuan Miao, Yuanyuan Liu and Yiqing Zhang for technical assistance; Jing Li, Jiaxing Yue, Haiyun Gan, and Qing Ma for critically reading the manuscript; and Laura Merlin, Aleksandar Vještica, M-Henar Valdivieso, Teng Wang and Xiao Yi for useful discussions. GL was supported by HFSP long-term fellowship LT-001727/2017-L. This work was supported by Swiss National Science Foundation grant #155944 and European Research Council grant SexYeast to SGM. This research was also supported by grants from the National Natural Science Foundation of China (Grant No. 32300496) to GL, Guangdong Provincial Key Laboratory of Synthetic Genomics (2023B1212060054) and Shenzhen Key Laboratory of Synthetic Genomics (ZDSYS201802061806209). All authors declare that no conflicts of interest exist.

---

**Supplemental Figure 1.**
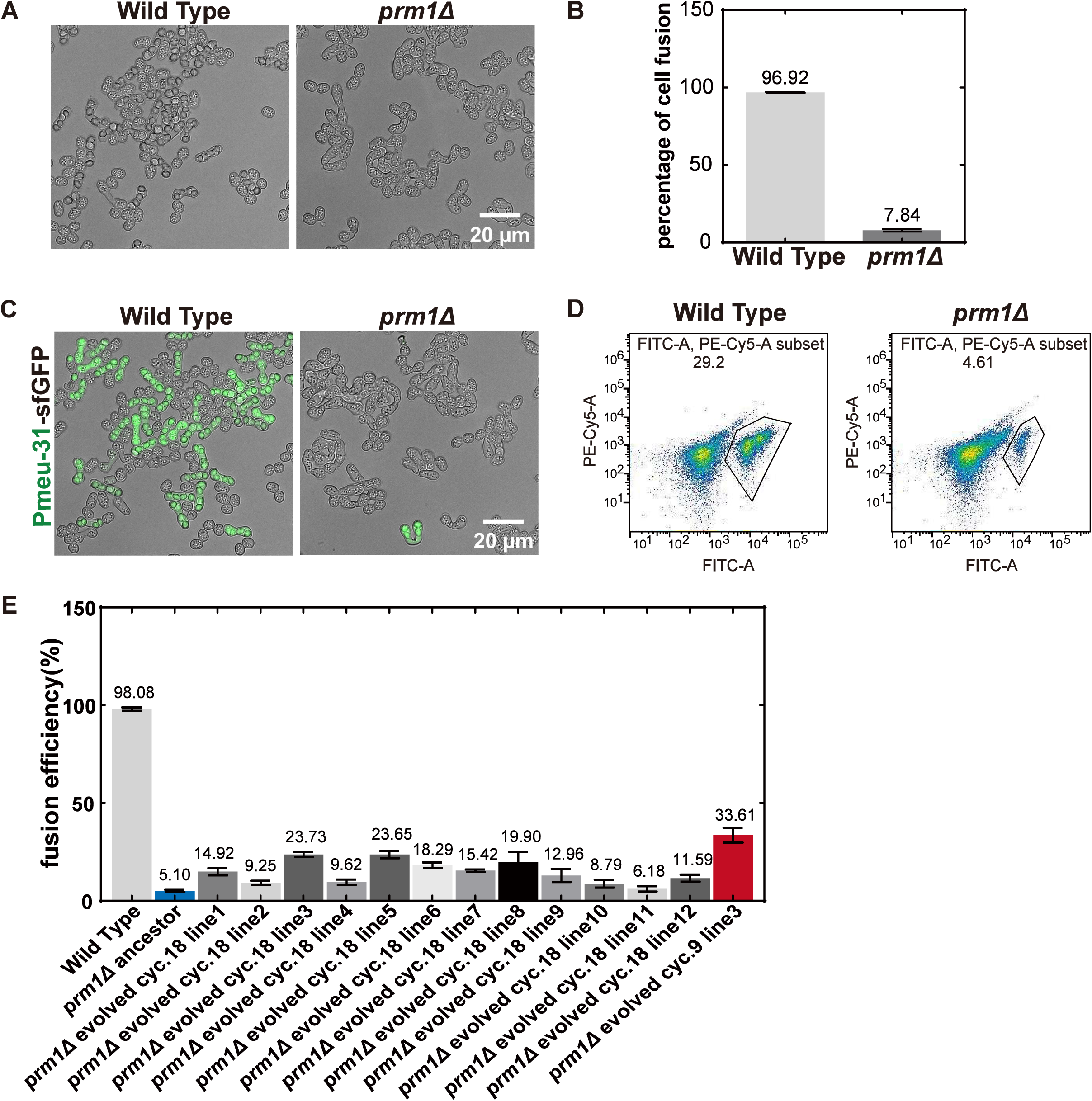
Prm1 deletion causes severe fusion defect and *prm1Δ* ancestors strain construction, related to Figure 1. (**A**) Light microscopy representative mating pairs in wild type and *prm1Δ* ancestor. Scale bar = 20 μm. **(B**) Quantification of fusion efficiency of the microscopic picture shown in (A). N>396 for each sample (Table S1B). (**C**) Fluorescence microscopy representative mating pairs in wild type and *prm1Δ* ancestor expressing *Pmeu31*-GFP. Scale bar = 20μm. (**D**) Flow cytometry readouts of *prm1Δ* ancestor expressing *Pmeu31*-GFP. The gated population indicates relative fusion efficiency. **(E)** Quantification of fusion efficiency of wild type, *prm1Δ* ancestor, *and prm1Δ* evolved 18^th^ cycle strains. The e-Strain (line 3, cycle 9) is shown in red. Cells were analyzed after 48 hours of nitrogen starvation to induce mating and fusion (Table S1B).

**Supplemental Figure 2.**
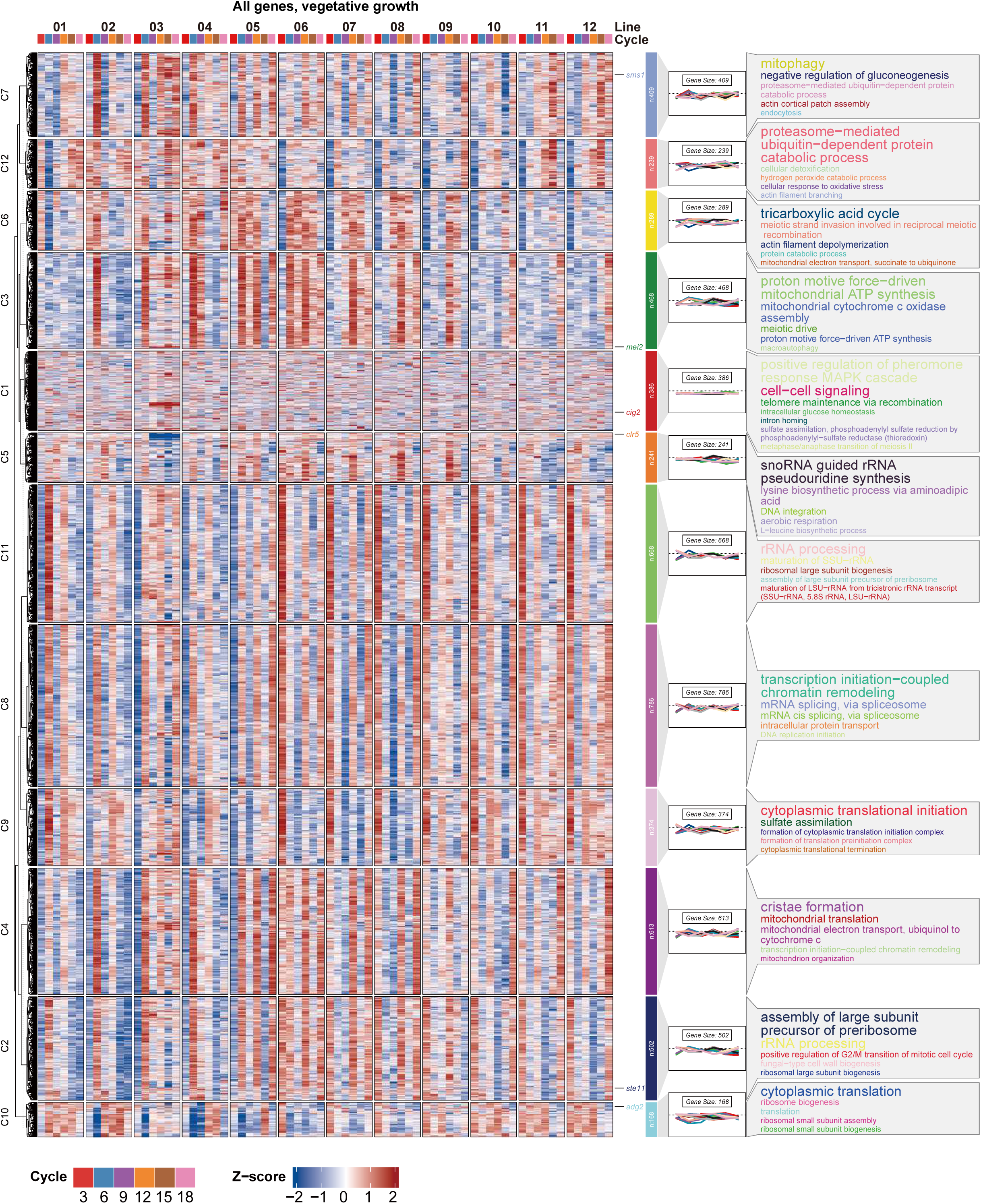
Gene expression level trajectory during vegetative growth, related to Figure 2. Complete clustering of all gene expression level pattern along the evolutionary timeline across all samples during vegetative growth (t0 time point). Right panel: GO enrichment, gene set ratio of enriched GO term (Table. S1C).

**Supplemental Figure 3.**
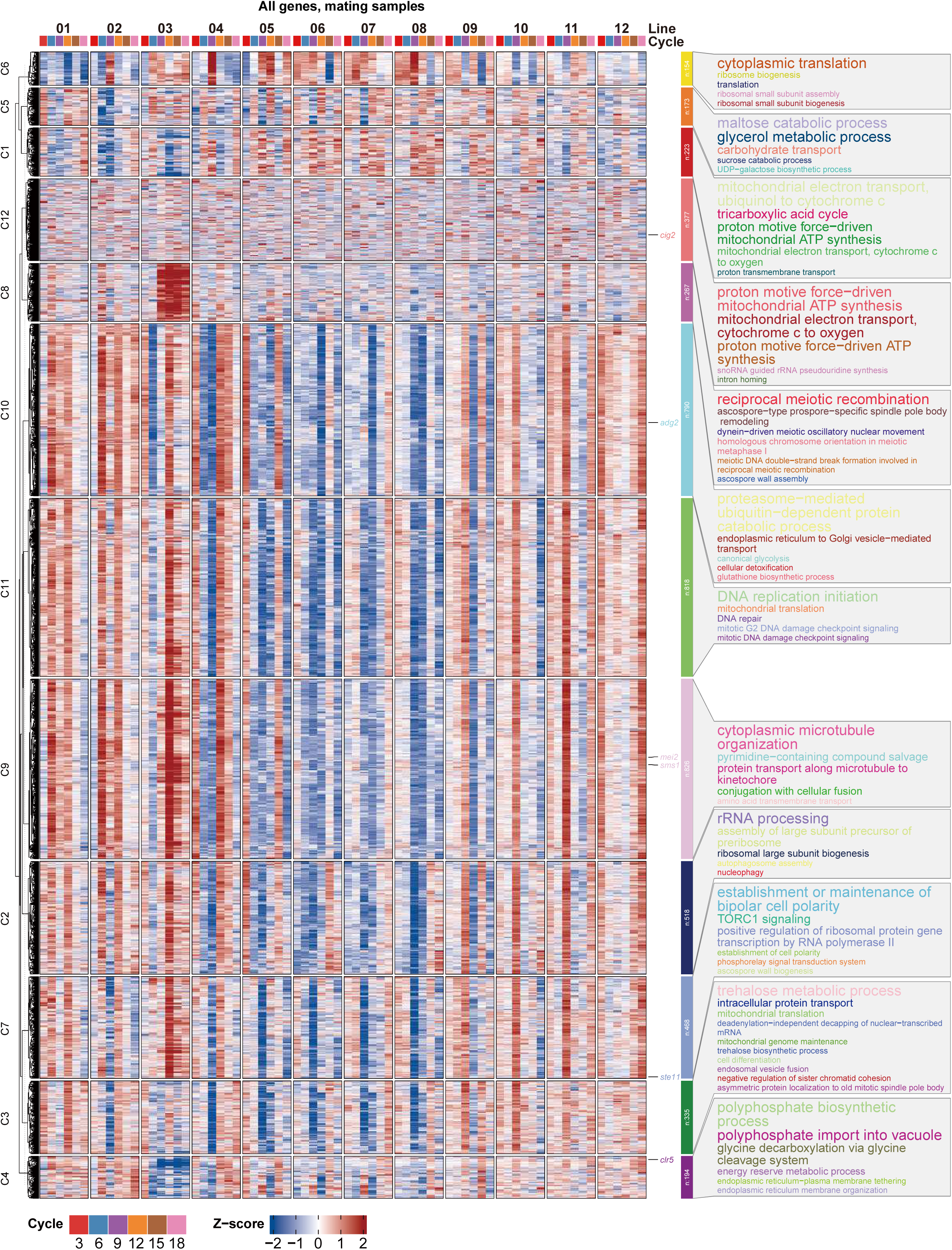
Gene expression level trajectory during 5h post-nitrogen starvation, related to Figure 2. Complete clustering of all gene expression level pattern along the evolutionary timeline across all samples at 5h under nitrogen starvation. Right panel: GO enrichment, gene set ratio of enriched GO term (Table. S1D).

**Supplemental Figure 4.**
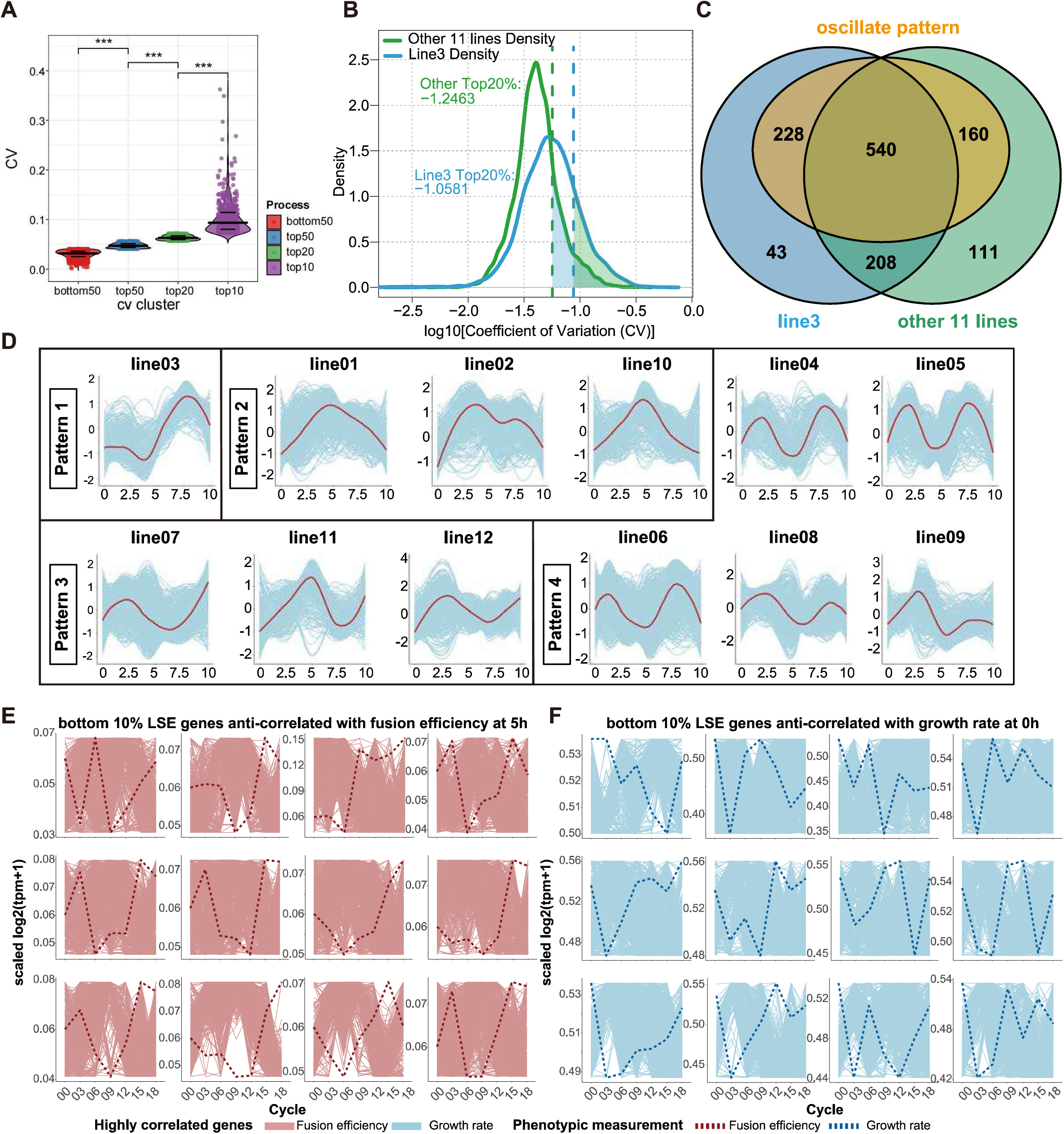
Integrative analysis of transcriptomic profile and phenotypes of fusion efficiency and growth rate, related to Figure 2 and Figure 3. (**A**) Distribution of the coefficient of variation (CV) across all protein-coding genes following control subtraction, stratified into four expression-variability clusters (bottom50: >50%, 2548 genes; top50: 50%-20%, 1528 genes; top20: 20%-10%, 509 genes; top10: 10%, 510 genes). (**B**) Coefficient of variation (CV) distributions for line3 (blue) and aggregate non-line3 variants (green) following control normalization. The corresponding dashed lines indicate the positions of the top 20% of CV values for each distribution, highlighting regions of significant variability in gene expression (Table S1E). (**C**) Venn diagram of the unique gene sets and overlap between line 3 (blue) and the remaining lines (green), for genes within the top 20% of CV. Genes with oscillatory pattern (928 genes) are annotated in orange. (**D**) Expression level (blue profiles) and average pattern (red lines) of the 928 oscillatory genes along pseudo-time in all evolving lines. (**E**) Expression trajectories (red lines) of the 10% of genes in each line least correlated with fusion efficiency (red dashed line, relationship measured by LSE; Table S1G,I) from RNAseq data 5h post nitrogen starvation (log2(TPM+1), min-max scaled per lineage); X axis = evolutionary cycles; Y axis = normalized scale of each line. X axis represents evolutionary cycles; Y axis represents normalized scale for each line. (**F**) Expression trajectories (blue lines) of the 10% of genes least correlated with growth rate (blue dashed line, relationship measured by LSE; Table S1G,J) from RNAseq data during vegetative growth (T0; log2(TPM+1), min-max scaled per lineage); X axis = evolutionary cycles; Y axis = normalized scale of each line.

**Supplemental Figure 5.**
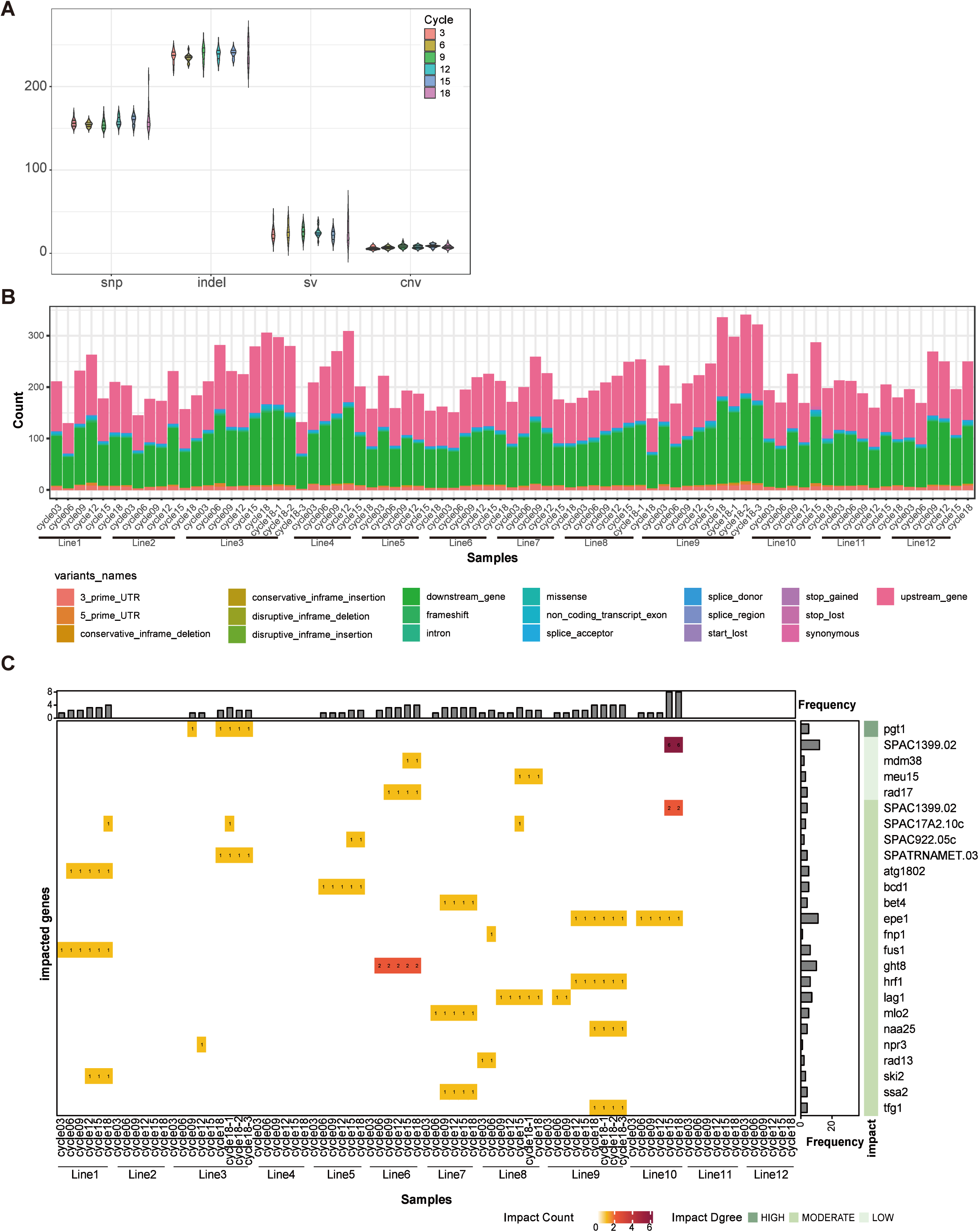
Summary of Genomic Variants Across All Strains, related to Figure 4. (**A**) Landscape of genomic variation stratified by cycle. (**B**) Stacked bar plots depicting variant counts across all experimental samples, stratified by SnpEff-defined functional categories (Table S1H). (**C**) Functional impact stratification of genomic variants via SnpEff annotation across strains compared to the ancestor strains. Heatmap (main panel) quantifies counts of high-, moderate-, and low-impact variants in 25 genes across all the evolved strains, with color intensity scaled to variant frequency (Table S1H). The color ranging from red to yellow suggests the number of mutations found on that gene. The top bar plot summarizes total impact counts per evolved strain. The right bar plot aggregates gene-specific burdens across all the evolved strain.

**Supplemental Figure 6.**
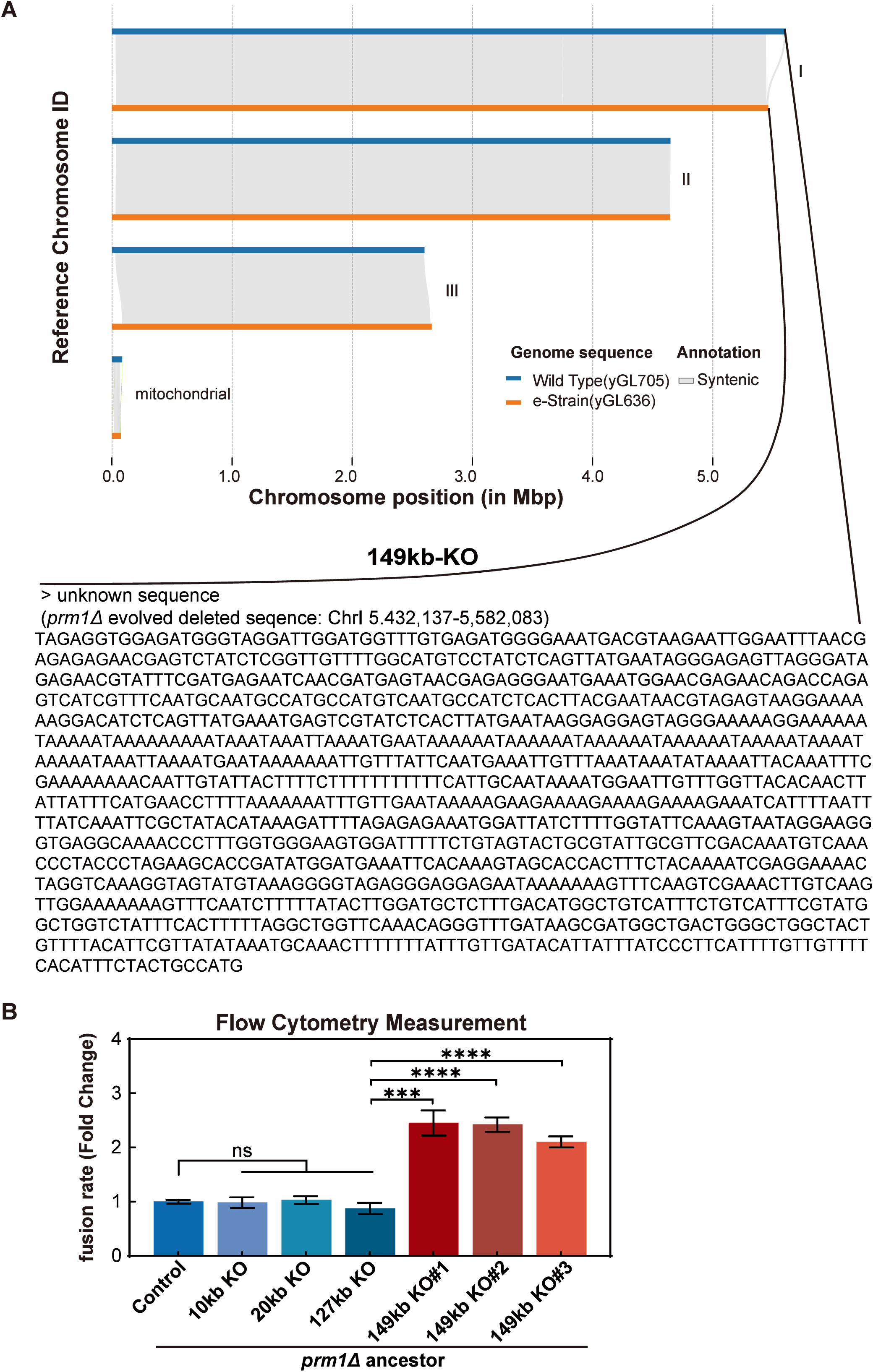
Architecture of Telomere-Proximal Genomic Remodeling in Evolved and Engineered Strains, related to Figure 4. (**A**) Integrated structural variant validation via LRS assembly, and Sanger sequence of the unknown region. Upper panel: strain-specific structural variations at the Chromosome I telomere region, combining long-read sequencing (LRS)-assembled genome alignments between wild type and e-Strain. Lower panel: Sanger sequencing-confirmed unknown Chromosome I -terminal fragments. (**B**) Fusion efficiency was measured by flow cytometry in *prm1*Δ ancestor backgrounds, along with derivative strains carrying 10 kb, 20 kb, 127 kb or 149 kb knockout (KO) (Table S1I).

**Supplemental Figure 7.**
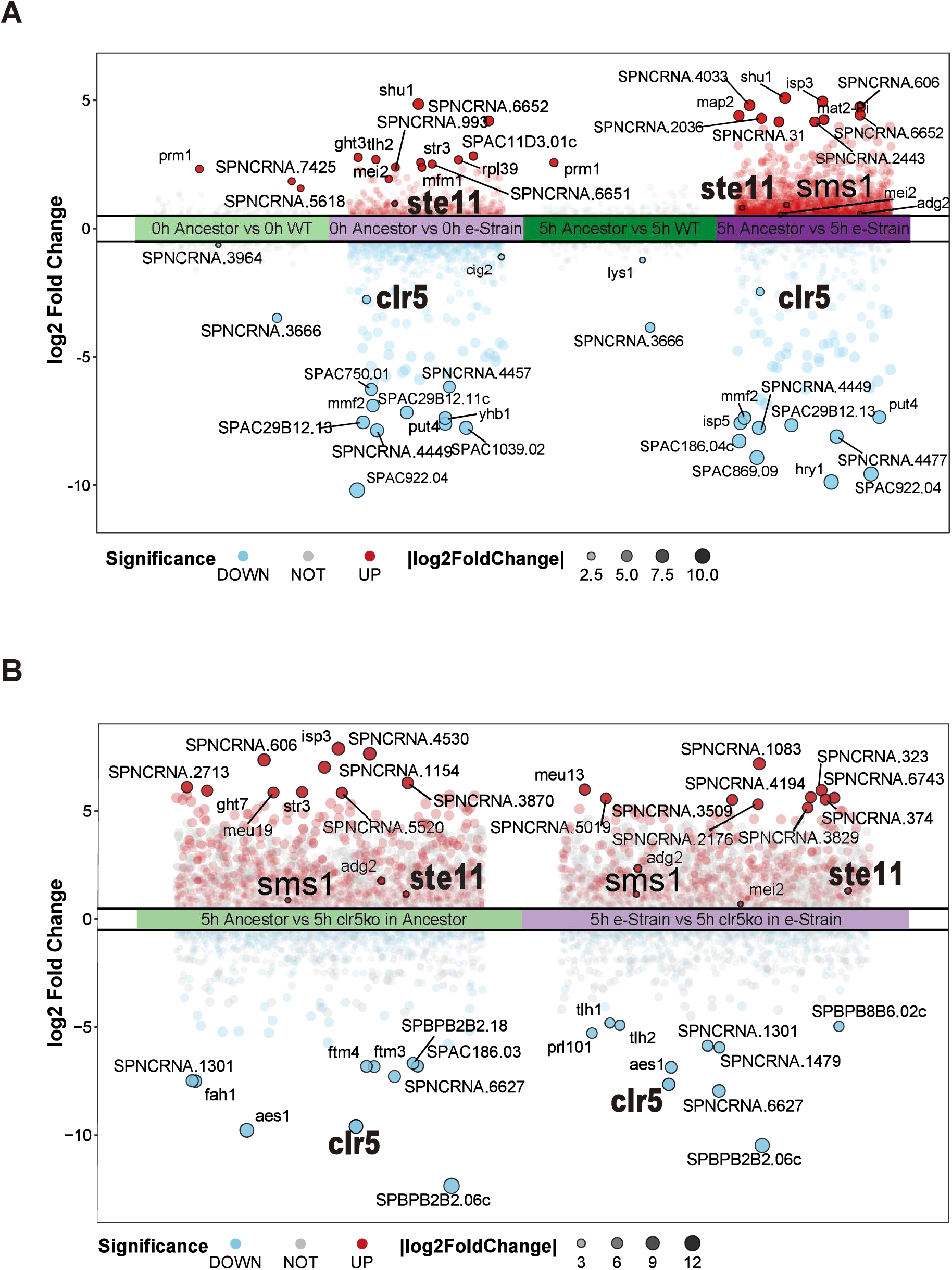
Integrated Analysis of Gene Regulation and Transcriptomic Profiles in Evolved Strains and Engineered *clr5Δ* Strains, related to Figure 5-6. (**A**) Comparative transcriptomic analysis of *prm1Δ* e-Strain versus *prm1Δ* ancestor and *prm1Δ* e-Strain versus wild type at both 0h and 5h. The central strip indicates the significance threshold of ±0.5 log2 fold change. *ste11, sms1 clr5* are highlighted in larger font. Many candidate genes for later screen for *prm1Δ* fusion deficiency rescue were identified from this analysis (Fig. S9, Table S1I). (**B**) Comparative transcriptomic analysis of *clr5Δ prm1Δ* ancestor versus *prm1Δ* ancestor and *clr5Δ prm1Δ* e-Strain versus e-Strain. The central strip indicates the significance threshold of ±0.5 log2 fold change. *ste11, sms1 clr5* are highlighted in larger font.

**Supplemental Figure 8.**
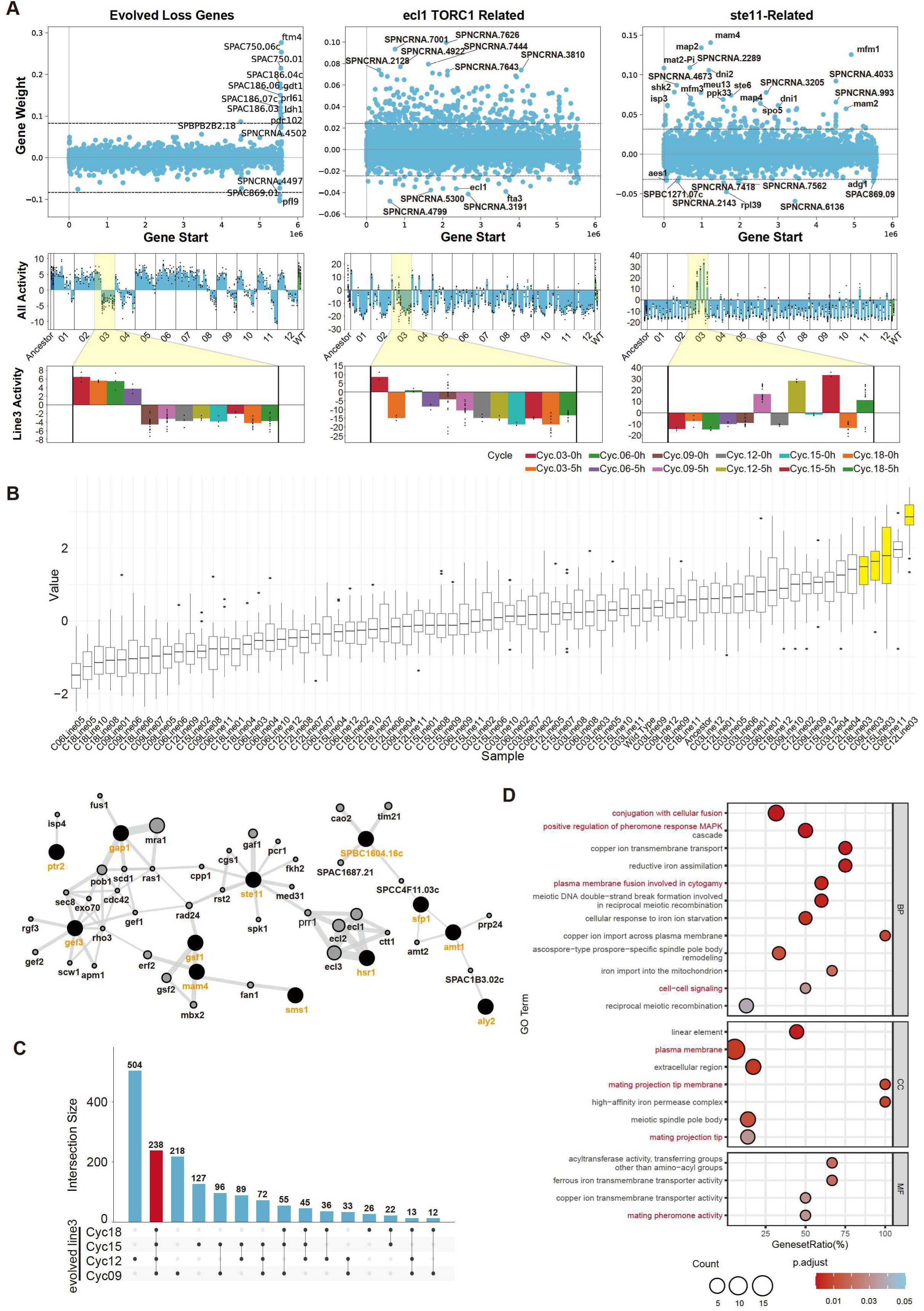
Candidate genes identification through machine learning methods, related to Figure 5-6. (**A**) Identify regulatory network in evolved strains by OptICA with three gene modules (Evolved Loss Genes, ecf1-TORC1 Related, and ste11-Related). Each module panel integrating gene weight scatter plots (upper: genomic position vs. gene weight) and activity distribution profiles (center: total activity, lower: line 3-specific activity). (**B**) Upper panel: normalized and ranked scores of gene modules highly corelated with samples of line 3 after cycle 9, which have high fusion efficiency profiles. Lower panel: the GeneMANIA-generated interaction network integrates co-expression (gray edges) of 12 module genes identified from upper panel (black nodes). The grey dots are not presented in the module identified, it is shown here to display the full picture of module gene interaction status. (**C**) Upset Analysis Identifies Conserved 238 Upregulated Genes (red bar) in Line3 Post-Cycle9 Strains. (**D**) Functional Enrichment of Conserved Post-Cycle9 Upregulated Genes. MAPK/membrane-related go terms were highlighted with red.

**Supplemental Figure 9:**
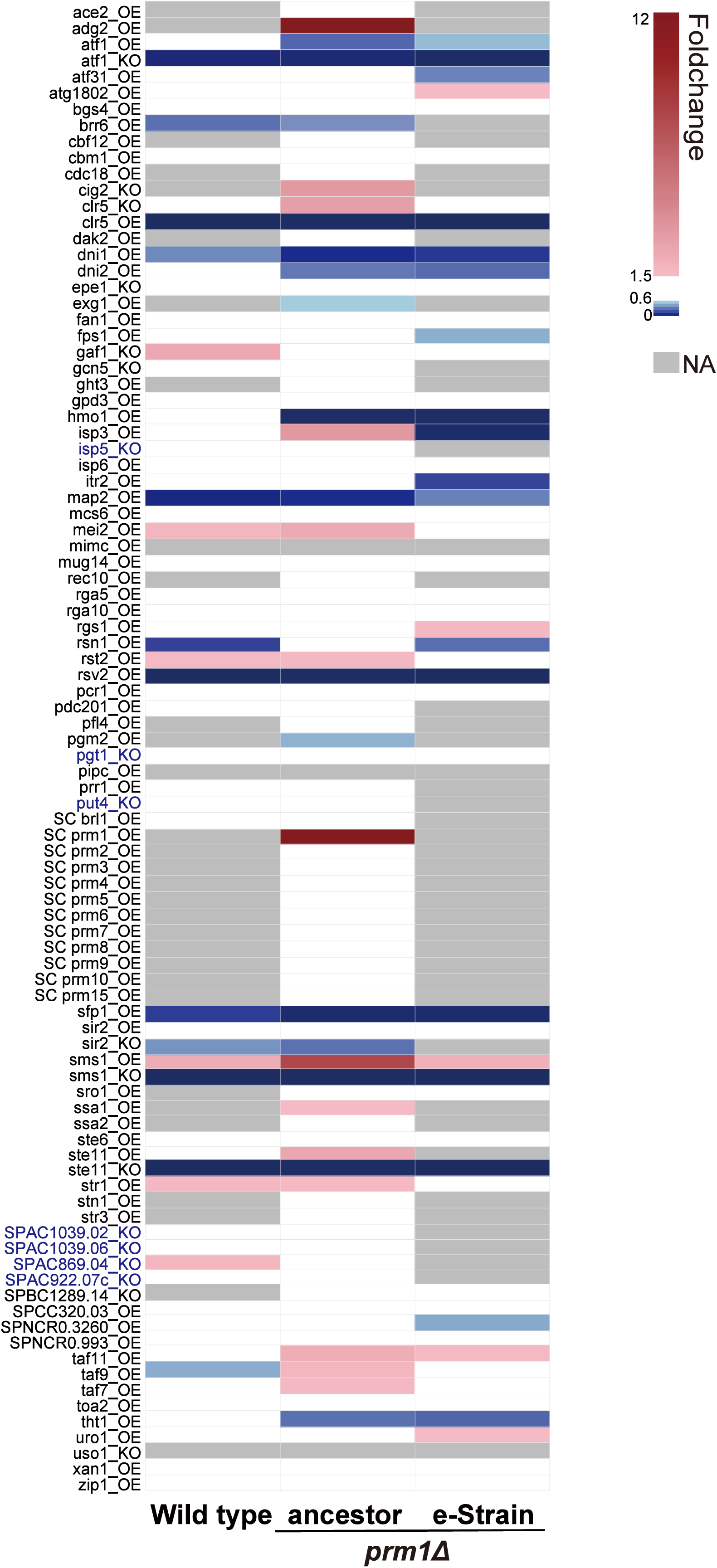
Screening results of candidate fusion efficiency increasing genes, related to Figure 5-6. Heatmap of fold change in fusion efficiency (performed in triplicate) relative to the *prm1Δ* ancestor control for all 98 candidate genes tested. The gene names labeled on the left are the genes being modified. *SC_prm1* means *S. pombe prm1* ortholog in *Saccharomyces cerevisiae.* Other *SC_prm\** encode membrane proteins involved in mating in *Saccharomyces cerevisiae*, which were identified together with *prm1*^1^. OE means that gene was overexpressed; KO means that gene was knocked out. The modification decisions of OE or KO were based on the gene expression change comparing higher fusion efficiency strains (both evolved and engineered strains’ transcriptome data) to the *prm1Δ* ancestors. For some candidate gene, the wild type, *prm1Δ* ancestor and e-Strain have all been engineered with the same OE/KO modification per row to compare the fusion efficiency changes while for others we only engineered the OE/KO in the *prm1Δ* ancestor background. The color code shown on the right represents the fusion efficiency fold change. Gray means that the strain was not engineered. The raw fusion efficiency data were acquired from flow cytometry GFP population readouts (Table S1I).

**Supplemental Figure 10:**
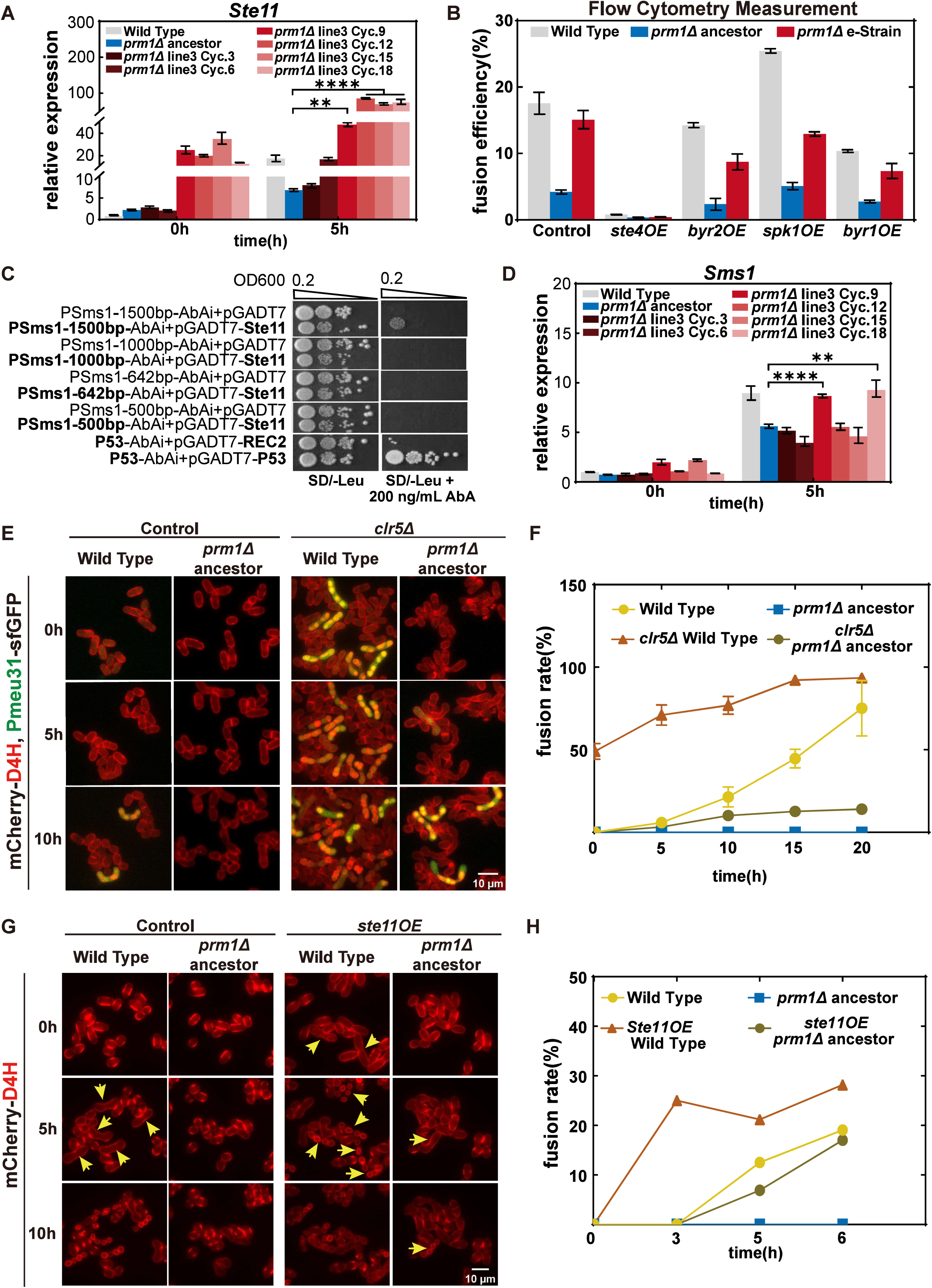
*clr5* deletion and *ste11* overexpression increase fusion efficiency by promoting a faster G1 arrest, related to Figure 5. (**A**) Relative expression levels of *ste11* in the line3 evolved strain measured at 0 hours and 5 hours by qPCR (Table S1K). (**B**) Flow cytometry measurement of fusion efficiency of overexpression of MAPK pathway components (Ste4, Byr2, Spk1, Byr1) in wild-type (WT), *prm1Δ* ancestor, and e-Strain background. All strains express *Pmeu31*-sfGFP. Flow cytometry analysis was performed by collecting fluorescence signals from a total of 30,000 cells (Table S1I). (**C**) Yeast one-hybrid assay to validate the interaction between Ste11 and the *sms1* promoter. The images show the growth of strains co-transformed with the Ste11-pGADT7 plasmid and different lengths of the *sms1* promoter in the pAbAi plasmid, on plates with or without 200 ng/ml AbA. Growth was assessed through serial dilution, starting with an initial OD = 0.2. (**D**) Relative expression levels of *sms1* in line3 evolved strain measured at 0 hours and 5 hours (Table S1K). (**E**) Representative fluorescence microscopy images of early mating events in *h90* strains. Shown are *clr5Δ* in the wild-type and *prm1Δ* ancestor backgrounds, along with wild-type and *prm1Δ* ancestor controls. All strains express *Pmeu31*-sfGFP and mCherry-D4H to monitor fusion rates during early cell fusion. Images were captured starting at the 0-minute time point, which corresponds to 5 hours of nitrogen starvation after mating induction in MSL-N liquid. Scale bar = 10 μm. (**F**) Quantification of the fusion rate as shown in the representative images in (E) at 0, 5, 10, 15, 20 hours (Table S1B). (**G**) Representative fluorescence microscopy images of early mating events in *h90* strains. Shown are Ste11OE in the wild-type and *prm1Δ* ancestor backgrounds, along with wild-type and *prm1Δ* ancestor controls. All strains express *Pmeu31*-sfGFP to monitor fusion efficiency during early cell fusion. Images were captured starting at the 0-minute time point, which corresponds to 5 hours of nitrogen starvation after mating induction in MSL-N liquid. Scale bar = 10 μm. (**H**) Quantification of the fusion rate as shown in the representative images in (G) at 0, 3, 5, 6 hours (Table S1B).

**Supplemental Figure 11:**
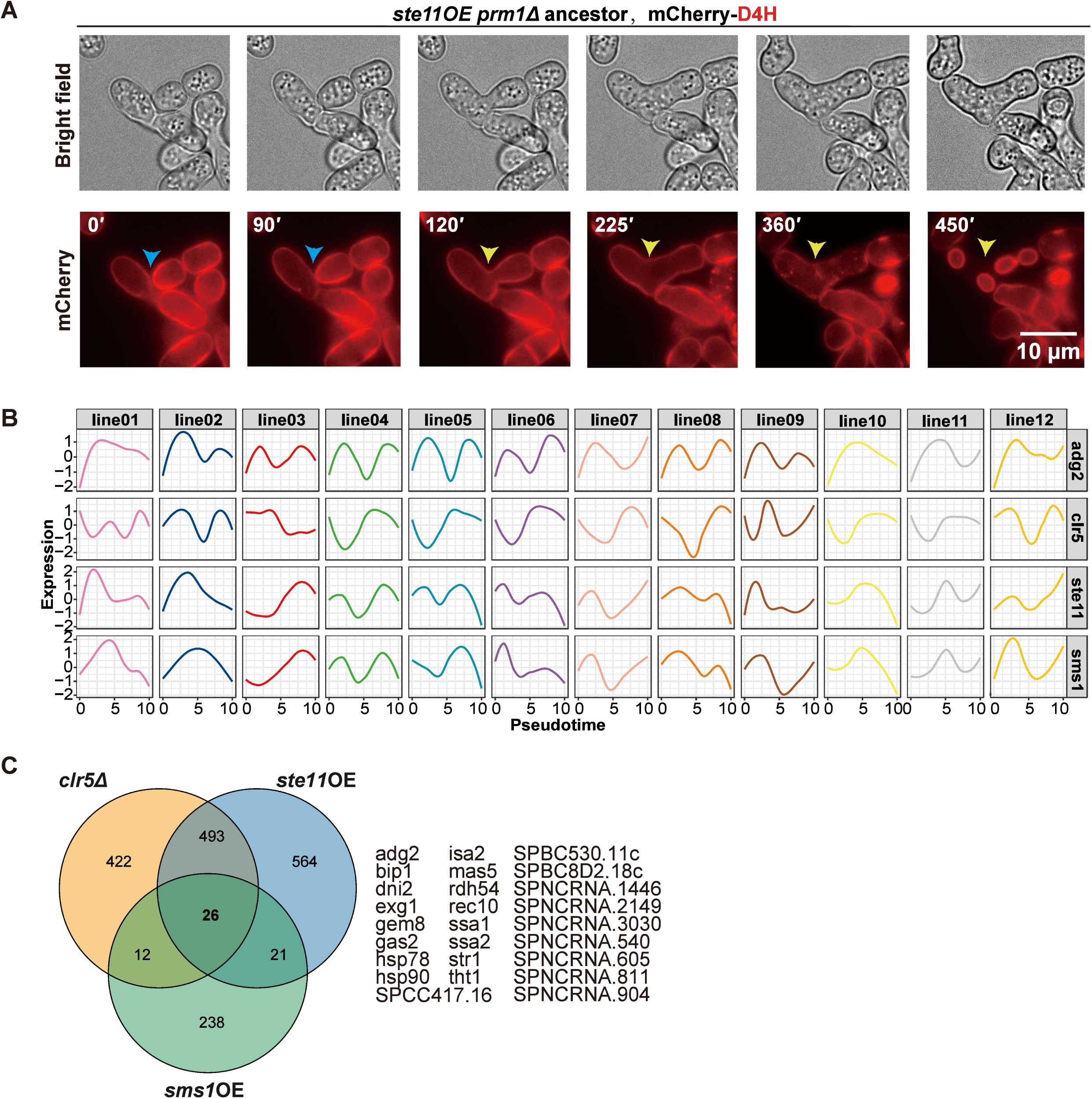
Influence of *ste11* on fusion efficiency and identification of downstream targets, related to Figure 6 and Figure 7. **(A**) Representative light and fluorescence microscopy images of *ste11OE* in the *prm1Δ* ancestor background, expressing mCherry-D4H during the fusion process, showing an example of fusion attempt with two partners in succession. The first attempt is unsuccessful (blue arrowhead). The second one is successful (yellow arrowhead). Images were captured starting at the 0-minute time point, which corresponds to 5 hours of nitrogen starvation after mating induction in MSL-N liquid. Scale bar = 10 μm. **(B)** Expression level changes of *adg2*, *clr5*, *ste11*, and *sms1* in all 12 lines of *prm1Δ* evolved strains after 5 hours of mating across pseudo-time. X axis is pseudo evolutionary time and Y axis is normalized expression level. **(C)** Venn diagram showing the intersection of differentially expressed gene sets after 5 hours of mating in *prm1Δ* ancestor strains with *clr5Δ*, *ste11*OE, and *sms1* OE, respectively (RNA-seq analysis). The 26 genes co-differentially expressed are listed on the right.

**Supplemental movie S1. Aberrant behavior of CRIB in sms1OE strain, related to Figure 6C**

CRIB-GFP fluorescence time-lapse images captured starting approximately 4 hours after starvation in homothallic strains, either overexpressing *sms1* or not, both in the *prm1Δ* ancestor background. The green arrows mark cell pairs that have CRIB-sfGFP signal. Time is in hours: minutes. Scale bars, 5μm.

