## supplementary information for "Genomic and Epigenetic Interplay Drives Adaptive Fusion via Reproduction Trade-Off"

#### 1. Evolution Experiment: Setup and Rationale

##### 1.1. Ancestral Strain Construction for Experimental Evolution

To preclude any potential evolutionary biases attributable to nutrient dependencies in auxotrophic strains, the Prm1 deletion cassette was obtained from the corresponding deletion library via Polymerase Chain Reaction (PCR) and subsequently introduced into the prototrophic h90 strain ySM1396 (yGL705). To high-throughput screening of fusion efficiency, a variety of fusion-specific promoters were evaluated, including P*isp3*, P*meu14*, P*meu31*, and P*SPAC13A11.06*. Among these, the P*meu31*-GFP construct was identified as the most suitable for specific expression following successful cell fusion (Fig. S1C). The utilization of this genetic marker facilitated the screening of approximately 10,000 cells per colony, enabling the determination of relative fusion efficiency based on the percentage of GFP-positive cells (Fig. S1D). Consequently, the *prm1Δ* P*meu31*-GFP h90 strain was selected as the foundational strain for the evolutionary experiment.

##### 1.2. Quantification of Relative Fusion Efficiency via Flow Cytometry

Flow cytometry, specifically using the NovoCyte Flow Cytometer, was determined to be a viable method for distinguishing between fused and unfused cells, based on the histogram and scatter plot analyses of the FIFC-A (X-axis) and PI channel (PE-Cy5-A) (Y-axis, Fig. S1D). Cells that were successfully gated were successfully fused mating pairs. It is important to note that flow cytometry registers tetrads as single events, and unfused cells as either a single event if they remain adhered or as two separate events if they disassociate. The exact fusion efficiency was calculated using the formula: fused pairs / (unfused pairs + fused pairs) (Fig. S1A-B). Therefore, the readings from the flow cytometer do not represent absolute fusion efficiency but rather provide a clear indication of the relative changes in fusion efficiency. In contrast to the nearly 100% fusion efficiency observed in wild-type strains as measured by microscopy, the relative fusion efficiency of wild-type cells after 24 hours of mating, as measured by flow cytometry, ranged from 10% to 33%, contingent upon batch-specific conditions (Fig. 1D, S1E). To mitigate this experimental variability, the *prm1Δ* ancestral strain was included as a control in each batch of measurements throughout the experimental evolution, and its measured fusion efficiency was used as a normalization value for the evolved strains.

##### 1.3. Experimental Evolution Protocol

Initially, a tetrad dissector was employed to isolate 96 isogenic single *prm1Δ* P*meu31*-GFP h90 cells, which were then plated on YE plates and allowed to proliferate for four days (Fig. 1C). Subsequently, the cells were transferred to MSL-N plates for a 24-hour incubation period via gentle replica plating, as excessive cell loading was found to compromise fusion efficiency, even in wild-type cells. The resulting 96 sporulation products were then analyzed by flow cytometry to determine the GFP-positive population. A consistent gating methodology was applied across all experiments to measure the percentage of the GFP-positive population.

Recognizing the potential for multiple evolutionary pathways for the *prm1Δ* strain to achieve higher fusion efficiency, the 12 colonies exhibiting the highest fusion efficiencies were selected as the starting points for parallel evolution experiments. To maintain a high selection pressure, the tetrad dissector was utilized to isolate successfully fused spores for each subsequent round of evolution, as it was observed that batch treatment with glusulase to eliminate non-spore, vegetative cells was not completely effective. Given the laborious nature of spore isolation via tetrad dissection, six spores from independent tetrads of the strain with the highest fusion efficiency in each of the 12 evolving lines were chosen to perpetuate the experiment.

Following their placement on YE medium, the cells were incubated for four days. These cells were then arrayed with their immediate parent strains in row A and all tetrad-dissected spores in rows B through G for all lines, alongside 12 *prm1Δ* ancestor controls in row H. A portion of each culture was cryopreserved in 96-well plates with 25%

glycerol at -80°C. The remaining cell cultures were replica-plated onto MSL-N medium, along with freshly inoculated cultures of their immediate parents and the *prm1Δ* ancestor, which had been pre-cultured for 24 hours after a two-day inoculation period. These 96 strains were then subjected to flow cytometry analysis in the same manner as the *prm1Δ* ancestor strains. The fold change for each strain was calculated by normalizing the GFP-positive readings from rows A-G to the value obtained from the *prm1Δ* ancestor in row H.

Following this procedure, the spore with the highest fusion efficiency among the six spores (from rows B-G) within the same line was selected, and six spores from six independent tetrads of that highest-performing sporulation product were isolated for the next round of evolution. These spores were subsequently arrayed on YE plates to initiate the subsequent evolutionary cycle. Each cycle involved a four-day incubation on YE medium, followed by a 24-hour period on MSL-N medium, with screening conducted on the sixth day and the subsequent round of selection commencing on the seventh day. Each mitosis-meiosis evolution cycle thus spanned a total of seven days. The entire experiment was conducted over a period of six months, with some cycles (2, 3, and 4) being repeated due to instances of contamination. The evolution experiment was concluded at the 18th cycle due to the COVID-19 lockdown imposed on March 13, 2020, in Switzerland.

##### 1.4. Selection of Strains for Sequencing

The 18 rounds of evolution resulted in a total of 18 × 96-well plates of cryopreserved stock. For all sequencing analyses, the spore exhibiting the highest fusion performance in each of the 12 lines from the 3$^{rd}$, 6$^{th}$, 9$^{th}$, 12$^{th}$, 15$^{th}$ and 18$^{th}$ cycles was selected, resulting in a total of 72 strains for sequencing. For each of these strains, three biological replicates were sequenced. As controls for the RNA sequencing experiments, wild-type, *prm1Δ* ancestor, and e-Strain (line 3, cycle 9) samples were included in every batch.

### 2. Transcriptomic Analysis Pipeline

#### 2.1. RNA Sequencing Sample Preparation

For each strain under investigation, three biological replicates were collected for RNA sequencing. To revive the wild-type, *prm1Δ* ancestor, and evolved strains from -80°C storage, they were streaked onto YES solid agar plates and incubated at 30°C for two days. On the first evening, freshly streaked colonies were inoculated into 50 mL culture tubes containing 10 mL of YES liquid medium and incubated overnight with shaking at 30°C and 200 rpm. The following morning, the cell suspensions were diluted in YES liquid medium to an optical density at 600 nm ($OD_{600}$) of 0.025, to ensure that the $OD_{600}$ of the culture on the subsequent morning would be within the range of 0.4 to 0.8. On the second evening, the cells were diluted to an $OD_{600}$ of 0.2 in 20 mL of YES liquid medium. For each sample, six tubes were prepared, with three designated for the 0-hour treatment and three for the 5-hour nitrogen deprivation treatment. The strains were then incubated overnight at 30°C with shaking at 200 rpm.

On the third morning, the $OD_{600}$ was measured to confirm that the cultures had reached a cell density of approximately 0.8. The cells were then pelleted by centrifugation at 1,000 × g and transferred to 1.5 mL tubes. For the 0-hour samples, the pellets were washed three times with PBS buffer following centrifugation. The supernatant was carefully and completely removed, and the samples were subsequently stored at -80°C. For the 5-hour nitrogen starvation treatment, the cells were washed three times with 1 mL of MSL-N medium, resuspended in 3 mL of MSL-N medium, and the cell density was adjusted to an $OD_{600}$ of 1.5. A multichannel pipette was used to transfer 96 aliquots of 20 μL cell suspension at an OD of 1.5 onto MSL-N solid square agar plates, which were then incubated at 25°C for 5 hours.

Following the 5-hour incubation under nitrogen-depleted conditions, a sterile plastic spreader was used to thoroughly scrape the cells from the plate surface to maximize recovery. The collected cells were transferred to a 1.5 mL microcentrifuge tube and washed twice with PBS buffer to remove any residual medium or debris. The supernatant was carefully removed with a micropipette to minimize any remaining liquid, and the cell pellet was rapidly frozen in liquid nitrogen, stored at -80°C, and subsequently sent to Novogene Corporation for RNA sequencing. A dry cell mass of 20-30 μL was collected from each sequencing sample for RNA extraction. RNA extraction, quality control, and the preparation of RNA sequencing libraries were performed by Novogene, and sequencing was conducted on the Illumina NovaSeq 6000 platform, yielding 2G of 150 bp paired end reads per sample. The quantity of the extracted RNA was assessed using the Bioanalyzer 2100 (Agilent Technologies).

##### 2.2. RNA-Sequencing Quality Control

The raw RNA-seq data were processed within a Linux environment. An initial quality assessment was carried out using FastQC (v1.4.0). Low-quality reads and adapter sequences were subsequently removed using Trim Galore (v0.6.10). The cleaned reads were then aligned to the reference CDS genome (2024.08.20 release, PomBase). Transcript quantification was performed using Salmon (v1.4.0).

##### 2.3. Batch Effect Correction

Transcripts-per-million (TPM) values were calculated based on the gene-specific effective lengths as reported by Salmon (v1.4.0)[1]. To correct technical variability arising from six independent sequencing runs, a unified TPM expression matrix was constructed and subjected to batch correction using the ComBat method from the sva R package (v3.48.0)[2]. The batch variable was defined by the sequencing run identifiers (six batches), while the group variable represented the biological conditions, including Ancestor, e-Strain, and Wild Type. Prior to the correction, TPM values were log-transformed using the formula $log_2(TPM + 1)$ to stabilize the variance across genes. The batch correction was then applied to the entire dataset in a single, unified step. The resulting adjusted expression matrix was retained for all subsequent downstream analyses. Gene-specific effective lengths were maintained throughout the analysis pipeline to allow for the recalculation of TPM values if required.

##### 2.4. Quality Control of Replicates

Principal Component Analysis (PCA) was conducted using the FactoMineR R package (v2.8) on $log_2(TPM + 1)$ transformed data[3]. The analysis was performed in three distinct modes: 1) a full dataset analysis including all time points (0h and 5h nitrogen starvation), and 2) time-point-specific analyses conducted separately for the 0h and 5h conditions. The biological replicates demonstrated close clustering in the resulting PCA plots, indicating a high degree of reproducibility.

##### 2.5. Gene Differential Expression Analysis

Differential expression analysis was carried out in R (v4.3.3) utilizing the limma package (v3.56.2) on the batch-corrected, log-transformed expression values ($log_2[TPM + 1]$)[4]. The analyses were conducted independently for the two experimental time points (0 h and 5 h under nitrogen starvation). For each time point, a linear model was constructed using a design matrix in which each evolved strain and the ancestral control (yGL0447) were encoded as independent factors, without an intercept term. Pairwise comparisons between each of the 72 evolved strains (in triplicate) and the ancestral reference at each time point were performed using moderated t-statistics, followed by empirical Bayes shrinkage. Differentially expressed genes were identified based on Benjamini-Hochberg adjusted p-values below 0.05, and two distinct fold-change thresholds were applied: an exploratory cutoff of $|log_2$ fold change$| \geq 0.5$ and a more stringent threshold of $\geq 1$. The resulting contrast-specific gene lists were then exported for downstream interpretation.

#### 3. Evolutionary Transcriptome Trajectory Profiling

##### 3.1. Gene Expression Dynamics Heatmap

The temporal expression dynamics of nitrogen starvation-responsive genes across the evolved lineages were analyzed using an integrated pipeline that combined fuzzy c-means clustering from the clusterGVis R package (v0.1.2) with functional enrichment analysis. The adjusted $log_2(TPM+1)$ values were clustered into 12 co-expression modules using the clusterData function, with the Euclidean distance metric and the default fuzzification parameter (m=1.25)[5]. The cluster-specific gene sets were then subjected to Gene Ontology (GO) enrichment analysis using the clusterProfiler R package (v4.8.1)[6], querying the GO database (AmiGO 2, 2025.03 release) with stringent thresholds (p-values ≤ 0.01). Biological Process terms were prioritized, and the top enriched pathways for each cluster were visualized using radial heatmaps and concept networks generated with ClusterGVis (v1.1.3)[5]. The visualization pipeline incorporated a lineage-stratified color-coding scheme (12 distinct hues from #00468B to #EFB6C8), with directional expression gradients derived from the row-wise Z-score normalization of the gene expression values (Fig. S2-3).

##### 3.2. Highly Variable Gene Set Identification

To investigate transcriptomic variability across the experimental lineages, a comprehensive computational pipeline was employed, integrating a multi-scale analysis of gene expression dynamics. The methodology began with baseline normalization, which involved subtracting the control sample (Ancestor strain) expression values through vectorized matrix operations, while excluding genes associated with the chromosome I loss in line 3 to eliminate artifacts from structural variation. Transcript consistency was quantified using a modified coefficient of variation (CV) metric with ε-shifting (1e-6) to prevent division errors in low-expression genes: $CV = \sigma/(\mu+\varepsilon)$, where $\sigma$ represents standard deviation and $\mu$ denotes mean expression across experimental lineages.

Genes were dynamically stratified into variability quartiles through adaptive thresholding (90th, 80th, and 50th percentiles, with the remainder in the last 50%, Fig. S4A) using quantile regression, which enabled the identification of hypervariable transcriptional signatures. Density plots were then generated to analyze the CV distribution in both the line3 and expline3 sets (Fig. S4B). A comparative analysis was performed by intersecting the high-variability gene sets (top 20% CV thresholds) between the lineage-specific datasets (line3 vs. expline3), which identified 271 unique genes in line3, 271 unique genes in expline3, and 748 overlapping genes that were subsequently validated through phased visualization (Fig. S4C). The total number of highly variable genes (HVGs) used for the subsequent analysis was 1290.

The adjusted $log_2(TPM + 1)$ values of the intersecting high-variability gene sets were then clustered using ClusterGVis with fuzzy C-means clustering to identify biologically relevant patterns within the data. Gene Ontology (GO) enrichment analysis was performed on the resulting clusters. Biological processes with adjusted p-values below 0.01 were considered significantly enriched and were annotated on the heatmap to aid in the interpretation of the functional relevance of the clustered gene sets. To further enhance interpretability, the GO terms were grouped into functional categories: pheromone-related (orange), metabolic (green), and cell cycle-related (blue), and were displayed on the heatmap (Fig. 2A).

##### 3.3. Pseudotime-Based Classification of Temporal Expression Patterns

All identified high-variability genes were initially subjected to trajectory profiling using ClusterGVis, which resulted in the classification of the genes into four distinct trajectory classes. Two of these classes, comprising a total of 928 genes, exhibited clear oscillatory behavior and were consequently merged to form a single gene set for subsequent pseudotime analyses. The pseudotemporal ordering of gene expression dynamics across the evolutionary strains was performed using the bulkPseudotime R package (v0.1.0), in conjunction with ClusterGVis (v0.1.2) for visualization[5].

For each evolved lineage, post-correction $log_2(TPM + 1)$ values were Z-score normalized and re-ordered in pseudotime space, with the Ancestor strain yGL0447 serving as the root, using the default settings of bulkPseudotime. A pseudotime heatmap was generated using the pseudotime_heatmap() function from the bulkPseudotime package. The order of the genes along the y-axis was maintained consistently for all lines, following the gene order established after clustering into three major groups in line 3. The heatmap annotations above the plot indicate the evolutionary cycle to which each strain belongs, with each cycle being annotated with a specific color code: "#1B1919FF", "#00468BFF", "#42B540FF", "#0099B4FF", "#FDAF91FF",

"#AD002AFF", "#ADB6B6FF". The Z-scores of the gene expression values in the heatmap were color-coded, ranging from blue (low expression) to red (high expression). Gene-wise expression trajectories were visualized using ggplot2 (v3.4.2)[7], with individual genes represented as semi-transparent blue lines and the overall pattern summarized by a red median trend line. An inspection of the resulting profiles allowed for the manual assignment of the 12 evolved lineages into four major expression-trajectory categories. The computational reproducibility of the pseudotime values was ensured by setting a fixed random seed (set.seed = 123, Fig. 2B).

To further investigate the temporal expression patterns of key rescue genes involved in differentiation and signaling, the pseudotime-aligned expression trajectories of four biologically relevant genes—*sms1*, *ste11*, *clr5*, and *adg2*—were extracted. The Z-score normalized $log_2(TPM + 1)$ values were obtained from the full pseudotime matrix of each evolved lineage (excluding WT), which had been previously aligned using the bulkPseudotime framework. The lineage-specific expression dynamics were plotted using ggplot2, with pseudotime on the x-axis and normalized expression on the y-axis[7]. The expression of each gene was faceted independently and grouped by evolutionary lineage to facilitate a direct comparison across different genetic backgrounds and time points (Fig. S11B).

##### 3.4. Correlation Analysis of Fusion Efficiency and Growth Rate

To determine whether the observed wave-like gene expression pattern was a result of a shift between mitosis and meiosis, the growth rate (measured by a spectrometer, Table S1J) and fusion efficiency (measured by FACS, Table S1I) of all 72 strains were quantified in triplicate. The experimental details for these measurements are described in supplementary information sections 5.5 and 5.6. Linear regression analyses were performed using ggplot2 (v3.5.1) and ggpmisc (v0.6.1) to model the relationships between the average cellular growth rates and the average relative fusion efficiency across the evolutionary cycles (03–18)[7]. Five pairwise comparisons were evaluated: (1) 03 vs. 06, (2) 03 vs. 09, (3) 03 vs. 12, (4) 03 vs. 15, and (5) 03 vs. 18. Additional analyses of adjacent cycle transitions (06–09, 09–12, 12–15, 15–18) were also conducted (Table S1J). For each comparison, the datasets were filtered and subset using dplyr (v1.1.4), and linear models were fitted using geom_smooth(method = "lm") with 95% confidence intervals. The equation coefficients (slope ± SE) and adjusted $R^2$ values were directly annotated on the plots via stat_poly_eq, implementing the formula $y = \beta_0 + \beta_1 x$ (Fig. 3A), and the corresponding R values were reported.

##### 3.5. Integration of Transcriptomic and Phenotypic Patterns

Batch-corrected TPM values, excluding those from the wild-type strains, were subjected to non-linear dimensionality reduction using t-Distributed Stochastic Neighbor Embedding (t-SNE, perplexity=30), as implemented in the Seurat R package (v5.2.1)[8]. The cells were clustered using a shared nearest neighbor graph (resolution=2) based on the first 10 principal components, which collectively explained over 85% of the variance, as determined by an ElbowPlot. Differential expression between the identified clusters was assessed using DESeq2 (v1.40.2), with a Benjamini-Hochberg False Discovery Rate (FDR) correction ($\alpha$=0.05)[9].

To establish a connection between the transcriptional states and the experimental phenotypes, the t-SNE coordinates were integrated with lineage-specific fusion efficiency measurements (quantified via FACS) and growth rates ($OD_{600}$ curves) using multivariate regression. Through unsupervised clustering, the transcriptional data were grouped into 8 distinct clusters, which were annotated with colored, transparent oval shapes (details can be found in Table S1E). Fusion efficiency was visualized using a color-mapped gradient scale from white to red, with specific breaks set at 0.04, 0.09, and 0.15. The growth rate was similarly visualized with a color gradient from white to dark blue, ranging from 0.26 to 0.52. Each point on the plot represented a cell, with its size corresponding to the cycle time point (0, 3, 6, 9, 12, 15, 18 cycles). In addition, the trajectory of the cells across the different time points was tracked and annotated with a dashed arrow line. This trajectory line was overlaid on the t-SNE plot to highlight the movement of the cells during the course of evolution, providing a visual representation of the transitions between the cycle time points (Fig 3B, Table S1F).

##### 3.6. Mitotic and Meiotic Correlation Analysis: *Linking Transcriptional Patterns to Functional Phenotypes*

The transcriptomic dynamics associated with cellular fusion efficiency and growth rate were systematically investigated using a computational pipeline based on Least Squares Estimation (LSE). The analysis utilized post-correction $log_2(TPM + 1)$ values, with long non-coding RNAs excluded to focus on protein-coding genes. For each experimental lineage, the gene expression trajectories were scaled through minimum-maximum normalization relative to the phenotypic parameters (fusion efficiency and growth rate) to enable a cross-comparison of expression patterns. The LSE was calculated between the scaled expression profiles and the corresponding phenotypic measurements using row-wise matrix operations. Genes were subsequently stratified into variability quantiles through adaptive thresholding, with genes falling into the top 10th and bottom 10th percentiles being selected for further analysis. Genes with LSE $\leq$ 10% were considered to be phenotype-synchronized candidates, as they displayed minimal deviation from the phenotypic trajectory. Conversely, genes with LSE $\geq$ 90% were considered to have an inverse relationship with the phenotype.

To visualize the expression trends of the top 10 percent and bottom 10 percent of genes for each phenotype (fusion efficiency and growth rate) for both phenotype-synchronized candidates and genes with an inverse relationship, ggplot2 was employed, with the x-axis ordered according to the chronological progression of the lineage cycles. The data were presented in a facet grid to distinguish the expression patterns across the different lineages. The gene expression trajectories were represented as lines, with each gene's expression across the selected lineages grouped accordingly, allowing for a detailed comparison of their temporal dynamics. The corresponding phenotypic data were overlaid as dashed lines to facilitate a direct comparison with the gene expression patterns (Fig. 3C-D, S4E-F).

Temporal clusters were identified through fuzzy C-means clustering (centers=4), which was applied to the intersecting gene sets that exhibited inverse correlations between fusion efficiency (at 0h and 5h) and growth rate (at 0h and 5h). Two specific conditions were examined: 1) 5h FE-GR anti-correlation, where genes showed a positive correlation with fusion efficiency (≤ 10% LSE) but a negative correlation with growth rate (≥ 90% LSE) at 5h; and 2) 0h inverse FE-GR coupling, where genes displayed a negative correlation with FE (≥ 90% LSE) but a positive correlation with GR (≤ 10% LSE) at 0h (Fig 3C-D, S4E-F, Table S1G). These intersecting gene sets reflect temporal trade-offs in transcriptional regulation and were subjected to functional enrichment analysis using the enricher function of the clusterProfiler package, which implemented custom Gene Ontology annotations to account for organism-specific characteristics. Custom Gene Ontology annotations, tailored to the specific characteristics of the organism, were incorporated into the enrichment analysis. Biological processes (BP) within the GO framework were considered significant if the corresponding p-values were ≤ 0.1. GO terms related to specific biological functions were visualized and color-coded as follows: orange for pheromone-related processes, green for metabolic processes, red for meiotic processes, and blue for mitotic processes (Fig. 3E).

### 4. Candidate Gene Mapping through Transcriptional and Genomic Information Integration

##### 4.1. Transcriptional Modularity Analysis and Candidate Gene Identification

***Candidate Gene Identification Through Comparative Transcriptomic profiling***

To investigate the differential gene expression patterns in the e-Strain under nitrogen starvation conditions (0h vs. 5h) in comparison to the ancestral controls (Ancestor) and the wild-type strain, the processed RNA-seq data from the e-Strain were analyzed using customized scripts that integrated differentially expressed genes from two distinct modules. The first module included the following comparisons: 0 h Ancestor vs WT, 0 h Ancestor vs e-Strain, 5 h Ancestor vs WT, and 5 h Ancestor vs e-Strain. The second module comprised the comparisons of 5 h Ancestor vs *clr5Δ*-in-Ancestor and 5 h e-Strain vs *clr5Δ*-in-e-Strain. For visualization purposes, the top 10 most upregulated and

downregulated genes from each contrast were selected and supplemented with biologically relevant candidates (*sms1*, *ste11*, *clr5*) to construct a highlight gene set.
A multi-contrast differential expression plot was generated using ggplot2 (v3.5.1)[7], displaying the $log_2$ fold-change on the y-axis and the contrast groupings on the x-axis. The genes were plotted using jittered horizontal positions (width = 0.4 via geom_jitter(); reproducibility ensured with set. Seed (123)), with the point size scaled to the magnitude of the $log_2$ fold-change. Significance categories were color-coded (red for upregulation, blue for downregulation, and grey for non-significant genes), and intelligent label placement with collision avoidance was implemented using ggrepel (v0.9.6, box. Padding=0.4 units). The contrast groupings were annotated using colored tile strips, and reference lines were included at ±0.5 $log_2$ fold-change to indicate the significance thresholds (Fig. S7).

***OptICA Analysis***

Independent Component Analysis (ICA) was performed on the batch-corrected $log_2(TPM+1)$ expression matrices using PyModulon (v0.2.1) to identify condition-specific transcriptional regulons, known as iModulons[10]. The analysis pipeline integrated two key biological annotation resources: 1) a manually curated Transcriptional Regulatory Network (TRN) containing 3,142 regulator-target pairs from BIOGRID (v4.4.242), which enabled hypergeometric enrichment testing for transcription factor-associated iModulons with Benjamini-Hochberg FDR correction; and 2) Gene Ontology (GO) term annotations from PomBase (2024.08 release), filtered for high-confidence associations. Regulatory iModulons were identified through a combinatorial enrichment analysis (maximum of 2 regulators per module) of TRN interactions, while their functional characterization was based on hypergeometric tests against GO terms ($q<0.05$). Single-gene iModulons were detected via z-score thresholding ($|z|>5$) of the gene weights relative to the module distributions. The final annotations categorized the modules as regulatory (TRN-enriched) and were visualized using the plot_activities() and plot_gene_weights() functions.

The analysis successfully identified three iModulons that were related to evolved loss genes, *ecl1*-TORC1related genes, and *ste11*-related genes, as shown in the top panel of the corresponding figure. Each plot in this panel displays the gene start positions on the x-axis and the gene weights on the y-axis, where the gene weights indicate the strength and direction of regulation. Labeled genes were highlighted due to their prominent roles in the regulation of these modules, serving as key drivers of the observed transcriptional changes. The middle panel illustrates the overall activity of the identified iModulons across the different time points, with the x-axis representing the lineage cycles (e.g., Cyc.03-0h, Cyc.03-5h) and the y-axis depicting the iModulon activity. This activity reflects how strongly each iModulon is expressed during the various experimental cycles, with color-coded data points highlighting the temporal fluctuations in activity in line 3. This panel emphasizes the temporal regulation of these iModulons, showing peaks or troughs in activity that indicate specific cycles where the gene expression associated with each modulon is notably high or low. The bottom panel focuses on the activity in Line 3, specifically illustrating the differential activity of the modulon in this lineage across the experimental cycles. The bar plots display the mean activity of Line 3 during the various cycles, with color-coded bars representing the intensity of this activity. These results reflect the degree of modulation of gene expression related to the iModulons in Line 3 across the different cycles. A list of genes was identified as candidates for the rescue of fusion efficiency based on this analysis (Fig. S8A).

***cMonkey Analysis***

A co-expression network analysis was performed using cMonkey2 with the post-correction $log_2(TPM+1)$ values[11]. cMonkey2 is a computational framework for identifying condition-specific transcriptional modules through the iterative integration of multi-omics data. The analysis pipeline was optimized by disabling operon prediction (--nooperons) and de novo motif discovery (--nomotif), focusing instead on precomputed regulatory features from the provided reference feature database from PomBase. The algorithm executed 250 iterations (--num_iterations) to ensure the convergence of the probabilistic models, iteratively refining the gene clusters based on expression correlation, evolutionary time points, and functional annotation enrichment. The output modules were annotated with transcription factors binding sites and pathway associations, using the results from the cMonkey2 bi-clustering where line 3 was present but the ancestor was not (Fig. S8B upper panel). Protein-protein interaction data from the STRING database (--string) were applied to visualize the connectivity between the modular genes (Fig. S8B lower panel). A list of genes was identified as candidates for the rescue of fusion efficiency from this analysis.

***UpSetR Analysis***

To identify genes that were consistently upregulated across the late evolutionary stages, beginning with the e-strain, the analysis focused on the differentially expressed genes (DEGs) from evolved line 3 at cycles 9, 12, 15, and 18. Genes were classified as upregulated if they met a strict threshold of $log_2$ fold change $\geq 1$ and an adjusted p-value $\leq 0.05$ (Benjamini–Hochberg corrected). To assess the overlap of the upregulated gene sets across these cycles, the intersections of line 3 at cycles 9, 12, 15, and 18 were visualized using the UpSetR package (v1.4.0, Fig. S8C)[12]. The visualization highlighted the number of genes that were shared across the different combinations of cycles, with the global intersection emphasizing the genes that were upregulated in all post-Cyc09 samples. This intersection represented the core set of persistently upregulated genes during the later stages of evolution in line 3.

Enrichment analysis was performed using custom Gene Ontology (GO) annotations that spanned the Biological Process (BP), Cellular Component (CC), and Molecular Function (MF) ontologies. Significantly enriched terms (adjusted p $\leq$ 0.05) were visualized, with functional categories of interest color-coded in red for emphasis. Genes that were found to be consistently upregulated from cycle 9 through cycle 18 and were enriched in mating-related GO terms were selected as candidate genes. The lists of genes obtained from the OptICA, cMonkey, and UpSetR analyses were combined, and an arbitrarily selected list of 98 genes was formed for subsequent experimental validation[12].

To identify co-regulated gene modules within the full dataset, two complementary machine learning approaches were integrated: Independent Component Analysis (OptICA), which revealed a key module driven by *ste11* (Fig. S8A), and a bi-clustering algorithm (cMonkey), which pinpointed a gene cluster that was highly correlated with the phenotype of line 3. The protein interaction analysis from this approach showed *ste11* as an interaction hub and *sms1* as a promising candidate (Fig. S8B). The analysis also specifically filtered genes whose expressions commonly changed in line 3 after cycle 9 (Fig. S8C-D), thereby focusing the search on the drivers of the fixed phenotype.

**4.2. Shotgun Whole-Genome Sequencing and Data Processing**

The spore with the best fusion performance from each of the 12 lines in the 3rd, 6th, 9th, 12th, 15th, and 18th cycles, totaling 72 strains, were sequenced along with the wild-type (yGL705) and ancestor strains (yGL447), which were the same strains used for the RNA sequencing. For each strain, 10 ml of a saturated culture was obtained, and DNA was extracted following the standard phenol-chloroform DNA extraction protocol. The sequencing library was prepared by the University of Lausanne genomic center, and sequencing was performed using an Illumina Hi-seq instrument.

The whole-genome sequencing data were processed through Varathon (v1.0.0; GitHub: yjx1217/Varathon) using the default parameters, which are optimized for the detection of multiple variant types, including single nucleotide variants (SNVs), small insertions/deletions (INDELs), structural variants (SVs), and copy number variations (CNVs). High-confidence variants were retained by applying a quality threshold (QUAL > 30). The functional annotation of the variants was performed using SnpEff (v5.1d), with the *Schizosaccharomyces pombe* reference genome (from the PomBase release in 2024.08) configured as a custom database. The SnpEff (v5.1d) database was built by integrating the reference genome (FASTA) and structural annotation (GFF3) files into the snpEff.config configuration, followed by the generation of an index[13].

***Strain Sequence Coverage Analysis***

Sequencing coverage depth files (.txt) were processed using a custom R pipeline. The coverage data were extracted for chromosomal region I (positions ≥5,428,356) across all samples. The samples were labeled according to their experimental IDs (e.g., Ancestor, Cycle03line03) and were combined into a unified data frame. The coverage profiles were visualized using ggplot2 (v3.4.2) as multi-panel line plots (geom_line, linewidth=0.6), with a facet grid (facet grid) used to vertically align the individual sample trajectories (Fig. 4B)[7].

***Structural Variation Analysis***

Variant Call Format (VCF) files containing the genomic variant information were processed by the circlize package (v0.4.15). The chromosomal architecture was reconstructed based on the *Schizosaccharomyces pombe* reference genome coordinates (Pombase, 2024.08 release) [13], with the centromeres annotated as trapezoidal regions using experimentally defined positions (Chr I: 3,753,687–3,789,421; Chr II: 1,602,264–1,644,747; Chr III: 1,070,904–1,137,003). Single nucleotide polymorphisms (SNPs) and insertions/deletions (INDELs) were binned into 10-kb intervals, and the variant frequencies were calculated as the proportion of mutant alleles across the 80 sequenced clones. Copy number variations (CNVs) were mapped from a read-depth analysis (manta v1.5.0) and were classified as either losses (deletions) or gains (amplifications). All these mutations were manually curated by excluding any mutations that were already present in the ancestor strains (yGL447).

Circos plots were generated to integrate four distinct data layers: 1) Chromosomal ideograms with centromeric constrictions visualized in maroon (#8B0000); 2) SNP density (points scaled 0.6–1.0×, with red indicating >50% allele frequency and blue indicating ≤50%); 3) INDEL distribution (orange for high-frequency and purple for low-frequency); and 4) CNV segments (black for loss and magenta for gain). All coordinates were normalized to chromosomal lengths (Chr I: 5.58 Mb; Chr II: 4.54 Mb; Chr III: 2.45 Mb) using non-overlapping circular tracks (Fig. 4A outer ring, table S1H).

The variant burden across single nucleotide polymorphisms (SNPs), insertions/deletions (indels), structural variants (SVs), and copy number variants

(CNVs) was quantified through a multi-step visualization pipeline. Comparative burden analyses were performed using violin plots (ggplot2::geom_violin), with cycle-stratified distributions showing the median/quartile lines (draw_quantiles = c(0.25, 0.5, 0.75)) and alpha-blended kernels ($\alpha$=0.8) (Fig. S5A).

Genetic variant classification was carried out using custom R scripts that processed the SnpEff annotation outputs. For each sample, the variant effects were quantified by summing the counts across the different functional categories (e.g., missense, synonymous, splice-site) and biotypes (protein-coding, lncRNA) from the snpEff-generated genes.txt files. The sample-level counts were then aggregated into a matrix through iterative merging, with any missing values imputed as zero. A comprehensive overview of the genetic variants present across the experimental samples was visualized using a stacked bar chart created with ggplot2. The x-axis represented the samples, and the y-axis displayed the variant counts. Each bar in the chart was stacked according to the variant type, with different colors representing the different variant classes (e.g., missense, frameshift, 3_prime_UTR) in comparison to the Ancestor strain (yGL0447). By visualizing the counts of the different types of variants and their distribution within each sample, insights were gained into the genomic variations that differentiate the evolved strains from the Ancestor (Fig. S5B).

To analyze the distribution of variant types across the samples, the percentage of each variant type relative to the total number of variants in each sample was calculated. This was achieved by summing the total variant counts per sample and then calculating the proportion of each variant type. The dataset was grouped by line name, and the percentage of each variant type was computed for each sample. The results were then aggregated to compute the average percentage of each variant type across all samples. The final visualization was generated using a pie chart to represent the distribution of variant types across all experimental strains. The mean percentage of each variant type was plotted, with less common variants (those with a mean percentage below 1%) being grouped into an "Others" category for clarity. The chart was generated using the ggplot2 package with the geom_bar() function, followed by the coord_polar() function to convert the bar chart into a pie chart. The labels and proportions for the two most common variants were annotated within the pie chart (Fig. 4A, Table S1H)[7].

The analysis was further extended by generating a heatmap to visualize the impact of the variants in a more intuitive manner. The ComplexHeatmap package was used to create the heatmap, with each row representing a specific gene and each column representing a particular experimental condition or line name[15]. The heatmap was color-coded based on the impact count, with higher counts being represented by a gradient of colors ranging from white to red, where red indicates a higher impact. Additionally, the circlize package was used to customize the heatmap annotations, including the addition of bar plots to show the total variant count for each sample and text annotations to display the gene names and variant impacts[14]. To focus on the most relevant data, the SNPEff annotations corresponding to the genes of interest were filtered based on a check of the mapping bam file through IGV (v2.15.4)[13]. This allowed for a more targeted analysis of the specific genes that were of biological interest. The cleaned dataset was then used to generate the final impact analysis (Fig. S5C).

##### 4.3. Long-Read Whole-Genome Sequencing

The samples analyzed in this study were prepared using a standard protocol for yeast strains, with each sample consisting of three biological replicates. To begin, the yeast strains were revived from -80°C storage by inoculating them onto yeast extract (YE) agar plates, followed by a one-day incubation to ensure active growth. Subsequently, 15 mL of YE liquid medium was inoculated with a single colony and grown overnight at 30°C until the optical density ($OD_{600}$) reached a range of 0.5-0.8, indicating that the cells were in the logarithmic phase of growth. The cultures were then harvested by centrifugation at 1,000 × g for 3 minutes. For each sample, three separate tubes were prepared and collected from each replicate.

The cell pellet was resuspended in phosphate-buffered saline (PBS) and transferred to 1.5 mL centrifuge tubes. The cells were then washed three times with PBS to remove any residual growth medium. Any excess liquid was carefully removed by aspiration to minimize the volume of the sample, and the tubes were tightly sealed. The combined samples from the three replicates were then rapidly frozen in liquid nitrogen to preserve the DNA and prevent its degradation. Following DNA extraction, sequencing was performed using the Oxford Nanopore Technologies (ONT) platform, utilizing the Long-Read Sequencing (LRS) method. GrandOmics employed this method to generate long reads, which enabled the sequencing of complex genomic regions and the identification of large-scale structural variants. The sequencing library was prepared following the standard protocol for the ONT platform. To identify structural variations (SVs) between the engineered (e-strain) and wild-type (WT) *Schizosaccharomyces pombe* genomes, a long-read sequencing (LRS) analysis was performed using the LRSDAY pipeline (https://github.com/yjx1217/LRSDAY), in conjunction with the NGS data. The e-strain genome was de novo assembled using canu (v2.2) with LRSDAY, which integrates long-read polishing, error correction, and scaffolding to generate chromosome-level contigs[16,17].

##### 4.4. Structural Variation Analysis Based on LRS Data

To identify and visualize large-scale genomic rearrangements between the wild-type and e-strain assemblies, a whole-genome alignment between the wild-type and e-strain was performed using nucmer (v3.1) with the parameters --maxmatch -c 50 -b 100 -l 20 to maximize the sensitivity for detecting conserved regions while allowing for rearrangements (Fig. S6A). The --mismatch flag enabled the exhaustive anchoring of exact matches, and the -c 50/-l 20 parameters ensured the clustering of matches ≥50 bp with individual matches ≥20 bp, thereby prioritizing biologically relevant syntenic blocks. The resulting alignments were filtered using delta-filter (-m -i 80 -l 50) to retain high-confidence alignments with ≥80% sequence identity and a length of ≥50 bp, which reduced the number of matches arising from repetitive or low-complexity regions.

Structural variants (SVs), including inversions, translocations, and duplications, were systematically annotated using SyRI (v1.7.0)[18], which leverages synteny and local sequence divergence to classify rearrangements. The inputs for SyRI included the filtered alignment and a coordinate file, with homologous chromosomes being standardized to ensure accurate synteny mapping. Finally, plotsr (v1.1.1) was used to generate a visualization of the SVs, integrating the output from SyRI and the genome metadata to render a plot with configurable dimensions (-W 10 -H 4). Although SyRI provides robust tools for SV annotation, certain events, such as translocations, required manual curation. Specifically, translocation events were observed between regions that appeared to show significant similarity, particularly at the chromosome ends. The high sequence homology in these regions, which is often found at the telomeric regions, can complicate SV detection algorithms by incorrectly labeling these regions as translocations. These flagged regions were manually reviewed and cross-referenced with existing genomic annotations from PomBase to confirm that what appeared to be a translocation event was, in fact, an artifact due to the inherent sequence similarity of the telomeric regions (Fig. S6A).

##### 4.5. Sequencing-Based Confirmation of Deletion Breakpoints on Chromosome I

Sanger sequencing was employed to confirm the deletion junctions that had been identified through PCR amplification. PCR products were generated using the specific primers OGL5501 and OGL5502, which were designed to amplify the regions flanking the deletion junctions. The lengths of the amplicons were subsequently verified via gel electrophoresis, and the corresponding bands were excised from the gel and purified for Sanger sequencing. The observed band was mapped to the chromosomal telomere region, based on sequence homology and alignment with known sequences at the chromosomal ends from the LRS data and the reference genome from PomBase (2024.08.20 release). The final confirmation of the 1181 bp deletion junctions was achieved through sequence comparison and alignment, which allowed for the precise characterization of the genomic rearrangements (Fig. S6A).

##### 4.6. RT-qPCR Analysis of Gene Relative Expression

Prompted by the findings from the NGS and LRS analyses, which revealed a 149 kb deletion in the e-strain located near the *clr5* locus, and considering the potential involvement of subtelomeric silencing, the expression levels of *clr5* in the evolved strains were examined. Following a procedure similar to the previously described mating assay, yeast cells for each sample were cultured in MSL-N medium for 5 hours. After the incubation period, $1 \times 10^7$ cells were collected from each condition, and total RNA was extracted using the Yeast Total RNA Extraction Kit (Sangon Biotech, B518657).

***RNA Extraction Procedure***

The cells were initially treated with 600 µL of reaction buffer and 50 µL of 12 mg/mL snailase (Sangon Biotech, B518657) for cell wall digestion, followed by an incubation at 37°C in a water bath for 5 minutes. The samples were then centrifuged at 10,000 rpm for 2 minutes at 4°C, and the supernatant was discarded. Immediately thereafter, 400 µL of Buffer Lysis-YS was added, mixed thoroughly by vertexing, and incubated at 65°C for 5 minutes. The tubes were then transferred to ice for 5 minutes before the addition of 200 µL of Buffer YK. After gentle mixing, the samples were centrifuged at 12,000 rpm for 5 minutes at 4°C, and the resulting supernatant was collected.

To bind the RNA, an equal volume of 100% ethanol was added to the supernatant and mixed well. The entire mixture was then transferred to an adsorption column placed in a collection tube and allowed to stand for 1 minute. The column was centrifuged at 12,000 rpm for 1 minute at room temperature, and the flow-through was discarded. The column was then washed twice by adding 500 µL of RPE Solution, incubating for 1 minute, and centrifuge at 10,000 rpm for 1 minute at room temperature. After the second wash, the column was centrifuged again at 12,000 rpm for 2 minutes to remove any residual liquid. The adsorption column was then transferred to a new RNase-free 1.5 mL centrifuge tube, and 35 µL of DEPC-treated dd$H_2O$ was added directly to the center of the membrane. After standing for 2 minutes, the tube was centrifuged at 12,000 rpm for 2 minutes to elute the RNA. The purified RNA was either used immediately or stored at –80°C for future use.

***RNA Quality Assessment and cDNA Synthesis***

The concentration and purity of the RNA were assessed using a NanoDrop One (Thermo Scientific, Cat. No. 840-317400), and the integrity of the RNA was verified by agarose gel electrophoresis. First-strand cDNA synthesis was then performed using the All-In-One 5× RT Master Mix with gDNA Removal Kit (Cat. No. G592; Applied Biological Materials, Richmond, BC, Canada). The volume of RNA required to obtain a total amount of 2 µg was calculated based on the NanoDrop One reading of the RNA concentration. The RNA was then diluted to 16 µL with sterile $ddH_2O$, and 4 µL of the All-In-One 5× RT Master Mix was added. All solutions were thawed, mixed thoroughly, and assembled on ice. The reaction was gently mixed, briefly centrifuged, and incubated at 37°C for 30 minutes to remove any genomic DNA, followed by a 10-minute incubation at 60°C for reverse transcription. The reaction was terminated by heating at 95°C for 3 minutes, and the resulting cDNA was placed on ice. The first-strand cDNA was then used immediately (Table S1K).

***Quantitative Real-Time PCR***

Relative quantification was performed using the PerfectStart® Green qPCR SuperMix (TransGen Biotech, AQ602). The PCR reaction system and conditions were as follows: each 10 µL qPCR reaction contained 1 µL of template DNA, 0.2 µL of 10 µM forward primer, 0.2 µL of 10 µM reverse primer, 5 µL of 2× PerfectStart® Green qPCR SuperMix, and 3.6 µL of nuclease-free water. Quantitative PCR (qPCR) was performed using a $qTOWER^3$ real-time PCR thermal cycler (Analytik Jena). The thermal cycling protocol consisted of an initial denaturation step at 94°C for 30 s, followed by 42 cycles of denaturation at 94°C for 5 s, annealing at 55°C for 30 s, and extension at 72°C for 10 s. A final data acquisition step was performed at 60°C for 15 s. To verify the specificity of the product, a melt curve analysis was subsequently performed. Fluorescence data were acquired in real-time during the extension step of each cycle. The resulting quantification cycle (Ct) values were exported for analysis. The relative expression levels of *clr5*, *ste11*, and *sms1* were calculated using the $2^{-\Delta\Delta Ct}$ method, with the expression levels normalized to the endogenous control gene, *actin1* (Table S1K).

### 5. Dissection of the Compensatory Pathway in prm1Δ Evolved Strains

#### 5.1. Strain Construction

To investigate the molecular mechanisms underlying the increased fusion efficiency observed in the *prm1Δ* evolved strains, a rescue experiment was performed on the Prm1 mutant, based on the results of the bioinformatics analysis of the evolved strains. A list of the genes investigated can be found in Fig. S9 and Table S1I. The genetic background of all *S. pombe* strains used in the present study are homothallic (h90) strains. All strains are derivatives of the standard laboratory strain ySM1396 (yGL705). The homothallic background was chosen due to its convenience in the mating and fusion experimental processes.

In the *prm1Δ* ancestor strain, *sfGFP* expression, driven by the *meu31* promoter (as described in detail in section 1.1), was introduced at the *ura4* locus with the NatMX (resistance to nourseothricin) selection marker. In the subsequent rescue experiments for the construction of the *prm1Δ* mutant strains, the overexpression of candidate genes was driven by the actin promoter and integrated at the *ade6* or *lys3* loci for those candidate genes whose higher expression correlated with better fusion efficiency. For candidate genes whose lower expression correlated with better fusion efficiency, the genes were knocked out and replaced with resistance markers. These strains were constructed using either hphMX1 (Hygromycin B) or bleMX6 (Zeocin) as selection markers. The detailed genotypes of the constructed strains can be found in Table S1Q. To account for batch-to-batch variation in fusion efficiency and to ensure that the observed changes were due to the modification of the candidate genes, controlled comparisons were implemented among the engineered strains with the backgrounds of wild-type, *prm1Δ* ancestor, and e-strains. To achieve this, all genetic modifications were attempted not only in yGL447 (h90, *prm1Δ* ancestor, ura4::pMEU31-sfGFP-NATMX), but also in yGL450 (h90, wild-type like, ura4::pMEU31-sfGFP-NATMX) and yGL636 (e-Strain, h90, *prm1Δ* evolved cycle 9 line 3, ura4::pMEU31-sfGFP-

NATMX), which served as the control strains for the rescue experiment. However, the construction of some strains failed after three trials, and these are indicated in grey in Fig S9.

#### 5.2. Media Preparation

##### *Nutrient-Rich Medium*

Yeast extract with supplements (YES) medium was prepared by mixing the following components: 5 g/L yeast extract, 30 g/L Glucose, 0.225 g/L leucine, 0.225 g/L Adenine, 0.225 g/L Uracil, 0.225 g/L Histidine, and 0.225 g/L Lysine hydrochloride. The media was then sterilized by autoclaving at 121°C for 20 minutes. For the preparation of solid media, 2% Difco Bacto Agar was added before autoclaving.

##### *Nitrogen Deficient Medium-induced meiosis* [19]

To prepare the minimum sporulation medium (MSL-N), the following components were mixed: 10 g/L Glucose, 1 g/L $KH_2PO_4$, 0.1 g/L NaCl, and 0.2 g/L $MgSO_4 \cdot 7H_2O$. This mixture was supplemented with minerals (10,000×) at a concentration of 100 µL/L, vitamins (1000×) at 1 mL/L, and 0.1 M $CaCl_2$ at 1 mL/L. The solution was then sterilized by filtration through a 0.22 µm pore size filter and stored at room temperature (RT). The 10,000× mineral stock solution was prepared with the following components: 5 g/L Boric Acid, 4 g/L $MnSO_4$, 4 g/L $ZnSO_4 \cdot 7H_2O$, 2 g/L $FeCl_2 \cdot 6H_2O$, 0.4 g/L $MoO_3$, 1 g/L KI, 0.4 g/L $CuSO_4 \cdot 5H_2O$, and 10 g/L Citric Acid. This solution was sterilized by filtration through a 0.22 µm pore size filter and stored at 4°C. The 1000× vitamin stock solution contained: 1 g/L Pantothenate, 10 g/L Nicotinic Acid, 10 g/L Inositol, and 10 mg/L biotin. This solution was also filter-sterilized using a 0.22 µm pore size filter and stored at 4°C.

#### 5.3. Live-Cell Imaging of Prm1 Localization during Meiosis

##### *Strain Construction*

To construct a Prm1 tagging plasmid, the coding sequence (CDS) and the 3′ untranslated region (3′ UTR) of *prm1* were used as homologous arms, and a fluorescent protein coding sequence along with a selectable resistance marker was inserted between them. The construction process began with an overnight digestion of the pGL42 plasmid (pGL042/pFA6a-*sfGFP*-hphMX) using *Hind*III, *EcoR*V, *Pme*I, and *BamH*I to isolate two large fragments, which served as the plasmid backbone. The CDS of *prm1* was PCR-amplified from genomic DNA using primers OGL007 and OGL008 (Table S1R), while the 3′ UTR was amplified using primers OGL009 and OGL013 (Table S1R). These fragments were then assembled into the digested plasmid backbone using In-Fusion cloning (Takara Bio USA, Inc.), resulting in a construct that tags Prm1 with *sfGFP*.

To facilitate downstream transformations, the plasmid was further digested with MluI and PmeI, and the hphMX resistance cassette was replaced by the kanMX cassette, which was PCR-amplified using primers OGL0097 and OGL0098 (Table S1R). The final plasmid was digested overnight with *Stu*I and *Mfe*I to linearize the construct, and the resulting DNA fragment was integrated between the CDS and the 3′ UTR of *prm1* using the lithium acetate transformation method. Positive transformants were screened and subsequently stored at –80°C for future use. Using the same strategy for fluorescent tagging, additional plasmids were constructed to tag *myo52* with *mTagBFP2* and D4H with *mCherry*. These plasmids were sequentially transformed into the previously generated *prm1*-sfGFP strain, ultimately yielding an h90 wild-type strain with Prm1 tagged with *sfGFP*, D4H with *mCherry*, and Myo52 with *mTagBFP2*.

***Preconditioning of Strains for Time-Lapse Microscopy***

Frozen stocks of the strains were first revived by spotting them onto YES solid medium and incubating at 30°C for two days. Freshly grown colonies were then inoculated from the solid medium into 50 mL Falcon culture tubes containing 10 mL of YES liquid medium and incubated overnight at 30°C. The cells were cultured to the mid-log phase ($OD_{600}$ = 0.5–0.8) and then harvested by centrifugation at 1000 × g for 3 minutes. After discarding the supernatant, the cell pellet was washed twice with MSL-N medium. The cell suspension was then resuspended in MSL-N medium to an $OD_{600}$ of 1.5 and incubated at 30°C with continuous shaking at 220 rpm for 5 hours[20]. Following this incubation period, the samples were ready for microscopy.

The cell concentration was determined using a spectrophotometer (Biochrom, Ultrospec 10), for which a standard curve had been established by manual cell counting with a hemocytometer. An optical density (OD) of 1.0 at 600 nm ($OD_{600}$) was determined to be equivalent to approximately 9.6 × $10^6$ cells/mL. The $OD_{600}$ measurements are used as a proxy for cell concentration in fission yeast and are linear within the approximate range of 0.1 to 0.8. Therefore, if the initial $OD_{600}$ readings fell outside this linear range, the samples were diluted or concentrated accordingly to ensure accurate and reliable measurements.

***Preparation of Microscope Slides for Overnight Time-Lapse Imaging***[19]

To prepare 2% MSL-N agarose pads for the microscopy chambers, 10 mL of MSL-N medium was mixed with 0.2 g of agarose. The mixture was then heated in a microwave on high power for approximately 2 minutes, or until the agarose was completely dissolved. Aliquots of 0.5 mL were dispensed into microcentrifuge tubes and stored at room temperature (RT). One tube of MSL-N agarose was consumed for each slide preparation.

To prepare VALAP for sealing the microscopy chambers, equal weights of lanolin, Vaseline (or another petroleum jelly), and paraffin were combined in a beaker. The mixture was heated at a low temperature, with occasional stirring, until it was fully blended. The VALAP was then aliquoted into small Petri dishes and stored at room temperature (RT). For each slide preparation, a scoopful of VALAP was used to seal the slide.

To prepare the microscopy slides, 1.5 mL Eppendorf tubes containing 2% MSL-N agarose were placed in a heat block at 95°C for 15 minutes to fully melt the agarose. Next, 100 µL of the melted agarose was pipetted onto a glass slide that had been pre-positioned with ~5 mm thick spacers at both ends. These spacers can be made from strips of cardboard. The agarose was immediately covered with a clean second glass slide to flatten the pad, and it was allowed to set for 1–2 minutes. After solidification, the top slide and spacers were carefully removed, and the agarose pad was left to air dry for about 1 minute to allow any residual surface moisture to evaporate. Then, 1 µL of a 5-hour cultured cell suspension ($OD_{600}$ = 1.5) was applied to the surface of the agarose pad. Once the cell suspension had dried completely, a coverslip was gently placed over the pad at an angle, ensuring that no air bubbles were trapped, and the edges of the slide were sealed with VALAP. The prepared slide was then left undisturbed for at least 30 minutes at 25°C or room temperature (RT) before imaging to allow the sample to stabilize. Time-lapse microscopy was subsequently performed to monitor the dynamic changes in gene expression using the fluorescently labeled proteins.

***Overnight Time-Lapse Microscopy of Fluorescent Protein-Tagged Strains***

A Nikon Eclipse TI2-E inverted fluorescence microscope with a 100× objective lens was used for most of the imaging, unless otherwise specified. Immersion oil was first applied to the objective lens, and the prepared slide was secured on the microscope stage. The image acquisition settings were then configured, ensuring that the autofocus feature was enabled. Manual focus adjustments were made as needed to achieve sharp cell resolution[19]. Unless otherwise specified, images were captured every 15 minutes over a total duration of approximately 20 hours. The images were acquired using 50-60% laser power. The exposure times were set to be between 50 and 100 ms and were adjusted within this range for each sample based on the fluorescence intensity of the tagged protein to avoid signal saturation while maintaining an adequate signal-to-noise ratio. For each sample, ten distinct fields of view were selected, each containing approximately 50 cells.

***Membrane Invagination Phenotype in prm1Δ Cells***

In *Saccharomyces cerevisiae*, the deletion of the *prm1* gene is known to cause plasma membrane invagination, a phenotype that has also been observed in *S. pombe*. To determine if this invagination consists of two layers of membrane and to study the dynamics of its formation in *Schizosaccharomyces pombe*, the mating process of $h^-$

*prm1Δ* D4H-*mCherry* crossed with h$^{+}$ *prm1Δ* D4H-*sfGFP* was imaged (Fig. 1B). The cells were imaged overnight using microscopy to observe their phenotypes, with the experimental procedure being consistent with the steps described above. The same method was applied to quantify the invagination phenotype in the rescue strains. Specifically, both successfully fused cells (zygotes) and unfused cells were scored for the presence or absence of membrane invagination during the mating process, which resulted in four distinct categories (Fig. 7D-E).

##### 5.4. Quantification of Fusion Efficiency

To quantify the fusion efficiency, three distinct methods were employed, depending on the sample throughput and the specific experimental purpose. Unless otherwise specified, the cells were pre-cultured in MSL-N liquid medium for 5 hours prior to mating.

1) **Verification of candidate gene function and evolved strains:** Following the 5-hour pre-incubation, the cells were spotted onto MSL-N agar and incubated for 24 hours. They were then scraped off and analyzed by microscopy to quantify the absolute fusion efficiency (see Section 5.4.1).
2) **High-throughput screening:** The cells were similarly mated on MSL-N agar for 24 hours, after which they were spotted onto MSL-N agar and incubated for another 24 hours. The fusion efficiency was then assessed by flow cytometry (see Section 5.4.2).
3) **Live-cell imaging of the mating process:** For the dynamic monitoring of fusion events, the cells were transferred directly to MSL-N microscopy pads after the 5-hour liquid incubation and were imaged during a 24–48-hour mating process (see Section 5.4.3).

While the absolute fusion efficiency values may vary across these different methods for the same strain, the rescue effect was consistently evaluated by comparing each strain to both the *prm1Δ* ancestor and the wild-type control. This approach ensured a reliable assessment of the ability of the evolved strain and the candidate gene to restore fusion efficiency. For all fusion efficiency analyses, the statistical significance was assessed using the Student's t-test. The data are presented as mean ± SD (*$p < 0.05$, **$p < 0.01$, ***$p < 0.001$, ****$p < 0.0001$).

##### 5.4.1. Fusion Efficiency Quantification by Microscopy of Cells Cultured on Plates

In the previous evolutionary experiment, fusion efficiency was assessed by FACS, which involved measuring the fusion rates of six spores per cycle across 12 lines over 18 evolutionary cycles. However, as FACS provides only relative changes in fusion efficiency compared to the controls, a direct quantification of the improved fusion efficiency in the evolved strains was sought using microscopy and manual analysis. Given the labor-intensive nature of this approach, the 12 strains with the highest fusion efficiency as determined by the FACS results from the 18th cycle were selected for microscopic observation to evaluate their fusion efficiency (Fig. 4F and Table S1B). To complement the next-generation sequencing (NGS) analysis, the fusion efficiency of the Line 3 strain from cycles 3, 6, 9, 12, 15, and 18 was also assessed by microscopy. The statistical results for this analysis can be found in Fig. S1 and Table S1B.

***Cell Culture***

Frozen stocks of the strains were revived on YE solid medium and incubated overnight at 30°C. On the first evening, colonies were inoculated into 10 mL of YES in 50 mL tubes and cultured overnight at 30°C and 200 rpm. The following morning, the cultures were diluted to ensure that the $OD_{600}$ would reach a range of 0.4–0.8 by the evening. The $OD_{600}$ was measured using a spectrophotometer (Biochrom, Ultrospec 10), which was calibrated for linearity between 0.1 and 0.8 ($OD_{600} = 0.1$ corresponds to $9.6 \times 10^6$ cells/mL). The cultures were diluted or concentrated as needed. On the second evening, the cells were adjusted to an $OD_{600}$ of 0.2 in 20 mL of YES and were incubated overnight at 30°C and 200 rpm. The next morning, the cultures were confirmed to have reached an $OD_{600}$ of ≈ 0.8, were pelleted by centrifugation (1,000 × g), and were then washed three times with 1 mL of MSL-N. The cells were then resuspended in 3 mL of MSL-N and adjusted to an $OD_{600}$ of 1.5. A 20 µL aliquot was spotted onto MSL-N agar and incubated at 25°C for 24 hours [20].

*Imaging*

The cells were scraped from the agar surface and resuspended in 100 µL of MSL-N in a 1.5 mL microcentrifuge tube. After centrifugation at 1,000 × g for 3 minutes, the supernatant was discarded, leaving a residual volume of 2–4 µL. The pellet was then resuspended, and 2 µL of the suspension was placed onto a microscope slide. Using a 100× objective, 15 images were captured per sample, with each field containing approximately 100 cells.

***Quantification and Statistical Analysis***

The same calculation was used to quantify microscopy-based fusion efficiency. Briefly, the numbers of fused cell pairs (tetrad formation), unfused cell pairs, and single cells were quantified using the ImageJ (v1.53, NIH) cell counter, and the percentages were calculated using the following equation:

$$\%\ \text{cell fusion} = \frac{Fused\ cell\ pairs}{Fused\ cell\ pairs + unFused\ cell\ pairs} \times 100$$

**5.4.2. Fusion Efficiency Quantification by FACS**

***Cell Culture***

In the previous evolutionary experiment, the fusion efficiency was evaluated by FACS, which involved measuring the fusion rates of six spores per cycle across 12 evolutionary lines over 18 cycles. To further validate the fusion efficiency of the evolved strains, the spore with the highest fusion performance from each of the 12 lines at cycles 3, 6, 9, 12, 15, and 18 was selected, yielding a total of 72 samples. These strains were then grouped into three batches of 25 strains each and were subjected to revival and mating assays. The fusion efficiency of the strains with modified candidate genes (93 strains, Table S1I) was also rapidly assessed by flow cytometry, following the protocol detailed in section 5.4.1. The wild-type and *prm1Δ* ancestor strains were included as FACS controls in every batch. The cells were cultured as described in section 5.4.1. Following a 24-hour incubation, the cells were harvested by scraping with a sterile pipette tip and were transferred into a 96-well plate containing 100 µL of MSL-

N medium per well. The cell suspension was then homogenized by repeated pipetting to ensure a single-cell suspension and was immediately analyzed for fluorescence on a BD FACSCelesta™ flow cytometer.

***Flow Cytometry Data Analysis***

The flow cytometry data were analyzed using FlowJo software (v10.6.2). The primary cell population was first identified and gated based on its forward scatter (FSC-A) and side scatter (SSC-A) properties. To quantify the number of GFP-positive cells, this population was further analyzed by plotting the fluorescence intensity in the FITC-A channel (for GFP) against the PE-Cy5-A channel, with both axes on a logarithmic scale. A gate was then established to distinguish the GFP-positive population from the GFP-negative population. The percentage of GFP-positive cells within the total gated cell population was calculated and defined as the relative fusion efficiency for each strain. This efficiency was then compared with that of the control strain. Representative scatter plots and statistical tables were exported for further analysis. All samples were analyzed in biological triplicates. The statistical significance between the fusion efficiency of the *prm1Δ* strain and that of the evolved strains was determined using a two-tailed Student's t-test, with p-values < 0.05 considered to be significant (Fig. 3A, Table S1I).

##### 5.4.3. Fusion Efficiency Quantification by Microscopy of Cells Cultured on Slides

The fusion efficiency of the strains carrying modifications in *clr5Δ*, *ste11OE*, *sms1OE*, *adg2OE,* and *cig2Δ* was quantified in the wild-type, *prm1Δ* ancestor, and e-strain backgrounds using slide-based microscopy. The strain culturing and mating priming were performed as described in Section 5.4.1. Following a 5-hour incubation in MSL-N liquid medium, microscopy slides were prepared according to the procedure outlined in Section 5.3. For each strain, three biological replicate cultures were spotted onto a single slide. The prepared slides were then incubated at 25°C for two days. The cells that had mated for 48 hours were imaged on a Nikon Eclipse TI2-E inverted fluorescence microscope. The P*meu31*-sfGFP signal was captured using the 488 nm channel with 50% laser power and a 50 ms exposure time. For each sample, images were acquired from approximately 15 randomly selected fields of view. The fields were chosen to contain an estimated 100 cells. The quantification was then performed as described in section 5.4.1[19].

#### 5.5. Growth Rate Measurement

To assess the overall mitotic growth rate of the evolved strains, all 72 evolved strains were revived from –80°C stocks and were cultured on YE solid medium for two days prior to the measurement. A small amount of each colony was then inoculated into a 96-deep-well plate containing 1 mL of liquid YE medium and was incubated overnight at 30°C with shaking at 220 rpm. The following day, the $OD_{600}$ values were measured by a spectrometer (Biochrom, Ultrospec 10) to ensure that the cells were in the exponential growth phase. Each sample was then normalized to an $OD_{600}$ of 0.02 in 200 μL of YE medium and was arrayed in 96-well microtiter plates for growth curve measurement using a microplate reader (BioTek, Epoch2). The experiment was conducted at 30°C for 16 hours, with the absorbance being measured at 600 nm every 5 minutes. Continuous shaking was maintained at 237 rpm throughout the assay. The growth data were analyzed using a customized R script (Fig. 3A, Table S1J). This experiment was repeated three times to ensure its reproducibility.

#### 5.6. Plasmid Construction for Candidate Genes

Unless otherwise specified, both the overexpression and knockout constructs for the candidate genes in *Schizosaccharomyces pombe* were generated using the In-Fusion HD Cloning Kit (Takara Bio, Cat. No. 639650), which facilitates seamless cloning via homologous recombination. The In-Fusion enzyme facilitates the precise joining of DNA fragments with homologous ends of 15 bp, resulting in circular recombinant plasmids. The sequences of the candidate genes were obtained from PomBase and were used to design primers for PCR amplification. Overexpression plasmids were constructed for the genes that were found to be upregulated in the evolved strains compared to the *prm1Δ* ancestor, while knockout plasmids were designed for the genes that were downregulated.

Specifically, the pGL51 plasmid (*pAde6PmeI-pAct-mCherry-D4H-hphMX1*) was digested overnight with the restriction enzymes *Kpn*I and *Not*I, and the largest fragment was purified and used as the backbone. The coding sequence (CDS) of each candidate gene was PCR-amplified from *S. pombe* genomic DNA. For the overexpression constructs, the 5′ and 3′ untranslated regions (UTRs) of the *ade6* gene served as homologous arms to mediate genomic integration at the *ade6* locus. These constructions included the actin promoter (*pAct*), the candidate gene CDS, a C-terminal *mCherry* tag, and the *hphMX* cassette, which confers hygromycin resistance. For the gene knockout constructs, the 5′ and 3′ UTRs of the target gene were used as homologous arms to replace the endogenous CDS with the *hphMX* selection marker.

##### 5.7. Construction of Strains with Overexpression or Knockout of Candidate Genes

First, the wild-type (yGL450), *prm1Δ* ancestor (yGL447), and *prm1Δ* evolved (yGL636) strains were revived from –80°C frozen stocks by streaking them onto YES agar plates and incubating them at 30°C for two days. Individual colonies were then inoculated into 10 mL of YES liquid medium and were cultured overnight at 30°C with shaking at 230 rpm until they reached the exponential growth phase. In parallel, the plasmids intended for transformation were linearized by digestion with NEB restriction endonucleases at 37°C for 3 hours or overnight.

Next, the yeast cells were harvested by centrifugation, washed twice with 1× TE/LiAc buffer, and were resuspended in 100 µL of 1× TE/LiAc. For the transformation, 5 µL of denatured salmon sperm DNA (which had been pre-heated at 95°C for 5 minutes) and 15 µL of the linearized plasmid were added to the cell suspension. The mixture was gently mixed by tapping and was incubated at room temperature for 10 minutes. Subsequently, 500 µL of PEG/TE-LiAc solution was added, mixed gently by inversion, and incubated at 30°C for at least 3 hours. Then, 43 µL of DMSO was added, mixed by inversion, and the cells were heat-shocked at 42°C for 5 minutes.

Afterward, the cells were resuspended in 1 mL of YES liquid medium, centrifuged at 1,000 × g, and the supernatant was discarded. The cells were then resuspended in 40 µL of YES medium and were spread onto YES agar plates. After a one-day incubation at 30°C, the colonies were replica-plated onto YES agar supplemented with the appropriate antibiotics and were incubated for two days. Single antibiotic-resistant colonies were then isolated by streaking them onto fresh YES + antibiotic plates and were incubated at 30°C for two days. The selected colonies were then transferred to YES plates without antibiotics for further expansion. Genomic DNA was extracted from these cultures, and the successful transformants were confirmed by PCR genotyping.

**5.8. Genomic DNA Extraction and PCR Validation of Transformants**

Colonies of approximately 5 mm × 5 mm in size were scraped from the YES agar plates and were resuspended in 100 µL of freshly prepared isolation buffer (250 mM LiAc, 1% SDS). The suspension was then incubated at 70–80°C for 5–10 minutes to lyse the cells. Subsequently, 300 µL of 100% ethanol was added, and the mixture was vortexed thoroughly. The samples were then centrifuged at 15,000 × g for 3 minutes at room temperature, and the pellet was washed with 70% ethanol. After discarding the supernatant, the pellet was resuspended in 100 µL of sterile water or TE buffer and was centrifuged again at 15,000 × g for 30 seconds. The resulting supernatant, which contained the yeast genomic DNA, was used directly for PCR analysis (1 µL per reaction).

The genomic DNA was amplified using the 2× PCR Super Mix (TRANSGENE, AS221). Each 10 µL PCR reaction contained 1 µL of genomic DNA, 0.2 µL of each primer (10 µM), 3.6 µL of nuclease-free water, and 5 µL of the 2× PCR Mix. The PCR products were then analyzed via agarose gel electrophoresis on a 1% gel to detect the target fragments.

For the overexpression strains, the genotyping forward primer was designed to be within the coding sequence (CDS) of the target gene, and the reverse primer (Table

S1R) was located on the *mCherry* sequence. The overexpression plasmid served as the positive control, while the genomic DNA from the wild-type strain was used as the negative control. Colonies that yielded PCR products with the same size as the positive control were successfully transformed overexpression strains.

For the knockout strains, the genotyping forward primer (Table S1R) was designed to be in the 5′ untranslated region (5′ UTR) of the target gene, and the reverse primer was located within the gene's CDS. The knockout plasmid was used as the positive control, and the wild-type genomic DNA served as the negative control. Colonies that showed no PCR band (matching the knockout control) were identified as successfully transformed knockout strains. In all cases, the sizes of the PCR products were compared against a DNA ladder to confirm their consistency with the expected fragment size. The identified positive transformants were picked using sterile pipette tips and were resuspended in 1 mL of YES liquid medium. The suspension was then transferred to a cryovial, mixed with 1 mL of sterilized 50% glycerol, and stored at –80°C.

### 6. Analysis of Correlation and Cellular Morphology in Rescued and Evolved Strains

Based on the identification of several genes that rescue the fusion rates of *prm1Δ* cells, an investigation was conducted to determine whether the increased fusion rate in the evolved strains occurs through multiple pathways or a single, unified pathway. To this end, the correlation of these rescue genes in terms of their ability to rescue *prm1Δ* cells was analyzed, along with potential interactions between these genes, the impact of these rescue strains on the cell cycle and cell morphology, and their involvement in the timing of meiotic progression.

#### 6.1. Identification of Interactions Between Rescue Genes

***Yeast-One-Hybrid Assay for DNA-Protein Interaction Verification*[23]**

As one of the identified rescue genes is a DNA-binding transcription factor (Ste11), a yeast one-hybrid assay was chosen for the preliminary confirmation of the DNA-protein interactions, as this method allows for rapid observation. The results of the Y1H experiment are shown in Supplementary Figure S10C. First, the yeast one-hybrid interaction verification kit was purchased from Coolaber Bioscience[23], which contains the necessary plasmids, strains, and media for the experiment. Next, based on the predictions from PomBase, Sms1 promoters of 1500 bp, 1000 bp, 642 bp, and 500 bp were selected for primer design and PCR amplification. The amplified fragments were then ligated into the *pAbAi* plasmid, while the CDS of Ste11 was ligated into the *pGADT7* plasmid.

***Transformation of pBait-AbAi into Y1HGold***

The *pAbAi* plasmid is integrated into the yeast chromosome via homologous recombination and thus exists within the yeast cell. Prior to transformation, the circular plasmid must be linearized by restriction enzyme digestion. The linearized *pAbAi* plasmid is then introduced into the Y1HGold yeast strain. The transformed recombinants (bait-specific AbA reporter strains) can grow on SD/-Ura medium.

***Linearization of the pBait-AbAi Plasmid***

After sequencing the *pBait-AbAi*, *p53-AbAi*, and Mutant Bait *pAbAi* plasmids, the *E. coli* cultures were expanded, and the plasmids were extracted. The plasmids were then linearized using the *BstB*I restriction enzyme (NEB). The digestion was carried out at 65°C for 2 hours, and the digestion products were run on a 1% agarose gel to verify the complete digestion of the vector, after which the fragments were purified and recovered.

***Transformation of pBait-AbAi into Y1HGold***

100 µL of ice-thawed Y1HGold competent cells (catalog number: CC308) were taken, and the pre-cooled linearized *pAbAi* plasmid (2-5 µg), Carrier DNA (10 µL, which had been heat-shocked at 95-100°C for 5 minutes, followed by a quick ice bath, repeated once), and 500 µL of PEG/LiAc solution were sequentially added. The mixture was mixed by pipetting up and down several times and was incubated in a 30°C water bath for 30 minutes (with mixing 6-8 times at the 15-minute mark). The tube was then placed in a 42°C water bath for 15 minutes (with mixing 6-8 times at the 7.5-minute mark). The mixture was then centrifuged at 10,000 rpm for 30 seconds, and the supernatant

was discarded. The pellet was resuspended in 400 μL of ddH2O, centrifuged again for 30 seconds, and the supernatant was discarded. The pellet was then resuspended in 50 μL of ddH2O, and 100 μL was plated onto SD/-Ura plates, which were then incubated at 30°C for 3-5 days. (Simultaneously, the lyophilized Y1HGold positive and negative control strains were rehydrated and activated by streaking them on SD plates). The bait strains were then identified by genotyping.

***Screening for the Optimal Working Concentration of AbA for Bait Yeast Strains***

The bait yeast strains exhibit very low basal AbA expression in the absence of the Prey vector. The optimal working concentration of AbA varies for different bait fragments. Therefore, the optimal AbA concentration for each bait yeast strain needs to be screened, with the following steps being taken:

After successful transformation verification, fresh single colonies (2-3 mm) were picked from each sample and were resuspended in 1 mL of 0.9% NaCl solution. The $OD_{600}$ was adjusted to 0.2 (alternatively, the cells were grown in SD-Ura liquid medium until the $OD_{600}$ reached 0.2). The resuspended culture was then serially diluted in 0.9% NaCl solution by 10-fold, 100-fold, and 1000-fold (i.e., to $OD_{600}$ values of 0.2, 0.02, 0.002, and 0.0002). 10 μL of each dilution was then spotted onto SD/-Ura and SD/-Ura plates with varying concentrations of AbA (100 ng/mL, 200 ng/mL, 300 ng/mL, 500 ng/mL, 800 ng/mL, and 1000 ng/mL). The plates were incubated at 30°C for 2-3 days, and the growth of the bait yeast on the plates with the different AbA concentrations was observed to determine the optimal working concentration of AbA (e.g., for the Y1HGold[p53-AbAi] bait strain, at an $OD_{600}$ of 0.002, no growth was observed on the SD/-Ura plates with AbA at a concentration of 200 ng/mL). (Note: The optimal AbA working concentration is determined by the lowest concentration at which there is a complete absence of yeast colonies on the plates with the different AbA concentrations (100 ng/mL, 200 ng/mL, 300 ng/mL, 500 ng/mL, 800 ng/mL, and 1000 ng/mL). This corresponds to the best inhibitory concentration, the minimum inhibitory concentration, the background expression concentration, or the self-activation concentration. In general, the AbA concentration should not exceed 1000 ng/mL).

***Preparation of Competent Y1HGold[pBait] Cells and Prey Transformation***

Competent cells of the successfully transformed Y1HGold[pBait-AbAi] strain were prepared using the TE-LiAc transformation method. Then, 2-4 µg of the pre-cooled target plasmid and 10 µL of single-stranded DNA (ssDNA) were sequentially added. The mixture was pipetted up and down to mix thoroughly, and then 100 µL of the yeast competent cells (prepared as described above) was added to the tube and gently mixed. 500 µL of PEG/LiAc solution was then added, and the mixture was mixed by inverting the tube. The mixture was incubated at 30°C for 3 hours, with the tube being gently inverted every 30 minutes. After the incubation, 20 µL of DMSO was added, and the mixture was mixed by inverting the tube. The tube was then incubated at 42°C in a water bath for 15 minutes, with the tube being inverted every 5 minutes. The cells were then centrifuged at 12,000 rpm for 15 seconds, and the supernatant was discarded. 1 mL of YPDA medium was added to the pellet, and the mixture was incubated at 30°C with shaking at 250-270 rpm for 90 minutes. The mixture was then centrifuged at 12,000 rpm for 15 seconds, the supernatant was discarded, the pellet was resuspended in 50 µL of ddH2O, and the cells were plated on SD/-Leu medium. The inverted plates were then incubated at 30°C for 3-5 days.

***Interaction Validation***

After successful transformation, fresh single colonies (2-3 mm) were picked from each sample and were resuspended in 1 mL of 0.9% NaCl solution. The $OD_{600}$ was adjusted to 0.2 (alternatively, the cells were cultured in SD-Leu liquid medium until the $OD_{600}$ reached 0.2). The culture was then sequentially diluted in 0.9% NaCl solution by 10-fold, 100-fold, and 1000-fold (i.e., to $OD_{600}$ values of 0.2, 0.02, 0.002, and 0.0002). In the order of the experimental group followed by the control group, 10 µL of each dilution was spotted onto the corresponding SD/-Leu plates with AbA*. The plates were then incubated at 30°C for 2-3 days, and the growth of the recombinant yeast on the plates with the corresponding self-activation AbA concentrations was observed to determine whether an interaction had occurred.

***Analysis of Interaction Verification***

From the screening for the optimal working concentration of AbA for the bait yeast strains, it was found that the self-activation concentration of Y1HGold[p53-AbAi] is 200 ng/mL. As shown in Figure S10C, on the SD/-Leu plates, the control groups Y1HGold[p53-AbAi+pGADT7-p53] and Y1HGold[p53-AbAi+pGADT7-REC2] exhibited similar growth. However, on the SD/-Leu plates with AbA (200 ng/mL), Y1HGold[p53-AbAi+pGADT7-p53] showed significantly better growth compared to Y1HGold[p53-AbAi+pGADT7-REC2], which indicates that p53-AbAi interacts with pGADT7-p53. Similarly, PSms1-AbAi and pGADT7-Ste11 also interact. (Note: The interaction between GAL4 AD-p53 and the p53 binding sequence (cis-acting element) induces the expression of the AbA resistance gene AUR1-C, which allows Y1HGold[p53-AbAi+pGADT7-p53] to grow on SD/-Leu plates with AbA at a concentration of 1000 ng/mL).

##### 6.2. Cell Cycle Analysis[21]

Given that the rescue strains displayed a significantly faster mating phenotype during the initial assays, it was hypothesized that this could be due to altered cell cycle kinetics. To test this possibility, the cell cycle profiles of all the rescue strains were analyzed by measuring their DNA content with flow cytometry[21]. First, fresh single colonies were picked from the streak plates and were inoculated into YES liquid medium. The cultures were then grown overnight to the log-phase, and 1-10 × $10^6$ fresh yeast cells were harvested. The cells were then fixed by resuspending the pellet in 200 µL of 70% ethanol and were incubated for 1 hour at room temperature. The cells were then centrifuged, and the supernatant was aspirated to remove the ethanol. The pellet was then resuspended in 200 µL of RNase A solution and was incubated at 37°C for at least 3 hours and up to overnight. Subsequently, the cells were centrifuged, and the supernatant was aspirated to remove the RNase A. The pellet was then resuspended in 200 µL of proteinase K solution and was incubated at 55°C for at least 3 hours and up to overnight. The cells were then centrifuged, and the supernatant was aspirated to remove the proteinase K. The pellet was then resuspended in 200 µL of 0.05 M sodium citrate solution and was sonicated using a SonicMan sonicator (Matrical Bioscience) set at 60% power, for 10 seconds, and 2 cycles. PI (Propidium Iodide, 1 mg/mL) staining solution was then added at a 1:500 ratio, and the mixture was incubated at room temperature for 15 minutes, after which the cell DNA content was analyzed using a flow cytometer.

The DNA content experimental data were analyzed using FlowJo software (v10.6.2). First, the x-axis was set to FSC-A, and the y-axis was set to SSC-A. The 'T' icon was clicked, and a linear axis was selected, after which the range was adjusted using the 'Custom Range' tab. The cells were then gated, and the cell gate was double-clicked to proceed. Next, the appropriate fluorescence channel was selected. It was ensured that the x-axis represented the area, and the y-axis represented the height. The 'T' icon was clicked again, and both axes were set to a logarithmic scale. The range was then adjusted using the 'Custom Range' tab, and the appropriate population was gated to exclude any doublets. The gate was then double-clicked, the y-axis was changed to a histogram, and the G1 and G2 phases were gated. Returning to the main analysis window, 'Cells' → 'Single Cells' → 'G1' → 'G2' were selected from the last analyzed sample, and all four were dragged into the 'All Samples' section at the top. Then, 'Cells' was opened, and the gates were adjusted for each sample individually. For 'Single Cells' and the subsequent populations (G1/G2), the gates were adjusted across all samples accordingly. To visualize the data, the 'Layout Editor' was used to display or overlay the samples as required by the experiment. The G1/G2 populations from the sample analysis window were dragged into the 'Table Editor'—not from the main G1/G2 gate, but specifically from the sample data. 'Mode' was selected to determine the peaks for G1 and G2, ensuring that the appropriate fluorescence channels were selected. Additionally, 'Frequency of Parent' was chosen to calculate the percentage of the G1 and G2 populations. The results of this experiment are shown in Figures 5G and 5H, and in Supplementary Table S1L.

##### 6.3. Cell Biology Analysis

###### *Fluorescent Tagging Experiment*

The CDS and 3' UTR of the target gene were utilized as homologous arms, and a fluorescent protein and a resistance selection marker were inserted between the CDS and the 3' UTR of the target gene. First, the pGL94 (*pFA6a-Fus1 5'UTR-hphMX-Fus1 3'UTR*) plasmid was digested with *Sal*I, *Asc*I, *Sac*I, and *EcoR*V, and the two large fragments were used as the plasmid backbone. Then, Infusion cloning was used to ligate the CDS of the target gene, a fluorescent tag, a resistance selection marker, and the 3' UTR of the target gene into the backbone, thereby constructing a plasmid that labels the target gene with a specific fluorescent protein. Finally, lithium acetate transformation was used to insert the CDS and the 3' UTR of the target gene, which served as homologous arms, into the yeast genome. After screening for positive clones, they were stored at -80°C for future use. The tagged strains that were used are listed in Supplementary Table S1Q.

***Measurement of Fusion Neck Length***

The overnight images of the rescue strains and the control strains under nitrogen starvation were observed, and ImageJ (v1.53, NIH) was used to measure the fusion neck length of the cells in the early stages of fusion, as observed in the video. First, at least thirty mating pairs were measured for each sample. The 'Straight' tool was used to draw the fusion neck, and it was added to the ROI Manager. Once 30 pairs of cells had been recorded, 'Measure' was clicked to obtain the length measurements. The measurement results are presented in Figure 6E-F and in Supplementary Table S1N.

***Measurement of Cell Size***

Cellpose was used to segment each cell and to output the area and length of the cells[22]. First, ANACONDA was downloaded, 'Environments' was clicked, and then 'Create' was clicked. In the 'Name' field, 'Cellpose' was entered, 'Python' was selected in the 'Package' field, and 'Create' was clicked. Next, the anaconda prompt was located and opened on the computer, the commands were entered, and finally, Cellpose was launched. After opening Cellpose, the images to be analyzed were imported (the required images were pre-exported as PNG, TIFF, or JPG formats using ImageJ (v1.53, NIH) and were saved in a folder). The diameter parameter was then adjusted, as the typical diameter of pombe cells ranges from 65 to 75. Once the diameter parameter had been adjusted, 'Run Cyto3' was clicked to segment the cells. Then, 'File' was clicked, 'File-Save outlines as .zip archive of ROI files for ImageJ' was selected, and the ROI data were imported into ImageJ. The scale was then set according to the microscope parameters, and 'Measure' was clicked to output the cell area parameters. The measurement results are presented in Figure 6G-H and in Supplementary Table S1O.

***Analysis of Mating Cell Composition***

All the image analysis was performed on the live-cell imaging data. Two distinct parameters were quantified from the videos of the *prm1Δ* mating process. First, to determine the mating behavior, the time-lapse videos were visually inspected frame by frame. A cell was scored as "attempting to mate with multiple partners" if it first extended a shmoo projection and made physical contact with one partner, and then subsequently retracted that projection and initiated a new mating attempt with a different partner. If a cell fails to mate with the first partner it encounters and subsequently seeks another cell for mating, this event is recorded as a "second-partner" pair, and the percentages were calculated using the following equation (Figure 6A-B, and Supplementary Table S1M):

$$\% \text{ mating cell composition} = \frac{second\ partner}{first\ partner + second\ partner} \times 100$$

Second, the number of fluorescent foci per cell was also quantified using ImageJ (v1.53, NIH). A random sampling approach was used to quantify the foci. The number of discrete foci within each random ROI was then counted at specific time points using the multi-point tool (Figure 6C-D, and Supplementary Table S1M).

For all statistical analyses of the cell biological data, the Student's t-test was used. The data are presented as mean ± SD (*$p$< 0.05, **p< 0.01, ***p< 0.001, ****p< 0.0001).

#### 6.4. Measurement of Growth Rate and Fusion Efficiency for All Rescue Strains

During the study, it was observed that the rescue strains exhibited slow growth rates. To comprehensively analyze the relationship between fusion efficiency and growth rate in the rescue strains, experiments were conducted to measure both the fusion efficiency and the growth rate. The experimental procedures were performed as described in sections 5.4.2 and 5.5. The experimental results are shown in Figure 7G, with the complete supporting data provided in Supplementary Tables S1I and S1J.
